# Time-resolved functional genomics using deep learning reveals a global hierarchical control of autophagy

**DOI:** 10.1101/2024.04.06.588104

**Authors:** Nathalia Chica, Aram N. Andersen, Sara Orellana-Muñoz, Ignacio Garcia, Aurélie Nguéa P, Sigve Nakken, Pilar Ayuda-Durán, Linda Håkensbakken, Sebastian W. Schultz, Eline Rødningen, Christopher D. Putnam, Manuela Zucknick, Tor Erik Rusten, Jorrit M. Enserink

## Abstract

Recycling of cellular components through autophagy maintains homeostasis in dynamic nutrient environments, and its dysregulation is linked to several human disorders. Although extensive research has characterized the core mechanisms of autophagy, limited insight into its systems-wide dynamic control has hampered predictive modeling and effective *in vivo* manipulation. In this study, we mapped the genetic network that controls both the dynamic activation and inactivation of autophagy during nitrogen changes, using a combination of time-resolved high-content imaging, deep learning, and latent feature analysis. This approach generated a comprehensive genome-wide profiling repository, termed AutoDRY, categorizing 5919 mutants based on their nutrient response kinetics and differential contributions to autophagosome formation and clearance. Integrating these profiles with functional and genetic network data unveiled a hierarchical and multi-layered control of autophagy, identifying new regulatory aspects of the core machinery and established nutrient-sensing pathways. By leveraging multi-omics resources and explainable machine learning to predict genetic perturbation effects and infer new regulatory mechanisms, we identified the retrograde pathway as a pivotal, time-varying autophagy modulator through transcriptional tuning of core genes. By charting the systems-wide dynamical control of autophagy, we have laid the groundwork for connecting the complexity of genome-wide influences with specific core mechanisms. This represents a significant advancement in studying complex genetic phenotypes, guides functional genomics of dynamic cellular processes in any organism, and provides a powerful starting point for hypothesis-based research on autophagy.

## Introduction

Living organisms dynamically tune their metabolism to changes in nutrient availability. Macroautophagy, hereafter autophagy, sustains cellular homeostasis by adaptively degrading and recycling cellular components in response to nutrient scarcity, as well as other environmental cues, and its fine-tuning is crucial for a healthy lifespan^1,2^. This is evidenced by tight multi-layered control of the autophagy core machinery, and the association of several autophagy-related genes (ATGs) with human disorders^2–4^. Characterizing autophagy execution has been the subject of intense research and several regulatory pathways have been identified^5–9^. However, a generalized understanding of how living organisms dynamically control autophagic activity in the context of a broader genetic landscape is lacking. Along these lines, pharmacological manipulation of autophagy remains rudimentary and involves either complete inhibition of the process or strong activation through the use of TORC1 inhibitors, which have substantial side effects on growth and cell proliferation^10^. A systems-level understanding of the tuning of autophagic activity may help develop predictive models and support the development of effective treatments for autophagy-related conditions.

While emerging studies utilizing multi-omics approaches, mathematical modeling and network analyses have begun to elucidate the complex landscape of potential autophagy regulators, systematic genome-wide data on causal influences on the autophagic nutrient response are limited^11–15^. To date, the study of autophagy has largely been based on gain- or loss-of-function discoveries from autophagic flux assays, which measure the cumulative degradation or re-distribution of cargo over a fixed time window^16,17^. This approach has been successful in elucidating the central mechanisms orchestrating autophagy, as well as proximal upstream switches acting in response to starvation. Genome-wide screens of autophagy-related mutants in yeast have identified only a limited number of new genes because they screened solely for loss in viability during starvation^18^ or for changes in autophagy flux at a single time point^19^. However, these assays do not fully capture the dynamic nature of autophagy regulation and may overlook important genomic factors influencing activation thresholds, sensitivity to nutrient shifts, and more distant homeostatic tuning^17,20–23^. These shortcomings also extend to a limited focus on the deactivation of autophagy and return to homeostasis upon nutrient replenishment. Moreover, systematic information on the regulation of different stages of autophagy, such as the formation and clearance of autophagosomes, is lacking. To address these limitations, more sensitive real-time recordings of changes in autophagy during nutrient depletion and repletion are needed, along with a systematic analysis of cellular features associated with distinct stages of the process.

In this study, we performed a functional analysis of the genome-wide regulation of autophagy dynamics in *Saccharomyces cerevisiae*. By utilizing time-resolved high-content imaging combined with deep learning, we quantified the nutrient response kinetics and differential influences on autophagy execution in 5919 yeast gene mutants (90% of the yeast genome) with high precision. By assembling quantitative regulatory profiles, and integrating these with genome-wide resources that exist in yeast, we constructed a comprehensive map of the autophagy network response to shifts in nitrogen supply. Our approach reveals previously unexplored gene and systems-level regulators of autophagy dynamics, including phase-dependent control of autophagic flux, and enables genome-scale predictive modeling of autophagy.

## Results

### Genome-wide influences on autophagy dynamics

To map the genome-wide control of autophagy, we used an arrayed library of 4760 deletion mutants and 1159 Decreased Abundance by mRNA Perturbation (DAmP) mutants expressing the autophagosome marker mNeonGreen-Atg8 and the vacuole-resident protease Pep4-mCherry. Mutants were grown to log-phase in nitrogen-containing medium (+N), then imaged every hour for 12 hours during nitrogen starvation (-N) and for 7 hours following nitrogen replenishment (+N) using high-content fluorescence microscopy (Fig. 1A). Each array included triplicate wells of wild-type (WT) cells and autophagy induction-deficient *atg1*Δ mutant cells as controls. We also included triplicates of *vam6*Δ mutant cells, which exhibit defects in Gtr1-dependent TORC1 activation and in homotypic fusion with the vacuole, resulting in an abundance of uncleared autophagosomes^24–26^.

**Figure 1.**
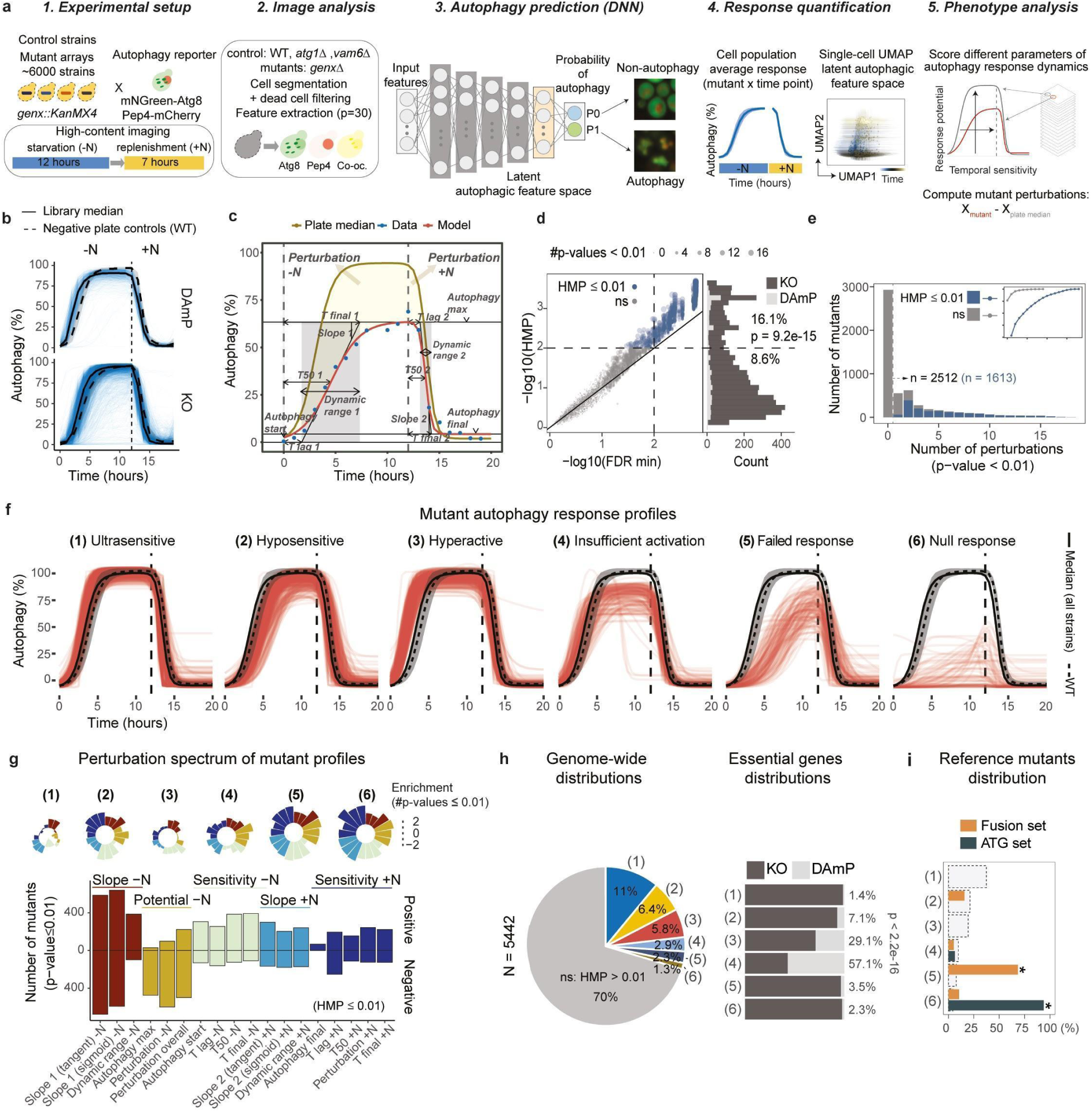
Deep learning-based profiling of autophagy nutritional response kinetics. **a,** Study workflow overview: 1. An autophagic reporter integration into a yeast gene deletion library; hourly image capture during starvation (-N) and nitrogen replenishment (+N). 2. Cell segmentation and extraction of 30 image features (p). 3. Neural network training to predict autophagy probability (P0 non-autophagy; P1 autophagy) from control data. 4. DNN output and latent space are used to predict autophagy response and states in mutants. 5. Computation of mutant perturbations as differentials from the plate median. **b,** Predicted autophagy response as a percentage of cells in autophagy for KO and DAmP collection strains. Median (solid line), negative plate control means (dashed line) and standard deviation (ribbon). **c,** Double sigmoidal model fitting was used to measure kinetic perturbations and interpolate the autophagy responses. **d,** HMP correlates with the minimum FDR over the set of hypothesis tests performed and is more sensitive than FDR for mutants with multiple significant perturbations. Distributions and percentages indicate the proportions of DAmP collection mutants; Fisher’s exact test. **e,** Distribution of mutants with significant parameter perturbations (p-value ≤0.01). Inset indicating the cumulative probability. Significant mutants (HMP ≤0.01, blue) have multiple significant perturbations, with 50% having four or more (see inset). **f,** Autophagy response dynamics for the six profiles. **g,** Distribution of significant parameter perturbations among significant mutants. The color coding indicates groups of related parameters based on clustering (see Suppl. Fig. *S4D*). The bar graph (bottom) represents the number of positive or negative significant perturbations per parameter. The radial bar charts (top) indicate log2-fold enrichment of significant perturbations (p-value ≤0.01) in the respective profile over the entire genome. **h,** Genome-wide proportions of mutant profiles (*left*) and representation of DAmP collection mutants within each class (*right*); chi-square test. **i**, Distribution of reference mutants across response profiles; chi-square with Bonferroni corrected post-hoc test,*p ≤0.001.

To robustly quantify the autophagy responses across the genome-wide library with minimal technical confounders and observation bias, we first assembled a large training dataset of extracted image features consisting of 1.34M automatically annotated WT and *atg1*Δ cells from 70 independent experiments. We then applied a deep learning approach to classify autophagic activity (Fig. 1A and S1A-K). The image features included 31 single-cell parameters summarizing the distribution and intensity of the two primary markers, as well as a third channel representing their pixel-wise co-occurrence. The output labels of the training data were determined by experimental control conditions where autophagy was fully activated (WT cells for 7 hours in -N) or completely inactive (either *atg1*Δ across all time points or WT cells for 4 hours in +N) in the cell populations, thus avoiding the limitations of human annotation. Subsequently, fully connected deep neural networks (DNNs) were trained to predict the activation state per cell based on the image features.

A hyperparameter grid search across 3096 model architectures demonstrated that multiple DNNs could predict the rates of autophagy activation with high accuracy (>98%) and reproducibility across experiments (Fig. S1B-F). Moreover, UMAP embedding of latent autophagic features represented in the DNNs revealed a continuous change in WT cells and intermediary phenotypes (*vam6*Δ), in which cells transition between distinct autophagic states with low levels of non-autophagic noise (variance in the *atg1*Δ embedding; Fig. S1G-I). Out of the top 30 model architectures, the DNN with the most consistent performance across several evaluation metrics (‘Model 22’) was selected to predict autophagy in the genome-wide screen (Fig. S1E-H). To ensure robust analysis of latent space features and avoid systematic measurement bias due to differences in activation functions, we compared the embeddings of two distinct DNNs: ‘Model 22’ and ‘Model 30’ (Fig. S1H-I). This enabled reliable profiling of nutritional responses across plates, resulting in 5678 unique starvation responses and 5442 replenishment responses after removing mutants with low-confidence predictions and low cell counts (Fig. S2A-E).

For most phenotypes, both the activation and inactivation phases in response to nutrient shifts displayed sigmoidal kinetics, with the inactivation phase exhibiting a much steeper decline (Fig. 1B and S2A). Using a double-sigmoidal model, we extracted 15 parameters that captured different aspects of the response kinetics (Fig. 1C and S3A-B)^27^. These included (1) the activation and inactivation rates in each treatment phase measured by the slopes or width of the dynamic ranges; (2) the sensitivity to nutrient shifts measured by the response times (‘T lag’, ‘T50’, and ‘T final’); and (3) the autophagy activation potentials at the start, maximum, and final points of the time course. Mutant perturbations were subsequently assessed by calculating the difference in each parameter from the corresponding array medians (Fig. 1A), along with the average autophagy perturbation (%) computed as the differential integral under the curve for each phase separately (denoted ‘-N’ and ‘+N’) or the entire time course (denoted ‘overall’; Fig. 1C and S3C-E).

To assess the statistical significance of individual parameter perturbations, we performed multiple hypothesis tests using an expected error model (H_0_) derived from replicate WT and negative control measurements within each plate (Fig. S3C-E and S4A). The global statistical significance of each mutant was scored by combining its test results using a harmonic mean p-value (HMP) weighted by the genomic H_0_-rejection rate under a 5% Benjamini-Hochberg (BH) rejection threshold for each parameter (Fig. 1D and S4B)^28^. A 1% HMP cutoff identified 1613 mutants with significant perturbations in multiple parameters, and a relative overrepresentation of essential genes (DAmP mutants; Fig. 1D, E and S4C; Data File S2). Grouping the spectrum of parameters into five distinct classes (Fig. S4D) revealed that mutants with changes in starvation responses, particularly in the slopes, were more numerous and varied compared to those with changes in replenishment responses, suggesting a greater regulatory complexity in the activation phase of autophagy (Fig. 1F, G and S4A).

Clustering the responses of significant mutants revealed six distinct autophagy perturbation profiles (Fig. 1F and S4E), which were categorized based on the type of kinetic changes that characterized each cluster (Fig. 1G)^23^. For instance, three major groups (‘*ultrasensitive*’, ‘*hyposensitive*’, and ‘*hyperactive*’) exhibited changes in temporal sensitivity and activation rates (Fig. 1G, H). Hyposensitive mutants displayed delayed responses to nutrient shifts, while ultrasensitive and hyperactive mutants displayed increased activation and inactivation rates (slopes), with hyperactive mutants further having elevated basal autophagy (at T0) and higher sensitivity to -N. Three minor groups (‘*insufficient activation*’, ‘*failed response*’, and ‘*null response*’) exhibited impaired response potential at different degrees of severity (Fig. 1G, H). The null responses displayed minimal autophagy activation across the entire time course, while failed responses exhibited severe deficiencies in activation. Lastly, mutants with insufficient activation responded more normally to nutrient shifts, but were unable to fully activate autophagy in -N. These mutants also displayed slight elevation in basal activity. Analyzing the composition of each cluster revealed that essential genes tended to be disproportionately represented among the hyperactive and insufficient activation profiles (Fig. 1G), reflecting either dynamic differences between the DAmP and KO libraries (Fig. 1B) or suggesting a potential relation between basal autophagy, starvation response potential, and growth control (Fig. S4A).

To inspect the profile distribution of well-characterized autophagy genes, we defined two ‘gold-standard’ reference sets of autophagy related genes through literature curation and manual inspection of images (see Fig.S5A, Methods 4.1 and Table S1). A set of ATG genes essential for the induction of autophagy (‘ATG set’, n=16), was significantly overrepresented among null responses, and a set comprising genes involved in the maturation or clearance of autophagosomes (‘Fusion set’, n=19), was significantly overrepresented among failed responses. These distributions indicate that the different response profiles meaningfully reflect the severity of known autophagy deficiencies.

### Accuracy and robustness of autophagy perturbation profiling

To evaluate the capacity of the perturbation statistics to identify true autophagy-associated genes, we analyzed the precision-recall curves (PRC) for the two autophagy reference sets, comparing them with a ‘negative control set’ consisting of deletions of dubious ORFs that did not overlap with verified genes (n=23), or with random samples from the genome-wide distribution. For the combined ‘ATG’ and ‘Fusion’ reference sets, the HMP yielded the best precision-recall on average, with an area under the curve (PRC-AUC) > 0.97 (Fig. S5B-C). The individual kinetic parameter p-values yielded PRC-AUCs that were in the 0.7-0.95 range when compared with genome-wide random samples, and in the 0.8-1 range when compared with the negative control set (Fig. S5D-F). Interestingly, while the ‘ATG’ and ‘Fusion’ mutants exhibited severe autophagic deficiencies in starvation, many ‘Fusion’ mutants with delays in autophagosome clearance displayed slow deactivation in response to nitrogen replenishment (see T50 in +N; Fig. S5A, F, G).

We also evaluated the reproducibility of the measured perturbation phenotypes. Repeating the starvation protocol for the top mutants (selected based on the HMP; Fig. S6) revealed that the dynamics of stronger autophagy perturbation phenotypes were more likely to replicate across experiments (Fig. S6G). Additionally, by assessing the representation of autophagy phenotypes among interacting genes in various yeast network databases based on physical and genetic interactions (see Methods 7.4.1), we observed a genome-wide concordance between a gene’s autophagy perturbation statistic and enrichment of phenotypes among its nearest neighbors (Fig. S6B-E)^29–31^. Discordant outliers within this data, along with ‘autophagy’ GO-annotated mutants yielding weak perturbations, were used to predict potential false positives (FPs) or negatives (FNs) that were subsequently retested (Fig. S6D-G). This analysis identified only 56 potential FPs and 109 potential FNs. More than half of predicted FPs produced replicable autophagy phenotypes, representing potentially novel autophagy-related genes, while only 27 mutants represented outliers that regressed in perturbation magnitude, resulting in weak or no correlation between experiments (Fig. S6F, G). Among the predicted FNs, 42 mutants were significant under a more liberal threshold (HMP ≤ 0.05) and displayed a discrete but replicable autophagy perturbation phenotype (Fig. S6H). These mutants were also supported by stronger network evidence (Fig. S6I-J) compared to non-significant mutants (HMP > 0.05), which were more likely to have been selected based on GO annotations (Fig. S6K). Further inspection revealed that many of these non-significant mutants represented genes with specialized roles, such as genes that are not essential for starvation-induced autophagy, including those involved in pexophagy (*ATG36*), the Cvt-pathway (*ATG20*, *ATG34*, *ATG19*), or mitophagy: *ATG32*, *ATG33*), as well as genes with weakly penetrant, but replicable phenotypes such as *VPS15*, *FAR11*, *PEP12* and *NPR3* (Fig. S6K).

Finally, to test the reliability of the genome-wide analysis, 33 strongly significant (HMP ≤ 0.001) deletion mutants, representing different dynamic profiles and biological functions, were reconstructed in triplicate and subjected to a set of validation experiments with variations in the growth protocol (Fig. S7A-B, see Methods 7.1-7.3). The reproducibility of phenotypes across independent clones and different protocols was similar to that of the same clones across all experiments, with a minor strain background effect and elevated noise in autophagy response caused by growing the cultures to stationary phase (Fig. S7C-G). Moreover, the time point-wise autophagy perturbations (Fig. S8A-D) and response kinetic parameters (Fig. S8E-F) showed strong correlations across mutants between the screens, with metrics for activation levels (Spearman correlation > 0.8), response times (Spearman correlation > 0.7), and slopes (Spearman correlation > 0.75) in -N being the most robust (Fig. S8F). These observations indicate that the genome-wide variability in mutant response kinetics is highly reproducible.

### Cellular processes that impact autophagy dynamics

To identify pathways affecting autophagy dynamics, we first performed gene-set enrichment analysis (GSEA) on each of the individual response statistics (parameter signed −log_10_ p-values) for sets defined by GO biological processes. Significantly enriched terms (at least one enrichment p-value < 0.005) were analyzed using principal component analysis (PCA) on the matrix of enrichment statistics (signed −log10 p-values) to identify which kinetic parameters captured the most functional variation. This approach allowed us to examine the GO term enrichment in a non-redundant manner by summarizing the functional variation along the primary principal components. Investigating the contributions of each PCA loading (representing a kinetic parameter) revealed that response potential and sensitivity to starvation aligned with the dominant direction of enriched GO biological processes (Fig. 2A), with terms related to macroautophagy and vacuolar activity yielding the strongest associations (Fig. 2B). In contrast, sensitivity to nitrogen replenishment aligned more with the second principal component (Fig. 2A), and was enriched for processes related to membrane trafficking and fusion (Fig. 2B). Interestingly, these GO terms were highly decorrelated from the direction capturing mechanisms controlling the response potential (Fig. 2B). Finally, perturbations in initial and final autophagy levels, which were the least correlated with the other parameters (Fig. S4D and 2A), were enriched for terms related to amino acid and nucleoside metabolism, nitrogen utilization, and mitochondrial activity (Fig. S9A).

**Figure 2.**
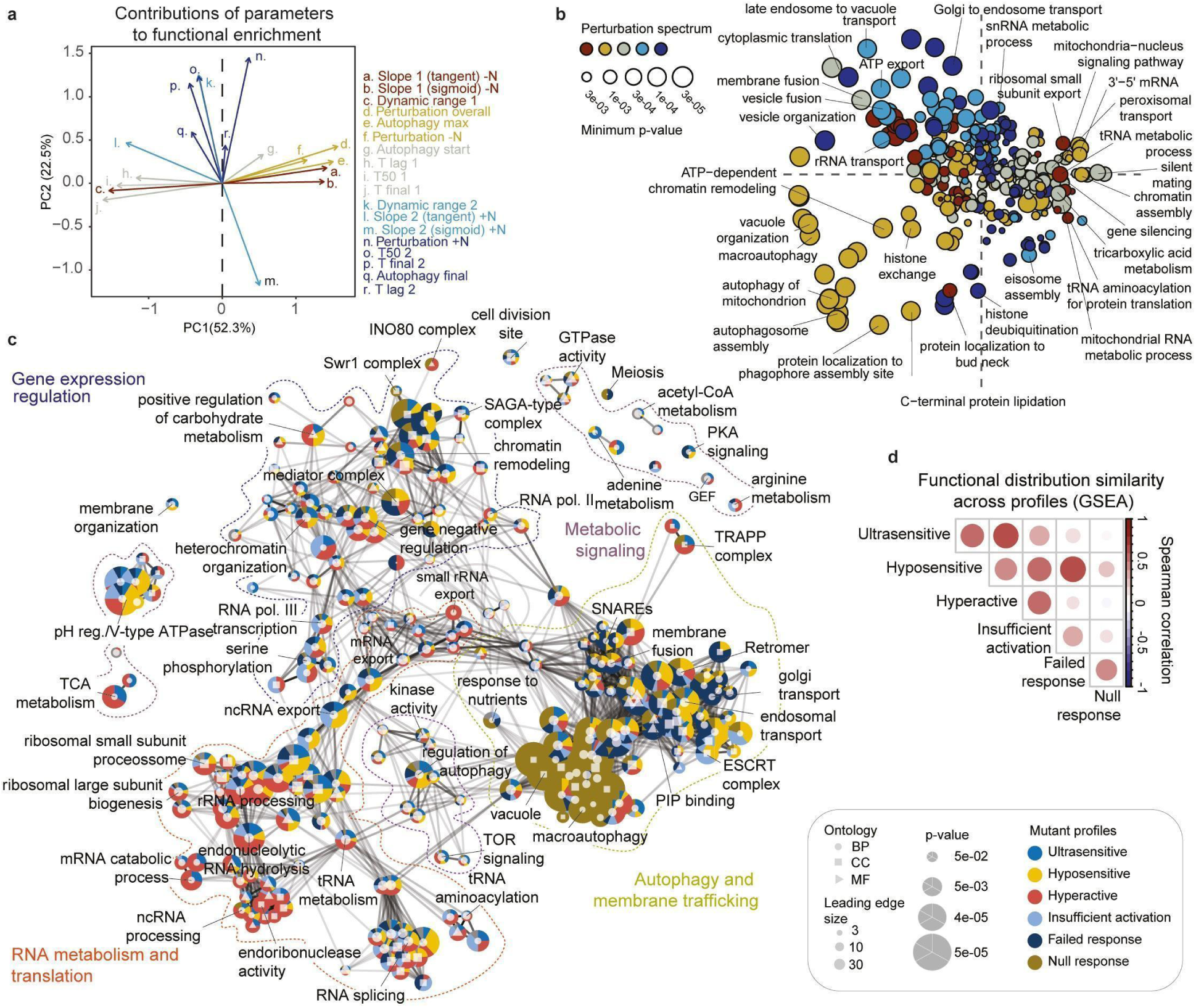
Kinetic perturbation profiling is highly sensitive and potentiates deeper annotation of regulatory modules. **a,** PCA loadings for kinetic parameters using a matrix of signed −log10 p-values from GSEA of GO-BP. Parameter class colors coding are the same as in Figure 1f**. b,** PCA of GO-BP terms from *a*. Colors indicate the perturbation parameter class (see Fig. 1f) with the most significant enrichment. **c,** Enrichment map based on Jaccard similarity of enriched GO terms from GSEA over the HMP. Edges are drawn for terms with 10% overlap. The pie charts represent the composition of perturbation profiles among enriched genes (leading edge). **d,** Similarity in functional distribution of perturbation profiles across enriched GO terms, assessed by Spearman correlation of GSEA-leading edge counts from *c*.

We then identified enriched GO terms by GSEA using the combined HMP statistic (Fig. 2C). An enrichment map provided an integrated overview of the primary functional categories influencing autophagy with three large, interconnected clusters representing autophagy and membrane trafficking; RNA metabolism and translation; and gene expression regulation. This map highlighted a graded distribution of perturbation profiles with homogenous enrichment of severe autophagy deficiencies (*null response* and *failed response*) close to autophagy-essential GO clusters, and less severe deficiencies in neighboring clusters (Fig. 2C). Quantifying the enrichment similarity between the different profiles revealed a gradual functional overlap between mutants causing partial loss-of-function phenotypes such as failed response, hyposensitive, and insufficient activation (Fig. 2D). Interestingly, the gain-of-function profiles *ultrasensitive* and *hyperactive* co-occurred more frequently with each other, as well as co-occurring more frequently with *hyposensitive* and *insufficient activation* profiles respectively, indicating consistency in regulatory phenotype pairing along the dimensions of sensitivity and response potential (Fig. 2D).

### Causal structure and network architecture of autophagy regulators

A phenotypic change in response to a genetic intervention does not necessarily imply that a gene is a direct regulator of the measured phenotype. We therefore more closely examined the organization of autophagy-influencing genes and their relation to the autophagy core machinery in functional networks. Previous work in yeast has shown that genome-wide networks can accurately capture the functional organization of the cell when constructed using similarities in genetic interactions, which are calculated using the Pearson Correlation Coefficients of interactions from Synthetic Genetic Array data (SGA-PCC), or when constructed using protein-protein interactions (PPIs)^29,32,33^. Using spatial analysis of functional enrichment (SAFE) for the different perturbation profiles, we found distinct genome-wide enrichment patterns for all the profiles except *ultrasensitive* in both genetic similarity and STRING-based PPI networks (Fig. 3A and S10A)^34^. It is noteworthy that the ‘ATG set’ did not form a distinct cluster in the genetic similarity network but did so in the PPI network, where its spatial enrichment overlapped with that of *null responses*, and it co-localized with processes related to membrane fusion (Fig. S10A).

**Figure 3.**
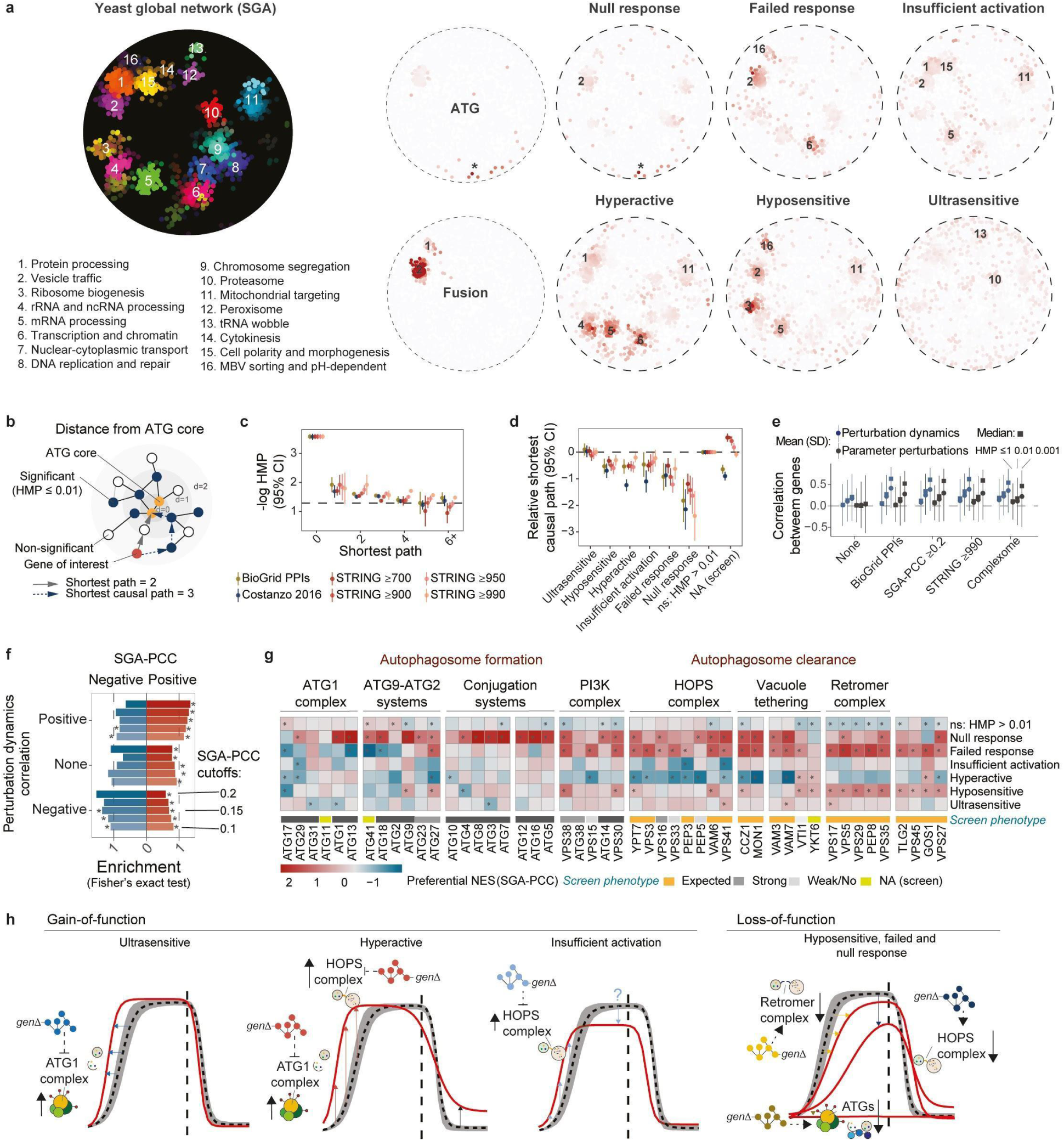
Genome-wide regulatory network architecture of autophagy-influencing genes. **a,** SAFE analysis of the global genetic similarity network. Reference map of bioprocesses (left) and of ATG and fusion set (middle) for interpreting the spatial enrichment of autophagy perturbation profiles (right); *ATG core enrichment. **b,** Quantification of shortest paths from the ATG core machinery. **c,** Average HMP (-log10) per shortest path distances in different functional interaction networks, computed as indicated in *b*). The dashed line indicates HMP = 0.05. **d,** Shortest causal pathlength per autophagy perturbation profile relative to the non-significant genes (HMP > 0.01). **e,** Correlations in temporal autophagy perturbations or sets of perturbation parameters (signed −log10 p-values) between genes that share network interactions or complexome association or lack any interaction altogether (none). Correlations were considered for different HMP cutoffs (1, 0.01 or 0.001). **f,** Enrichment of positive or negative genetic interaction similarities (SGA-PCCs) per correlation intervals of autophagy perturbations between pairs of strongly significant genes (HMP ≤ 0.001). *p-value ≤0.05 (Fisher’s exact test). **g,** Direction preferential enrichment (computed as the max-min difference between the GSEA normalized running-sum statistics, NES; see Suppl. Figure *S11c* and methods 5.4) of perturbation profiles along the SGA-PCC spectra for individual genes of the core autophagy machinery. *p-value ≤0.05 (permutation test). Screen phenotypes were manually categorized according to the expected or degree of perturbation. NA refers to gene mutants not available in the screen. **h,** Illustration depicting hypothetical regulatory entry points within the modules of the autophagy core machinery, derived from the significant enrichment of different perturbation profiles along SGA-PCC spectra in *g*.

Interestingly, more severe autophagy phenotypes were enriched closer to the ATG core sub-graph (‘ATG set’, including *ATG1* and *ATG8*). The proximity of autophagy-influencing genes to the ATG core in these networks was confirmed in two ways when analyzing the shortest path lengths (Fig. 3B). First, the HMP was more significant for genes that were closer to the ATG core than for those that were further away (Fig. 3C). Second, the average shortest path length for each autophagy perturbation profile, except *ultrasensitive*, was significantly closer to the ATG core than the average shortest path length for genes that were not in any perturbation profile (HMP > 0.01; Fig. S10B-F), suggesting that these phenotypes on average represent more direct influences on autophagy. This relationship became even stronger when pathlengths were calculated using the shortest causal path (requiring all intermediate nodes to have a significant influence on autophagy in response to a gene deletion, HMP ≤ 0.01; Fig. 3B, D). In contrast, the 11% of mutations in the *ultrasensitive* profile, which only affect response sensitivity, might involve more distant fine-tuning of the nutrient response, spanning through several intermediates as indicated by the observed path length.

Consistent with the clustering of autophagy-influencing genes in these networks, gene pairs with reported functional relations (BioGrid PPIs, SGA-PCC, and STRING)^29–31^ or complexome associations^35^ exhibited correlated autophagy perturbations (Fig. 3E and S11A-B), suggesting that the dynamic profiles were consistent with functional gene relations on a genome-wide scale. The signs of the pairwise SGA-PCC scores can distinguish between inhibitory and activating gene relations^36–38^. We examined whether these relations would also be consistent with the signs of autophagy correlations by assessing whether gene sets with different SGA-PCC cutoffs were enriched in the perturbation dynamics correlations using Fisher’s exact test. Indeed, divergences in correlations between strongly significant genes caused a shift in the representation of positive and negative SGA-PCC (Fig. 3F). Moreover, when testing for directional parametric enrichment of mutant profiles over the SGA-PCC of core autophagy genes by GSEA, *hyperactive* phenotypes (in contrast to loss-of-function phenotypes) were preferentially associated with negative SGA-PCCs for both ATG and fusion core genes, suggesting an inhibitory relation (Fig. S11C-D). Using this approach, we inferred differential regulation of the various sub-systems executing autophagy, which cannot be measured from their null or deficient individual deletion phenotypes (Fig. 3G and S11E). Here, *ultrasensitive* responses were negatively related to genes involved in the early stage of autophagosome formation, particularly Atg1, suggesting that this may be the main limiting factor for the starvation response rate. Interestingly, *failed response*, which was positively related to the fusion core genes, was negatively related to some genes involved in autophagy induction (Fig. 3H). This could indicate the existence of negative feedback circuits from stages involved in autophagosome clearance^39^, or pleiotropic functions of gene products involved in amino acid liberation and TORC1 re-activation (such as Vps41 and Vam6)^24,40^.

To explore the regulatory relations between specific autophagy-influencing genes, we mapped the autophagy perturbation phenotypes across curated complexome networks representing the three major regulatory cell layers: gene expression regulation; RNA metabolism and translation; and membrane dynamics and trafficking, which exhibited systems-wide influences on autophagy dynamics (Fig. S10G-J). This allowed us to observe the penetrance and distribution of phenotypes across well-characterized functional modules. For example, we observed clustering of multiple significant phenotypes, such as in the chromatin remodeling sub-module, and consistency in the direction of autophagy perturbation for regulatory sub-graphs (e.g., Ure2 and Gln3, Gal80 and Gal3, or the CURI complex)^41–43^. For RNA metabolism and translation in particular, we detected a systemic enrichment of stronger phenotypes along modules related to mRNA processing and translation decoding (e.g. binding of mRNA by the 40S small ribosomal subunit), where perturbations in mRNA maturation and binding resulted in a reduced autophagic response, while perturbations in mRNA decay caused the opposite phenotype.

### Dynamic latent-space analysis of autophagosome formation and clearance

We reasoned that the latent autophagic features generated by the DNN (Fig. 1A, see autophagy prediction) potentially provided more information about cellular states that could be captured by the binary classifier used to determine the autophagy response (Fig. S1I). We therefore projected these latent autophagic features onto two dimensions using UMAP (Fig. 4A-B and S12A-B), where information about autophagy was conserved together with cell-to-cell variation and dynamic data from the DNN latent space (Fig. S12C-F). WT cells progressed from a UMAP region characterized by no autophagosomes (-N, 0 hours), through a region characterized by free autophagosomes (-N, early time points, 3 to 5 hours), and eventually to a region characterized by cleared autophagosomes, where autophagosomes fused with the vacuole (-N, late time points, 5 to 11 hours); (Fig 4A-B and S12A-B). Upon nitrogen replenishment, the WT cell trajectory reversed. As expected, the *atg1*Δ trajectory never progressed out of the ‘no autophagosome’ UMAP region. In contrast, the *vam6*Δ cell trajectory overlapped with the early WT cell trajectory but never advanced to the ‘cleared autophagosome’ UMAP region. Instead, it exhibited a distinct UMAP flux characterized by the progressive accumulation of free autophagosomes (Fig. 4A-B), which was statistically separable from that of WT cells in the UMAP within similar intervals of low confidence DNN predictions (classification probabilities in the range 0.1<P_1_<0.9; Fig. S12G-H). Given the substantial genome-wide variation in UMAP dynamics and many mutants exhibiting elevated classification uncertainty (Fig. S12E-F), we tested the use of the *vam6*Δ latent space as an “out-of-distribution” intermediate reference state in a two-step model framework for autophagy execution (Fig. 4C and S13A-E). Here, Bayes factors (BFs) were used to score the UMAP latent space evidence of mutant cells executing autophagosome formation (VAM6:ATG1) or clearance (WT:VAM6) based on the position of cells relative to reference kernel densities computed for WT, *atg1*Δ, and *vam6*Δ cells (Fig. S13B, C; details in Methods 6.2). We employed a time-independent BF, computed using time-varying kernel densities, to classify the overall mutant behavior between the three reference phenotypes (Fig. S13B, D). This procedure effectively integrated out the time variable by averaging the BF scores per mutant, enhancing the classification accuracy and sensitivity for phenotype detection (Fig. S12F-H). Additionally, we scored the time-wise evidence for the state of the mutants as they moved between fixed reference kernel densities throughout the time-course, indicating changes in autophagy execution activities (BF_t_) (Fig. S13C, E; Data File S7).

**Figure 4.**
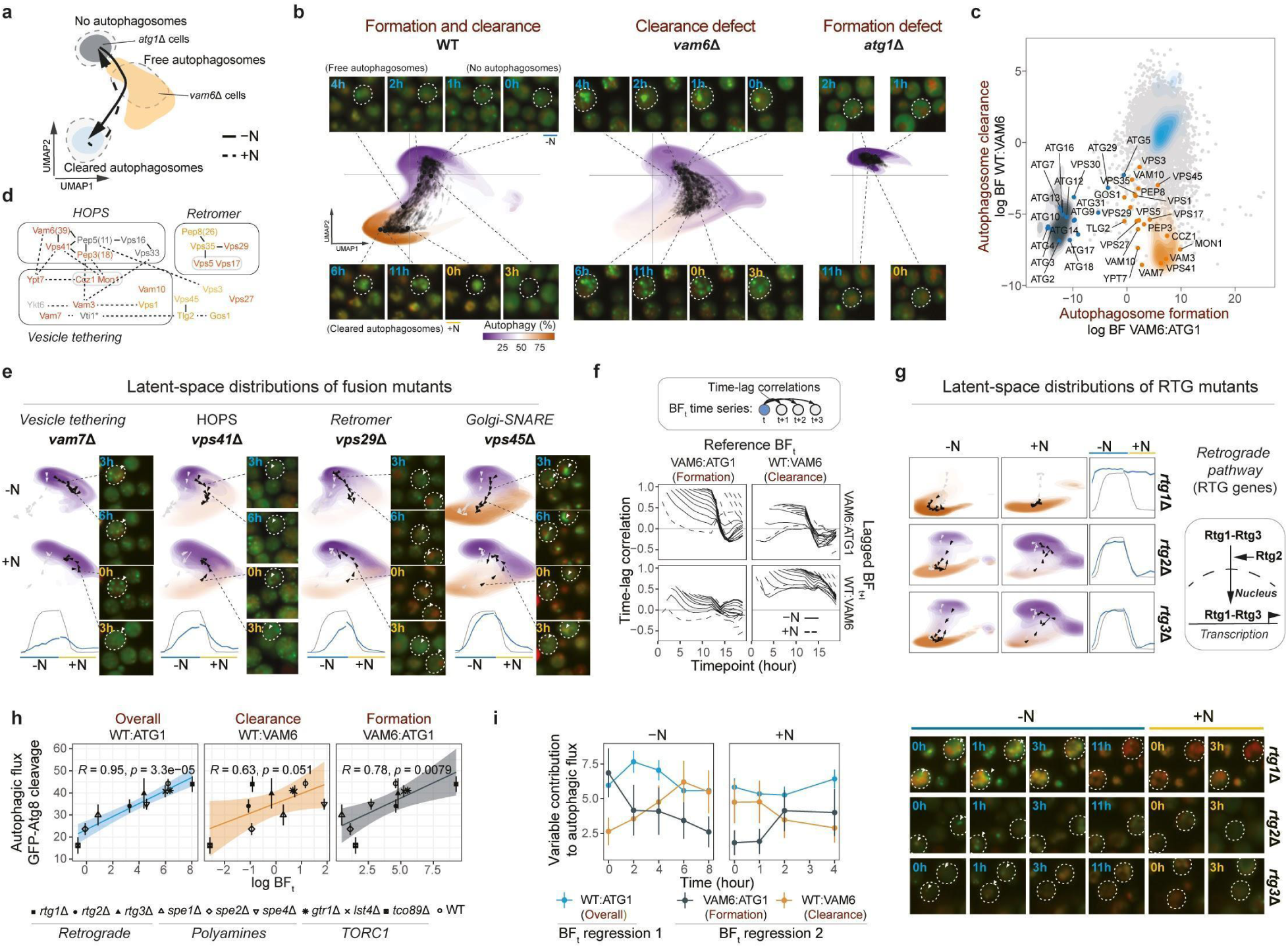
Dynamic latent-space analysis of autophagy execution. **a,** Latent-space model for autophagy execution. *atg1*Δ distribution overlaps with the region for ‘no autophagosomes’, *vam6*Δ distribution overlaps with the region for ‘free autophagosomes’, and the arrow represents the dynamic of WT cells as they execute autophagy. **b,** Comparison of UMAP-embedded latent-space distributions for WT (ORF) controls, *vam6*Δ, and *atg1*Δ cells with representative micrographs for the indicated time points. The densities represent distributions from six independent plates, grouped per time point and colored with the average autophagy prediction (%). Arrows indicate the position and direction of the average latent-space flux per plate. **c,** BF classification of autophagy execution phenotypes across the genome. Densities represent WT (blue), *vam6*Δ (orange), and *atg1*Δ (gray) cells. Reference ATG and fusion mutants are indicated and labeled. **d,** Color-coded functional overview of fusion mutants within each complex, ranging from red to orange, based on severity as determined by BF WT:VAM6. **e,** UMAP-embedded latent-space distributions and autophagy response dynamics for fusion mutants with representative micrographs from indicated time points. **f,** Correlations in genome-wide BFt variations for different time-lags (indicated by black arrows from BFt to BFt+l). **g,** UMAP-embedded latent-space distributions and autophagy response dynamics for *rtg1*Δ, *rtg2*Δ, and *rtg3*Δ with representative micrographs from indicated time points. **h,** Correlations of average BFt with autophagic flux measured as the average GFP-Atg8 cleavage (%) from 0, 2, and 4 hours in -N. *R* indicates the Pearson correlations, and *p* indicates their respective p-values. **i,** Relative contribution of log BFt scores in predicting autophagic flux per time point, estimated using mixed model regression (with time as grouping factor) based on a univariate model (WT:ATG1) or a bivariate model (VAM6.ATG1 + WT:VAM6). In *h* and *i,* points and bars indicate mean and standard error (see Data Table S12, *GFP quant.* for number of replicates per mutants and conditions).

The BFs accurately scored selective defects in autophagy execution, while also quantifying gradual increases in the severity of the mutant phenotypes (Fig. 4C-D, S13F-J, and S14A). For example, proteins involved in vacuole tethering and the HOPS complex exhibited a strong clearance defect and a high autophagosome burden, while retromer subunits and several other membrane trafficking proteins showed more moderate defects, with autophagosomes slowly docking and gradually resolving (Fig. 4C-E, and S14B). Analysis of correlations in genome-wide BF_t_ distributions between time points revealed highly nitrogen-dependent regulation of autophagosome formation, with a rapid decline in correlations between time-lags. This indicated that genetic contributions to autophagosome formation varied across distinct time intervals and treatment phases (Fig. 4F, upper left). In contrast, genetic perturbations in clearance activity remained relatively constant across time (Fig. 4F, lower right), with clearance in -N negatively correlating with the autophagosome burden in +N (Fig. 4F, upper *right*). These differences underscore the dynamic sensitivity of autophagosome formation to nitrogen availability, while revealing a steady and robust regulation of clearance activity through different mechanisms.

Parametric gene-set enrichment revealed that perturbations in amino acid synthesis or TORC1-mediated nutrient sensing were distinctly associated with elevated autophagosome formation, while perturbations in proteasomal activity and glucose uptake increased autophagosome clearance (Fig. S14B). Using time-independent BFs grouped over each treatment phase, we identified nitrogen status-dependent roles of some of these groups in autophagy execution (Fig. S14C-D). Interestingly, the deletion of genes in the RTG pathway^44^ resulted in excessive autophagosome formation in +N, and delayed clearance overall (Fig. 4G and S14E). *rtg1*Δ cells exhibited a particularly severe hyperactive phenotype that appeared decoupled from TORC1 signaling (with higher RpS6-p in +N, Fig. S15A). Moreover, all RTG mutants exhibited exhausted autophagic flux, with a moderate increase in autophagy in response to starvation as quantified by the delivery of distinct cargos, GFP-Atg8, and the modified phosphatase precursor Pho8Δ60, to the vacuole (Fig. S15A-B). This contrasted with spermidine synthesis mutants, which had a delay in autophagosome formation and clearance (Fig. S14E and S15C), and presented elevated RpS6-p levels in -N (Fig. S15D).

When measuring autophagic flux (GFP-Atg8 cleavage percent) across a set of selected mutants, the time-averaged BF_t_ WT:ATG1 yielded the best correlation out of any screen measurement (95%), supporting its use as a proxy for autophagic activity (Fig. 4H and S16A-B). Since this measure is the sum of the BF_t_ metrics with VAM6 as an intermediate step, we quantified the progressive contribution of formation (VAM6:ATG1) and clearance (WT:VAM6) activity on autophagic flux over time using a bivariate regression model (Fig. S16C-D). This analysis highlights the importance of clearance regulation, which emerges as the dominant factor for predicting Atg8 flux variation deep into starvation conditions and immediately upon replenishment (Fig. 4I).

### The TOR signaling pathway controls multiple stages of autophagy

Defects in the TORC1 signaling pathway caused nitrogen-dependent imbalances between autophagosome formation and clearance, varying by sub-complex (Fig. S14D-E, S15D-E). To further investigate these defects, we performed a secondary screen of selected mutants involved in TORC1 signaling, as well as the RTG pathway and spermidine synthesis. Upstream signaling components of TORC1 displayed substantial variations in the nutrient-gene interactions compared to the catalytic subunits (Fig. S14E). To isolate potential regulatory mechanisms independent of TORC1 phosphorylation, we treated mutants with rapamycin in combination with -N or +N for 10 hours and analyzed them using our deep learning approach (Fig. S17 and S18). Overall, rapamycin boosted autophagy induction and synergized with genetic disruptions of TORC1 signaling (Fig. 5A-C). Notably, rapamycin reduced autophagosome clearance activity in -N, as measured by the reduction in the BF_t_ WT:VAM6 values (Fig. 5A), an effect that was distinct from its impact in +N (Fig. S17G). This suggests a role of nutrient-regulated vacuole fusion mediated through TORC1 and other mechanisms. Interestingly, deleting the TORC1-independent Gtr1/2-Ltv1 branch^45^ selectively compromised clearance without inducing hyperactive autophagosome formation in response to starvation (Fig. 5B-D and Fig. S18). DAmP disruption of Tor2, Kog1 and Lst8, as well as non-essential deletion of Tor1 or the vacuole-specific component Tco89^46^, resulted in elevated autophagosome formation, particularly in +N or under rapamycin treatment (Fig. 5C-D). This outcome was also observed for several upstream positive regulators. However, disruptions in Lst8 and Tco89 caused rapamycin-independent clearance suppression, diminishing overall autophagy and diverging from the effects of disrupting the kinase domain components Tor1, Tor2, and Kog1 (Fig. 5C and Fig. S18). This clearance defect was validated in *tco89*Δ cells, which exhibited severely compromised autophagic flux in -N, as indicated by lower GFP-Atg8 cleavage (Fig. S15D).

**Figure 5.**
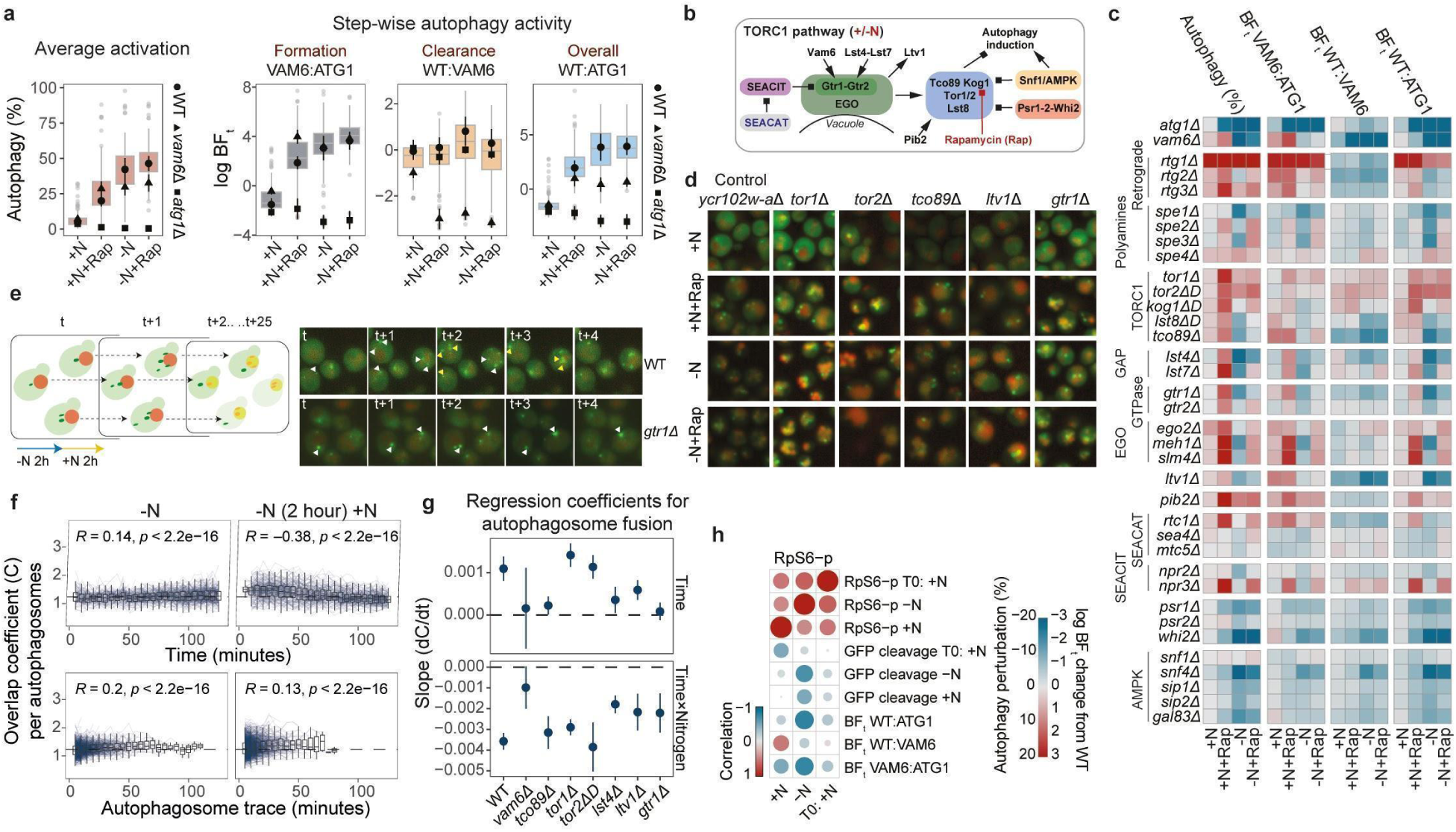
Multi-stage control of autophagy by the TOR signaling pathway. **a,** Average autophagy (%) and BFt levels in response to starvation, rapamycin or both. Points and bars indicate mean and standard deviation for the plate controls. **b,** Overview of components of the TORC1 pathway. **c,** Autophagy perturbation (%) and log BFt change from WT ORF controls for different mutants of the TORC1 signaling pathway, the RTG pathway, and the spermidine synthesis pathway in response to starvation, rapamycin, or both. **d,** Representative images of selected mutants after 3 hours of treatment in *c*. **e,** Segmentation and tracking of autophagic vesicles across time-lapse images, measured at 5-minute intervals. Fusion was quantified by computing the overlap coefficient per vesicle (see Methods 7.14); cell traces were collected from n=4 independent replicates. Representative images show fusion events indicated by yellow arrows. **f,** WT overlap coefficients per autophagosome over the treatment time (top panels) or life span of vesicle traces (*bottom panels*). Blue lines indicate individual vesicle traces. Dashed horizontal lines indicate the average of all measurements (including the mutants in Suppl. Figure *S19a-b)*. *R* indicates the Pearson correlations, and *p* indicates their respective p-values. **g,** Regression slopes indicating the changes in autophagosome overlap coefficients per mutant over time (*top panel*), and changes in slopes under addition of nitrogen (*bottom panel*). Points and bars indicate mean and 95% confidence intervals for the regression estimates. **h,** Correlations of RpS6 phosphorylation with GFP-Atg8 cleavage and average BFt scores for mutants in Suppl. Figure *S15a-d* (see Data Table S12, *Rps6 quant.* for number of replicates per mutants and conditions).

To investigate whether these effects were linked to an involvement of signaling components in nitrogen-dependent regulation of vacuole fusion, we performed high-frequency, high-resolution time-lapse imaging and tracked autophagic vesicles to measure potential changes in fusion rates in -N or after 2 hours of pre-treatment in -N followed by +N exposure (Fig. 5E). To robustly quantify the fusion events, we computed an overlap coefficient per vesicle (see Methods 7.14), which significantly increased over the vesicles’ lifespans in both conditions, except for *vam6Δ* mutants (Fig. 5F and S19A-B). Fusion rates were then estimated by performing a mixed model regression across the time series, with random intercepts for individual vesicle traces and interaction terms for potential treatment effects (Fig. 5G and S19C-D). Interestingly, while Tor1 and Tor2 disruption did not alter fusion rates compared to WT cells, *tco89*Δ, *lst4*Δ, *ltv1*Δ, and *gtr1*Δ cells exhibited severely delayed fusion in -N (Fig. 5G and S19B), corroborating the BF results (Fig. 5C). Additionally, *lst4*Δ, *gtr1*Δ, and *ltv1*Δ cells were significantly less sensitive to +N addition, which significantly decreased fusion rates in WT cells and other mutants except *vam6Δ* (Fig. 5G). This outcome suggests that the TORC1-independent Gtr1/2 signaling branch involving Ltv1 may directly regulate autophagosome fusion in a nitrogen-dependent manner. This multifaced modulation of TORC1 signaling, autophagosome formation, and clearance was further supported when comparing RpS6 phosphorylation with the BFs across TORC1-influencing mutants (Fig. S15)., Here, RpS6 phosphorylation was negatively correlated with autophagosome formation and autophagic flux across all conditions (Fig. 5H), reinforcing the role of TORC1 as a suppressor of autophagy induction. On the other hand, the analysis revealed a slightly positive correlation with clearance in +N (Fig. 5H), suggesting a decoupled regulation of fusion activity.

### Cross-omics inference reveals transcriptional mediators of autophagy regulation

Genome-wide quantification of dynamic signatures enables the construction of comprehensive systems-level models capable of predicting and simulating biological processes under different conditions, without sacrificing structural complexity or causal sufficiency. This approach has been successfully demonstrated in yeast using biologically informed machine learning models for growth and metabolism, leveraging the abundance of multi-omics data resources available^47,48^. As a proof-of-principle, we applied random forest to predict various autophagy perturbation metrics from several yeast gene deletion datasets. These datasets included transcriptomic, proteomic, and amino acid metabolomic features^49–51^, as well as genetic (SGA) and STRING-based genome-wide correlation networks^29,31^. All datasets were able to predict autophagy perturbations, with performance levels mirroring the experimental reproducibility of the different perturbation parameters (Fig. S20A and S8F). Notably, the genome-wide networks performed best overall, with test predictions achieving over 40% median correlations between predicted and true outcomes for the BF_t_ metrics in -N (Fig. S20A).

Given the prevalence of transcription factor (TF) deletions that strongly influenced autophagy dynamics, and the broader challenge of identifying the causal mediators through which TFs regulate cellular processes, we performed a mediation analysis to identify gene expression (GE) changes that predicted initial perturbations in autophagy dynamics. To achieve this, we analyzed the GE induction profiles of 186 overlapping TFs from the Induction Dynamics gene Expression Atlas (Fig. 6A)^52^.

**Figure 6.**
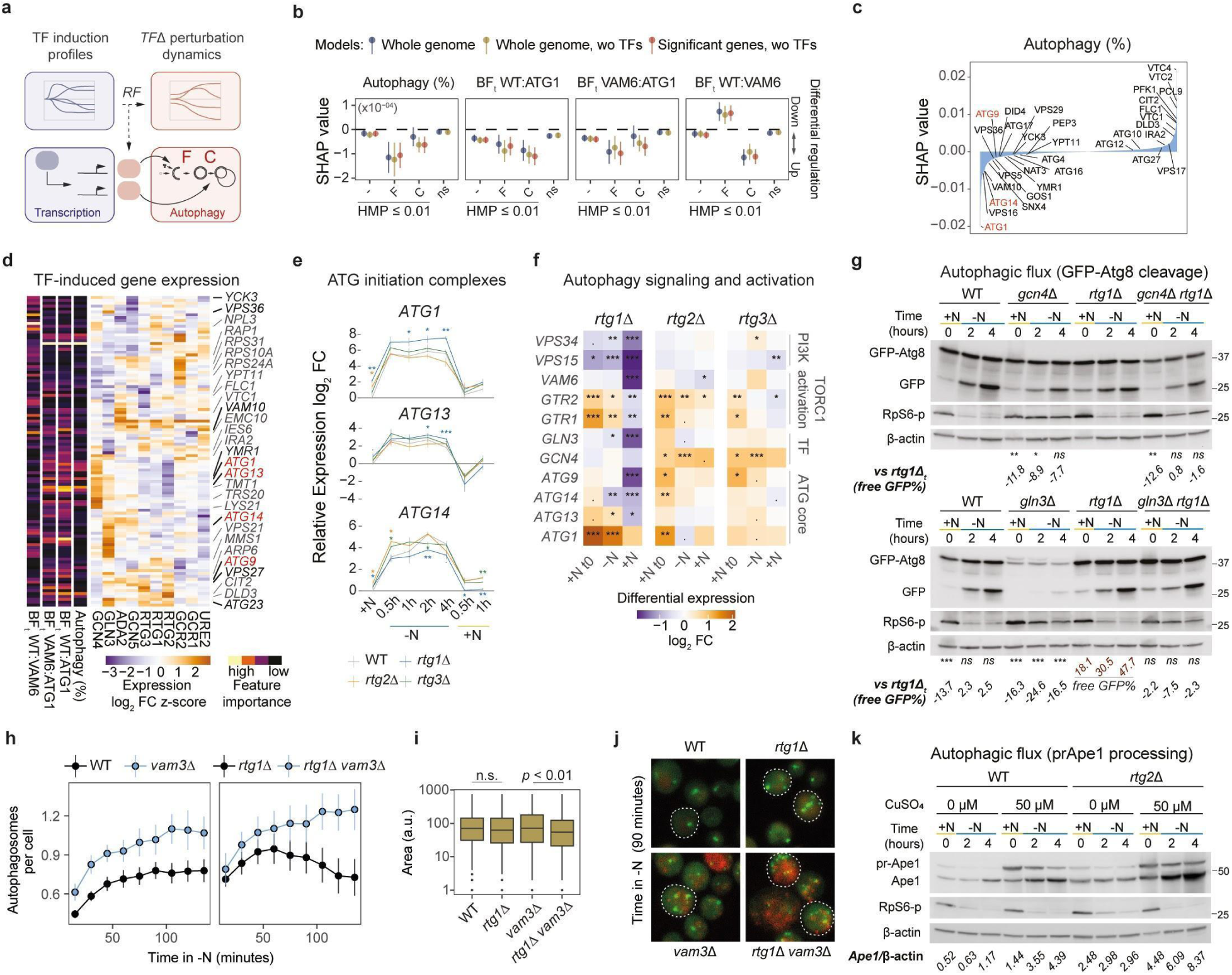
Cross-omics mediation analysis reveals transcriptional tuning of autophagy. **a,** Transcription factor (TF) gene expression induction profiles from Hackett et al.^52^ were used to predict the corresponding autophagy perturbation dynamics of TF gene deletions, using random forest (RF). Gene targets (model features) with high importance for predicting autophagy perturbations were evaluated as potential causal mediators of TF influence on autophagy. **b,** Adjusted SHAP values for differential regulation of gene expression comparing feature classes across models and autophagy metrics. F: formation genes; C: clearance genes with negative BFt z-scores beyond two standard deviations from the mean. Points and bars indicate mean and standard error. **c,** Gene-wise SHAP values for differential regulation of gene expression predicting autophagy responses (%). **d,** Hierarchical clustering of average TF-induced gene expression changes (z-scores) for gene targets that were selected as important model features (using sumGain). **e,** qRT-PCR for expression of *ATG1*, *ATG13*, and *ATG14* in *rtg1*Δ, *rtg2*Δ, *rtg3*Δ, and WT cells subjected to 4 hours of -N followed by 1 hour of +N. +N t=0h refers to cells before shifting to -N. Graphs show mean and standard error (n=3). **f,** Differential expression of various target genes in *rtg1*Δ, *rtg2*Δ, *rtg3*Δ cells per phase from (*e*) and Suppl. Figure *S20f*. *p ≤0.05;**p ≤0.01;***p ≤0.001; t-test in ***e*** and regression in ***f*** (n=3). **g,** Immunoblot analysis of GFP-Atg8 cleavage and RpS6 phosphorylation for WT, *rtg1*Δ, *gcn4*Δ, and *gcn4*Δ *rtg1*Δ cells (*upper panels*), and for WT, *rtg1*Δ, *gln3*Δ, and *gcn3*Δ *rtg1*Δ cells (*lower panels*) in -N for the indicated time points. +N t=0h refers to cells before shifting to -N. *p ≤0.05**p ≤0.01;***p ≤0.001; mixed model regression (n=4 for WT/*rtg1*Δ; n=2 for *gln3*Δ/*gcn4*Δ/double deletions). **h,** Average number of autophagosomes per cell in WT, *vam3*Δ, *rtg1*Δ, and *rtg1*Δ *vam3*Δ cells after 2 hours of nitrogen starvation-induced autophagy, measured at 15-minute intervals. Points and bars indicate mean and standard error (n=6). **i,** Area per autophagic vesicle in WT, *vam3*Δ, *rtg1*Δ, and *rtg1*Δ *vam3*Δ cells; t-test (n=6). **j,** Representative images for ***h-i,*** taken 90 min after nitrogen starvation. **k,** Immunoblot of Ape1 processing and RpS6 phosphorylation for WT and *rtg2Δ* cells in -N for the indicated time points, with or without overexpression of Ape1 through CuSO4 pre-treatment for 20 hours (n=2).

Models built with only potential causal mediators (i.e., significant genes, HMP ≤ 0.01, excluding the TFs themselves) were able to sustain predictive power, exhibiting only a moderate performance drop compared to genome-wide models (Fig. S20B). Moreover, expression changes for gene targets associated with severe autophagy perturbations were estimated to mediate stronger influences across all models (Fig. S20C), with targets involved in autophagosome formation or clearance acting as positive mediators of autophagy regulation overall (Fig. 6B and S20D). Interestingly, only targets involved in clearance were predicted to mediate a positive influence on clearance dynamics, whereas targets involved in autophagosome formation were predicted to exert a negative influence (Fig. 6B and S20D, *BF WT:VAM6 panels*). This suggests that transcriptional suppression of certain autophagy induction genes may play a role in promoting autophagosome clearance. *ATG1* and *ATG14* were identified as major mediators of autophagy initiation control (Fig. 6C), while *VAM10* and the ESCRT-0 subunit-encoding *VPS27* were identified as key mediators of autophagosome clearance regulation (Fig. S20E).

By clustering the average induction response of predicted mediators for select autophagy-regulating TFs, we found that RTG-mediated suppression of autophagy induction was likely mediated through Gcn4-regulated expression of *ATG1* (Fig. 6D)^52,53^. Indeed, RT-qPCR experiments revealed that *ATG1* expression was substantially greater in *rtg1*Δ cells than in WT control under both -N and +N conditions. Moreover, the effects of different RTG subunits on expression of ATGs and upstream TORC1 regulators, were decoupled during starvation (Fig. 6E-F and S20F-G). When *rtg1*Δ was crossed with *gcn4*Δ strains, the hyperactive autophagy phenotype was readily rescued; however, this effect was not observed in *rtg1*Δ*gln3*Δ strains (Fig. 6G). This epistatic effect of *rtg1*Δ on *gln3*Δ was consistent with *rtg1*Δ suppressing *GLN3* expression in +N, leading to decreased expression of Gln3-regulated gene targets in -N (e.g., *ATG14;* Fig. 6F)^41^.

The influence of the RTG pathway on autophagic flux was fully rescued in *rtg1*Δ *atg1*Δ strains (Fig. S21A). To confirm that Rtg1 exerts its effect specifically by controlling autophagy induction, we crossed *rtg1*Δ with *vam3*Δ strains to block autophagosome fusion and quantified the number and size of autophagosomes per cell using high-resolution imaging. Both *rtg1*Δ and *rtg1*Δ*vam3*Δ cells exhibited higher initial autophagosome counts than WT and *vam3*Δ cells (Fig. 6H). Upon starvation, *rtg1*Δ cells showed a transient increase in counts, which started to decline after 60 minutes, whereas *rtg1*Δ *vam3*Δ cells continued to accumulate autophagosomes throughout the entire time course (Fig. 6H, *right panel*). This difference in dynamics suggests that the larger autophagosome pools in *rtg1*Δ cells undergo a clearance burst upon starvation, an effect that is abolished in *rtg1*Δ *vam3*Δ cells. In contrast, WT and *vam3*Δ cells displayed a stable difference in autophagosome counts, with an asymptotic increase toward a steady state in both strains (Fig. 6H, *left panel*). These effects on clearance were further confirmed by tracking autophagosomes and measuring changes in overlap coefficients, showing that fusion was completely blocked by deleting *VAM3* (Fig. S21B). Interestingly, the hyperactive autophagosome formation in *rtg1*Δ cells was also associated with slightly smaller vesicles (Fig. 6I-J and S21B-C), suggesting potential exhaustion of resources. To investigate whether RTG pathway’s control of autophagy induction also influences cargo sequestration and clearance, we performed a giant Ape1 assay, in which we overexpressed Ape1 from a copper-inducible promoter to challenge autophagic capacity with large protein aggregates^54^. Because overexpression of Ape1 had a detrimental effect on growth in *rtg1*Δ strains, the assay was performed in *rtg2*Δ strains. Compared to WT cells, *rtg2*Δ cells exhibited an increased capacity to process Ape1 under both endogenous and overexpressed conditions (Fig. 6K), while also enhancing the membrane sequestration of Ape1 aggregates (Fig. S21D). Altogether, these findings highlight the RTG pathway as a crucial modulator of autophagosome formation.

## Discussion

Studying the dynamics of cell systems is becoming increasingly relevant as we strive to achieve predictability and guidance from our scientific models. This is particularly important for autophagy, which is intrinsically linked to an organism’s metabolism and growth and continuously fluctuates with environmental changes such as feeding cycles and circadian rhythms. To obtain a comprehensive understanding of autophagy activation during phases of declining nutrient levels requires deep insight into functional interactions with rate-limiting factors (Fig. 3G). Since genetic perturbations of the core machinery severely compromise autophagy execution, high-quality temporal data from regulating genes and multi-omics strategies to model changes in autophagy dynamics are essential. Our observation that there is a concordance between autophagy perturbation profiles and the direction of functional similarities with the core machinery (Fig. 3G and S11B), as well as a concordance in phenotypes between TF deletions and the predicted GE mediators (Fig. 6B), strongly supports the utility of the presented data.

The number of dynamic autophagy phenotypes detected with increased screening depth and sensitivity reflects the vast amount of cellular and environmental information integrated for optimal control of the process. The regulatory network complexity charted in this study is not surprising, given the critical role of autophagy in maintaining cellular homeostasis. By decomposing phenotypes into groups with distinct kinetic influences, we uncovered a regulatory asymmetry between the activation and inactivation of autophagy (Fig. 1G). Notably, the majority of mutants (*ultrasensitive*, *hyposensitive*, and *hyperactive*) influenced the timing of the starvation response. While a variety of activatory signals may be crucial for metabolic adaptation in a fluctuating environment, the timely shutdown of the process may be under stricter control to prevent excessive degradation and cell death. It is also possible that autophagy is subject to auto-inhibitory regulation through the depletion of ATG proteins, thereby increasing sensitivity to suppressive signals from nitrogen replenishment.

The six autophagy profiles discovered were remarkably well-organized across the genome-wide landscape (Fig. 2C–D, 3, and S11), with regulatory genes connected to the autophagy core machinery in a manner that reflected their tuning strength and functional characteristics (Fig. 3D, G). The main exception was the largest profile cluster, the *ultrasensitive* mutants, which exhibited weaker perturbation effects, no distinct network distribution, and no significant connection to the core machinery as a group. While this group includes several TOR signaling components, it likely also includes many genes with indirect effects on the starvation response (‘slope’) by influencing factors such as synchrony and cell cycle stage within the population. In contrast, *hyperactive* mutants exhibited stronger perturbation effects and were significantly closer to the core machinery, with multiple strong associations with ATGs, HOPS components, and vacuole tethering proteins (Fig. 3D, G). These findings suggest that further increasing overall autophagy during complete nitrogen withdrawal may require genetic interventions that engage multiple components of the process. Integrating phenotypic profiling data with network connectivity information could provide valuable design principles for modulating autophagy.

The genome-wide perturbations also provide a data source for parametric modeling of autophagy dynamics. In this study, we have provided some starting points by showing the predictive power across multiple genome-wide yeast datasets (Fig. 20A). By demonstrating the utility of this approach, we identified a novel phase-dependent tuning mechanism through the transcriptional repression of autophagy by the RTG signaling pathway. With Gcn4 positively regulating RTG gene expression (Fig. S20H)^52^, the retrograde pathway may function as a negative feedback loop, buffering a positive interaction between Gcn4 and Gln3. This mechanism supports temporal coordination of *ATG1* expression, limiting uncontrolled autophagy induction and promoting autophagosome clearance^52,53^. Recent discoveries suggest that Atg1 plays a role in activity-dependent disassembly of several ATG sub-complexes involved in autophagosome formation^55^. Thus, modulating the relative levels of ATG proteins may play a crucial role in tuning the strength and persistence of autophagy induction, and the imbalanced expression observed in RTG deletion cells could explain the diminishing returns in autophagic flux from starvation (Fig. 4G and S15A-B). This adaptive control of autophagy dynamics may be evolutionarily conserved, as the yeast retrograde response and analogous pathways in higher organisms support cellular survival in response to environmental perturbations or cellular challenges associated with aging^56^.

The versatility of our expectation-based deep learning approach provided a richer characterization of the autophagic feature space with high accuracy. Human annotation of heterogeneous data is a fundamental challenge for the utility of machine learning in cell biology^57^, while experimental generation of high-content reference data is becoming increasingly easy. Defining the ‘ground truth’ of a model using experimental control conditions can easily be extended to a range of other machine learning tasks^57^. When combined with a systematic analysis across functionally related gene sets, this approach revealed discrete phenotypes that might otherwise have gone unnoticed. An unexpected finding was the connection between the TORC1 pathway and autophagosome clearance. While this requires further investigation, one possibility is that TORC1 inhibition leads to a flux imbalance between autophagosome formation and clearance (Fig. 4H and 5), or that nutrient-reactivation of TORC1 promotes clearance of residual autophagosomes^39^. It has been shown that vacuole-associated activity of TORC1 depends on Tco89, while its endosome-associated activity and Atg13 phosphorylation remain intact in *tco89*Δ mutants^46^. Thus, removing Tco89 may lead to an imbalance between spatially and functionally distinct pools of TORC1. Additionally, Gtr1/2 regulation of Ltv1, which has previously been connected to endosomal membrane trafficking^45^, could play a role in supporting vacuolar fusion^26,58^. Treatment with rapamycin also revealed further levels of variation and otherwise unobserved nutrient-gene interactions (Fig. 5C and S17E-F). Hence, further manipulations of the culture environment and metabolic growth history can potentially unveil an even broader spectrum of unexplored autophagy regulatory mechanisms.

Due to the sensitivity of our approach, we were able to detect a vast number of genes with a significant influence on autophagy dynamics—approximately 30% of the investigated genome (Fig. 1). Our study also includes an analysis of essential yeast genes using the DAmP collection, which reduces protein expression by destabilizing mRNA molecules. While this approach does not always guarantee a hypomorphic phenotype^29^, it was preferred over conditional alternatives, such as temperature-sensitive alleles, which can introduce confounding effects on autophagy as a consequence of elevated temperatures^58,59^. Across the dataset, we observed library-wide differences in response kinetics and BFs, suggesting a potential confounding effect from strain background or DAmP treatment (Fig. 1B and S13J). However, discrete differences also emerged between the KO library and various WT control strains, varying according to the growth protocol used (Fig. 1G and S7F). Given the sensitivity of our methods and the multitude of factors influencing autophagy dynamics, these differences are unsurprising. Notably, essential genes were overrepresented among hyperactive and insufficient activation mutants (Fig. 1H), suggesting that elevated basal autophagy (Fig. S4A) may be a response to mRNA stress, potentially through stress granule formation, ribosome-associated quality control, or TOR-independent nutrient sensing^60–62^. Indeed, genes with these phenotypes were enriched in functional clusters related to nucleotide metabolism, RNA processing, and translation (Fig. 2C, S9, 3A, and S10A).

In conclusion, these data expand the knowledge base of autophagy-regulating genes, providing deeper insight into its phenotypic landscape within a broader regulatory context. The resulting genome-wide profiling repository not only represents a foundational reference for systems-level autophagy characterization, but also serves as a powerful hypothesis-generating tool. It is publicly available via the AutoDRY web portal (Autophagy Dynamics Repository in Yeast): https://cancell.shinyapps.io/autodry/. Our study thus presents a time-resolved, genome-wide map of autophagy activation and inactivation, offering a comprehensive view of its dynamic regulation in cellular nutrient responses.

## Supplementary Materials

Materials and Methods, Figs. S1 to S21, Table S1, References *63–103*, and Data S1 to S19, available for download from DRYAD. DOI: 10.5061/dryad.cfxpnvxdh

## Code availability

All code is freely available at https://github.com/Enserink-lab. For any additional information required for reanalysis of the data reported in this study, please contact the corresponding authors.

## High-content Image availability

All high-content images will be publicly available in the BioImage Archive at EMBL-EBI: https://www.ebi.ac.uk/bioimage-archive/. Additionally, high-content images will be shared by the lead contact upon request.

## Genome-wide profiling repository

All mutant profile autophagy responses are publicly available on the AutoDRY web portal: https://cancell.shinyapps.io/autodry/.

## Acknowledgements

We thank Dr. Anne Simonsen and Dr. Harald Stenmark for their critical comments on the manuscript. We also thank Dr. Eduardo Cebollero and Dr. Fulvio Reggiori for providing their valuable guidance on the experimental protocol for alkaline phosphatase (ALP) assay. We thank Dr. Claudine Kraft for providing reagents for Ape1 assay. We acknowledge Dr. Anna Lång and Dr. Vigdis Sørensen and Advanced Light Microscopy Core Facilities at Oslo University Hospital Gaustad and Montebello for technical support. We thank Dr. Karolina Spustova for her technical assistance with the Nikon live-cell imaging. We thank members of the Enserink group for their feedback and suggestions. This work was funded by Norwegian Cancer Society (project numbers 182524 and 208012), Norwegian Health Authority South-East (2017064, 2017072, 2018012, 2019096), Research Council of Norway (261936, 301268), Research Council of Norway through its Centres of Excellence funding scheme, project number 262652, and Fundación Alfonso Martín Escudero (SMO).

## Author contributions

Conceived the study and acquired funding: J.M.E. Conceptualization: N.C., A.N.A. and J.M.E. Methodology – construction genome-wide libraries: I.G., A.N.P., P.A.D. and C.D.P. Methodology – optimizing cell-growth and starvation protocols: N.C. Methodology – developing and optimizing high-content screening protocols: N.C., S.O.M. and A.N.P. Methodology – construction of validation libraries: N.C., S.O.M. and L.H. Methodology – image segmentation and feature extraction analysis: N.C. and I.G. Methodology – developing and optimizing deep learning pipeline: A.N.A. and I.G. Investigation – performing and validating high-content screening: N.C. and L.H. Investigation – optimization of autophagy assays: N.C., L.H., S.O.M., A.N.P., and E.R. Investigation – performing RTG, TOR, and SPE experiments: N.C., L.H., and E.R. Investigation – Electron microscopy and analysis: S.W.S. Data curation and analysis: N.C. and A.N.A. Formal analysis and statistics: A.N.A. Network analysis: N.C. and A.N.A. Visualization: N.C. and A.N.A. Web portal: S.N. and A.N.A. Writing – original draft: N.C. and A.N.A. Writing – review & editing: N.C., A.N.A., C.D.P., M.Z., T.E.R., and J.M.E. Project coordination: N.C. and J.M.E. Supervision: N.C., M.Z., T.E.R., and J.M.E. All authors discussed, revised, and approved the manuscript.

## Supplementary Materials for

### Materials and Methods

#### 1. High-content screening

##### 1.1 Genome-wide libraries details

The yeast mutant libraries harboring the autophagy reporter (autophagy library KO-A and DAmP-A) were constructed based on the collection of non-essential deletion mutants (YKO collection; Open Biosystems), and the collection of essential genes with a decreased abundance by mRNA perturbation (DAmP collection; Open Biosystems). The DamP mutations were chosen over conditional temperature-sensitive (ts) mutations to eliminate temperature as a variable that could impact the autophagy response^58,59^. The mutant strains were originally distributed across 56 KO-A and 12 DAmP-A 96-well plates. To introduce control strains into each plate, the mutant strains in row A were transferred to new plates designated as Mixed plates. The mutants with significantly reduced growth rates, that may affect the cell count for imaging, were re-inoculated at higher density into independent plates that were annotated as Recovery plates. A subset of mutants was rearranged into Repetition plates, which were used for validation and reproducibility experiments. Rapamycin plates were prepared for the analysis of rapamycin response.

Each plate included three control strains in triplicate, specifically the SGA starting strain WT (Y7092)^63^, the deletion mutant *vam6Δ::kanMX4* (KO-A), and the deletion mutant *atg1Δ::kanMX4* (KO-A). The libraries Mix, Recovery, Repetition and Rapamycin included the deletions of three dubious ORFs, *ydr535cΔ::kanMX4* (KO-A), *ycr102w-aΔ::kanMX4* (KO-A), and *yel074wΔ::his3MX6* (this study), in addition to *atg1Δ* and *vam6Δ,* to control for a potential effect of the kanamycin and histidine selectable markers on the autophagy response. The complete list of mutant strains and controls distributed into the arrays KO-A, DAmP-A, Recovery, Mixed, Repetition, and Rapamycin is provided in Data File S1.

##### 1.2 Genome-wide libraries preparations

To prepare genome-wide libraries for high-throughput screening, duplicates of the yeast libraries were created in 96-well plates (Nunc™ Microwell™, Thermo Fisher). In each well, 1 µl of both control strains and mutant strains was inoculated into 150 µl of synthetic medium containing amino acids (SC: 6.9 g yeast nitrogen base with ammonium sulfate and without amino acids, 20 g/ glucose anhydrous, and 0.790 g Complete Supplement Mixture Formedium™ per liter). To promote optimal growth and low cell death rate, the duplicate plates were incubated at 25 °C for 72 h, before being stored at −80 °C with 20% glycerol.

A working batch of 16 plates from duplicates was prepared five days before an imaging round. In each plate, 1 µl of both mutant and control strains was inoculated into 200 µl of SC medium. Following a 48-hour incubation at 25°C, these plates were stored at 4°C with a cover film. The batch was divided into four sets, and each set underwent pre-culturing and imaging on consecutive days.

##### 1.3 Cell growth and starvation protocol for high-content screening

The growth protocol was optimized to enable high-throughput cell growth while inducing an optimal autophagy response in yeast cells. This involved periodic medium replacement, which kept cells metabolically active and prevented premature autophagy initiation due to nutrient unavailability upon prolonged incubations without shaking. It ensured that yeast cells were in the log phase and minimized population response variability before the onset of starvation.

After a 48-hour incubation of the previous batch, 1 µl of strains with a normal growth rate and 4 µl of strains with a reduced growth were transferred into 200 µl of fresh SC medium. These plates were then incubated at 25°C for an additional 48 h, followed by reinoculation and another 24-hour incubation at 25°C. For time-lapse imaging, 4-6 µl of cells were inoculated into 96-well black plates (Corning® Black microplate, Sigma-Aldrich) coated with Con A (0.25 mg/ml) and incubated for 8 hours at 25 °C to ensure cells were in exponential phase. To induce autophagy, cells were washed twice with sterile MilliQ water and transferred to a nitrogen-starvation medium for 12 h (SD-N: 1.9 g yeast nitrogen base without ammonium sulfate and without amino acids, and 20 g glucose anhydrous Formedium™ per liter). Subsequently, the same cells were shifted to SC medium (at t=12 h) to record autophagy inactivation upon nitrogen replenishment for 7 hours. Live imaging was performed at 30 °C, capturing images every hour for 19 hours. The cells were maintained at room temperature during the intervals between each time point of image acquisition.

##### 1.4 Autophagy reporter details

To assess autophagy activity, translocation was monitored of fluorescent protein-tagged Atg8 from the cytoplasm to the vacuole using fluorescent microscopy^64^. For time-lapse experiments we used the optimized mNG fluorescent protein fused to the N-terminus of Atg8 under the control of the ScTDH3 (GPD) promoter. To visualize the vacuole, the mCherry fluorescent protein was fused to the C-terminus of the vacuolar proteinase A Pep4.

##### 1.5 Plasmid construction

The pYM-N natNT2-GPD1pr-mNeonGreen plasmid was generated by replacing the yeGFP coding sequence from pYM-N17^65^ by the codon-optimized monomeric NeonGreen (mNG) DNA sequence^66^. The mNG sequence was amplified from the pFA6-mNG-his3MX6 plasmid using primers mNG-Fwd-XbaI and mNG-Rv-EcoRI, which included *XbaI* and *EcoRI* sites for the tag replacement (see Data File S1, *primer sequences*).

##### 1.6 Integration of autophagy reporter into libraries and SGA query construction

The autophagy reporter was crossed into the collections of non-essential and essential mutants to create the YKO-A and DAmP-A collections, respectively. Triple allele strains harboring the autophagic reporter and the deletion of non-essential or DAmP alleles were obtained by adapting the Synthetic Genetic Array (SGA) analysis method^67,68^.

The query bait containing autophagy reporter was created using the SGA host strain *MAT*α haploid Δ*can1*::*STE2*pr-Sp_*his5* Δ*lyp1* (Y7092; kind gift from C. Boone, University of Toronto, Canada). Endogenous *ATG8* and *PEP4* were tagged with mNG and mCherry using a standard PCR-based approach ^65^. In brief, *natNT2:GPD1pr-mNG-ATG8* was constructed by PCR amplification of a *natNT2-GPD1pr-mNeonGreen* fragment from the pYM-N17-mNeonGreen plasmid using the primer pair ATG8-S1 and ATG8-S4, whereas *PEP4-mCherry:hphMX6* was constructed by integrating an *mCherry:hphMX6* PCR fragment amplified from plasmid pBS35 using primer pair PEP4-S3 and PEP4-S2 at the 3’ end of the *PEP4* locus (see Data File S1, *primer sequences*). This resulted in the construction of JEY11511 (MAT*α can1Δ::STE2pr-Sp_his5 lyp1Δ his3Δ1 leu2Δ0 ura3Δ0 met15Δ0 natNT2:GPDpr-mNeonGreen-ATG8 PEP4-mCherry:hphMX6*).

The SGA bait strain JEY11511 was crossed with a target array of 4760 viable MATa deletion mutants and 1159 conditional alleles of essential genes (KO and DAmP mutants respectively) in the BY4741 background (MAT**a** *his3Δ1 leu2Δ0 met15Δ0 ura3Δ0*), wherein each ORF or 3’ UTR was disrupted by a kanMX4 resistance cassette marker. Crossing of JEY11511 was performed in quadruplicate by pinning onto fresh YPD agar plates using a Singer RoToR (Singer Instruments) followed by growth for two days. Cells were subjected to two rounds of pinning onto diploid selection medium (YPD-agar containing 200 μg/ml geneticin (G418, ThermoFisher) and 200 μg/ml hygromycin (Gibco)) and grown for 1-2 days at 30°C. The cells were then pinned onto pre-sporulation medium (15 g Difco nutrient broth (BD Biosciences), 5 g Bacto-yeast extract (ThermoFisher), 10 g Bacto-agar (ThermoFisher), and 62.5 ml 40% glucose (ThermoFisher) per 500 ml) and grown for 3 days at 30°C. Cells from the pre-sporulation medium were then pinned onto sporulation medium (10 g potassium acetate, 0.05 g zinc acetate, 20 g Bacto-agar per liter, containing a final concentration of 50 μg/ml G418 and 50 μg/ml hygromycin) and incubated for 7 days at 30°C. The resulting spore-containing cells were then subjected to two rounds of pinning onto diploid killing medium (1.7 g yeast nitrogen base without amino acids and without ammonium sulfate (BD Biosciences), 1 g L-glutamic acid monosodium salt, 2 g SC dropout mix without lysine, canavanine, and histidine (US Biological), 20 g Bacto-agar, 50 ml of 40% glucose per liter, containing a final concentration of 50 μg/ml thialysine (Sigma Aldrich), 60 μg/ml canavanine (Sigma Aldrich), 200 μg/ml G418 (Gibco), 200 μg/ml hygromycin (Invitrogen), and 100 μg/ml nourseothricin (Jena Bioscience)) followed by growth for 5 days at 30°C for the first pinning and 2 days at 30°C for the second pinning. Cells were then subjected to two rounds of pinning and growth on haploid selection medium (1.7 g yeast nitrogen base without amino acids and without ammonium sulfate, 1 g L-glutamic acid monosodium salt, 2 g SC dropout mix without histidine, 20 g Bacto-agar, 50 ml of 40% glucose per liter, containing a final concentration of 200 μg/ml G418, 200 μg/ml hygromycin, and 100 μg/ml nourseothricin) and grown for 2 days at 30°C. Then the cells were pinned and grown on YPD-agar followed by storage at −80°C in YPD media containing 20% (v/v) glycerol. The resulting MATa haploid triple mutants were used for high-content screening.

##### 1.7 High-content fluorescent imaging

The high-content screening (HCS) for autophagy analysis was carried out in a 96-well plate format using an ImageXpress® Micro Confocal microscope equipped with a temperature control unit, LED light source, and specific filter sets (Molecular Devices). The images were acquired using automated acquisition in wide-field, utilizing the Nikon Plan Apochromat 60x/0.95 Air objective coupled to a 16-bit 4MPix scientific CMOS camera and a laser-based autofocusing unit. Image acquisition was controlled by the ImageXpress Analysis Software (Molecular Devices). Live cell imaging was conducted at 30 °C ± 2 °C. Fluorescence signals were captured using specific filter sets for FITC and TexRed, with exposure times of 200 ms and 800 ms, respectively. For transmitted light (TL) images, an exposure time of 10 ms was used. The same cells were imaged every hour for 19 hours with 2-3 images per well by selecting adjacent imaging sites near the center of the well and employing modus focus on the plate and bottom to maintain the same plane.

##### 1.8 Automatic image analysis using Fiji/ImageJ for high-content screening

We developed an automated image analysis tool using Fiji/ImageJ to extract features from the genome-wide dataset and evaluate changes in the autophagy reporter along the time-lapse experiments. The tool employed a macro to extract parameters for each image categorized by well (row+col), time point (starvation/replenishment), image within well (s1, s2, s3), and channel (w1=BF, w2=green, w3=red). The extracted features included parameters related to both fluorescent signals and cell morphology. Parameters related to fluorescent signals were mean and standard deviation of signal intensity, modal and minimum signal values, and integrated median signal intensity. Parameters related to cell morphology included *Cell Area, Centroid,enter Coordinates, Perimeter, Bounding box, Best-fit Ellipse Dimensions, Shape Descriptors, Ferret Dimensions, Skewness, and Kurtosis*.

For image segmentation, we first enhanced contrast using a saturation value of 0.3 and then sharpened it to improve clarity and pixel contrast. We further enhanced contrast with the same saturation value to refine the image after sharpening. We then applied the ‘Unsharp Mask’ plugin with a radius of 5 and a mask value of 0.70. We used the ‘Auto Threshold’ plugin with the default method and set the image to white. To finalize the segmentation and make objects white on a black background, we inverted the image and applied the ‘Minimum’ filter with a radius of 0.5. To eliminate small holes or gaps in the image, we used the ‘Close-’ plugin and followed it with another inversion of the image.

To identify and count the objects in the image, we used the ‘Analyze Particles’ plugin with the size range of 400-2500 and circularity range of 0.50-1.00. After correcting the background using a rolling value of 50px and measuring the signal intensity of ROIs in both channels, we quantified colocalization by multiplying the pixel values of the two channels with the ‘multiply create 32-bit’ command.

#### 2. Deep learning analysis

##### 2.1 Data preparation for model training

All neural net models were trained using the *Keras* API (version 2.2.5.0) for *Tensorflow* (version 2.00) in R (version 3.6.3)^69^. The training data with associated state labels was compiled from *atg1*Δ or WT cells under the experimental conditions where autophagic or non-autophagic features were uniformly maximized across the cell culture populations (Fig. S1A). *atg1*Δ cells across the entire experimental protocol or WT cells in T0 or from 5 hours in nitrogen replenishment were assigned state 0 (non-autophagic), while WT cells from 8 hours to 12 hours of nitrogen starvation were assigned state 1 (autophagic). Although autophagic features, including free mNeonGreen-Atg8 puncta, and colocalization of mNeonGreen-Atg8 and Pep4-mCherry, could be observed earlier in WT cells, 8 hours of starvation represented a demarcation where mNeonGreen-Atg8 and Pep4-mCherry were clearly overlapping homogeneously across all WT cells, representing that autophagosomes had been delivered to the vacuole. Experimental outliers visibly deviating from these standards were manually removed from the training data. Also, cells with low mNeonGreen-Atg8 signal (mean intensity < 150) or low (mean intensity < 120) or high (mean intensity > 5000) Pep4-mCherry signal were removed from the training data.

Single-cell parameters for variations in signal distributions from both color channels and the co-occurrence were used as input features. Prior to training, we computed the mean and standard deviation feature vectors from the entire WT dataset, including all timepoints, which were used to transform the training data features and transforming all future new data. The resulting training set included 1 073 981 observations in state 0 and 269 655 observations in state 1.

##### 2.2 Model training, hyperparameter search and model selection

To train the models binary cross-entropy was used as loss and the models were trained using the AdaGrad algorithm, with a batch size of 50 000, learning rate of 0.01, and patience of 5 epochs. In the hyperparameter search, 10% of the data was kept out for testing. Of the remaining dataset 80% of the data was used for training the models and 20% was used as validation data for early stoppage. To find the optimal model architecture a grid search was performed over all combinations of activation functions (*relu*, *elu*, *tanh, sigmoid*) with networks of different sizes (Fig. S1A). The networks were grown automatically by defining a number of layers (1-5 and 6-8, respectively) and neurons in the first layer (60-120 and 80-120, respectively, with increments of 5) and adding the layers with a decay rate in the number of neurons from the former layer (0.5-1 and 0.7-1, respectively, with increments of 0.05), with a minimum layer size of 10 neurons. This resulted in 3096 models that were trained for a maximum of 150 epochs (with early stoppage). The models were evaluated based on the test set prediction averages compared respective state labels, training and validation set loss and accuracy, and model overfitting (Fig. S1B-D). With exception of *sigmoid*-based models, which underperformed compared to the other models, most architectures resulted in similar test performances (Fig. S1B). Thus, the *sigmoid*-based models were excluded from further analyses.

Model complexities were quantified from the number of model weights. Since model test and validation performances were asymptotically scaling with model complexity, while more complex models also tended to have more overfitting for some architectures (*relu*), we defined a model complexity cutoff at the smallest model in the top 1% (validation loss; Fig. S1D). Moreover, network size may affect the rate of convergence and generalization of a model for complex function approximations, where 150 epochs may not be enough. Subsequently, we selected the top-performing 5 models (validation loss) among the least and most complex models (given the complexity cutoff) for *relu*, *elu*, and *tanh* activation functions, respectively (Fig. S1E). These 30 models were re-evaluated by re-training them for a maximum of 3000 epochs (with early stoppage) using all of the data in a 80/20 training and validation split. The models were evaluated based on training and validation set loss and accuracy, as well as the between-experimental correlation of predicted autophagy dynamics (plate averages for WT, *atg1*Δ, and *vam6*Δ data points combined). On average, the less complex models slightly outperformed the more complex ones for architectures using *tanh* and *elu*, but not *relu* activation functions (Fig. S1E).

Additionally, the predicted last latent layers (autophagic feature vectors) for the top 2 models (validation loss) for each activation function, were extracted for 500 000 randomly sampled WT, *atg1*Δ, or *vam6*Δ cells from any time point and embedded with uniform manifold approximation and projection (UMAP; Fig. S1A), using the *umap* package (version 0.2.7.0) in R^70^. The UMAP was used to force a low-dimensional and regularized representation of the predictive information in the neural networks and to investigate the relative contribution of noise and extracted autophagic features between three distinct autophagy phenotypes. The UMAP distance metric was set to Pearson correlation, and n_neighbors were set to 10, 15, or 30. The between-experimental correlation of predicted UMAP dynamics (coordinate-wise plate averages for WT and *vam6*Δ data points combined) was used to test latent space robustness (Fig. S1E). The two highest performing and most robust models across multiple metrics (*elu* model 30 - 7 layers, 90 neurons in layer 1 and 66 neurons in layer 7; *tanh* model 22 - 8 layers, 115 neurons in layer 1 and 37 neurons in layer 8) were deployed for GW predictions, and used to corroborate each other (Fig. S1H). Statistical analyses of percent autophagy response dynamics (Fig. 1-3 and ‘Autophagy %’ results of Fig. 5) were based on predictions from model 30. For latent space analyses, model 22 had less noise and were primarily used to visualize the UMAP dynamics (Fig. 4), but all statistical analyses and inferences from the UMAP distributions were averaged or corroborated by predictions from both models (Fig. 4-5).

##### 2.3 Parametric UMAP

Since transforming new data with UMAP is slow, we implemented a fast parametric transformation of the inputs (autophagic feature vectors) to the embedded coordinates using a neural net (pUMAP: fully connected, 8 layers, 120 neurons/layer, *tanh* activation functions, with bivariate linear output, and trained using the adam optimizer with MAE loss; Fig. S1A). The specific implementation here was conceived before the publication by Sainburg et al. 2020^71^, but follows the same essence.

##### 2.4 Genome-wide predictions and quality control

New screen data were filtered for cells with low (mean intensity < 120) or high (mean intensity > 2000) Pep4-mCherry signal. A more stringent upper bound was used for the red channel in order to ensure the removal of dead autofluorescent cells. The input features were z-transformed using the feature mean and standard deviation from the training data, and the probability for cells activating autophagy (P_a_) was predicted using the neural nets. Subsequently, the expected probability of autophagy (predicted autophagy %), 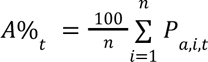, was computed for the mutant cell population at every time point. Alternatively, we also computed the autophagy classification %, 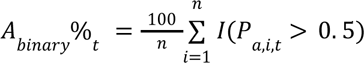, which strongly correlated with *A*%_*t*_, but was not used as it was less robust for smaller cell populations. The expected standard deviation per cell was computed as 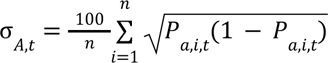, to assess the prediction uncertainty. We also computed the classification uncertainty using the entropy, 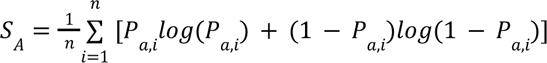 (See Data File S2, *autophagy % DNN predictions*).

Outlier plates were identified by computing the *IQR* from all plate median responses and finding plates that had a mean absolute deviation greater than 2*IQR* over the time series. These plates were subsequently repeated. We also identified mutant outliers based on the 99% high density region of average cell numbers and expected standard errors over the time course (Fig. S2E), which were subsequently repeated in new experiments with higher initial cell seeding concentration. Four DAmP plates that did not have the replenishment response, were manually removed from genome-wide analyses and clustering procedures.

#### 3. Statistical analyses

##### 3.1 Double-sigmoidal model fitting

Double-sigmoidal models were fitted to the data using the *sicegar* package (version 0.2.4) in R (version 4.2.1)^27^. This model includes six parameters (maximum autophagy, final autophagy, two transition midpoints/T50s, and two slope parameters) (Fig. 1C). Tangent lines crossing the transition midpoints (T50s) were computed to extract additional parameters describing the dynamic part of the curve. To de-noise, interpolate potential missing time points, and standardize responses with minor shifts in imaging times, the models were also used to predict the response for all mutants for the entire time series with 30-minutes increments. The goodness-of-fit was assessed using the log-likelihood (on normalized data) and RMSE (See Data File S2, *autophagy % curvefits* and *goodness-of-fit measures*).

##### 3.2 Hypothesis testing and perturbation parameter statistics

Using the double-sigmoidal model predictions, autophagy perturbations were computed from the difference between the interpolated curve fits (*A_f_*%) for all mutants (*m*) and the corresponding plate medians (ctr), Δ_*f*_%_*m,t*_ = *A*_*f*_%_*m,t*_ − *A*_*f*_%_*ctr,t*_, and subsequently averaged per phase (Perturbation -N or +N) or over the entire time series (Perturbation overall; Fig. S3D). Equivalently, all the mutant parameters were centered by subtracting the corresponding plate medians to compute parameter perturbations, Δ*parm*_*m*_ = *parm*_*m*_ − *parm*_*ctr*_ (Fig. S3E).

Autophagy perturbation/parameter errors were computed for each of the plate controls (WT, *atg1*Δ, and *vam6*Δ), by subtracting their respective plate-wise average. Then, the WT errors were used as H_0_-references for each parameter by fitting the error distributions using the *locfit.raw* function of the *locfit* package (version 1.5-9.8) in R^72^, and assessing significance of the parameters by computing p-values from the re-sampled fitted H_0_-densities. To control for the false discovery rate (FDR) the parameter p-values were adjusted using the Benjamini-Hochberg method (see Data File S2, *parameter statistics and parameter controls*)

##### 3.3 p-value aggregation and mutant statistics

Due to the high correlation between the different perturbation parameters, causing a lack of independence between the individual hypothesis tests (non-independent p-values), we used a harmonic mean p-values (HMP) technique to aggregate the p-values and score the overall significance of the mutants, while controlling for the false discovery rate^28^. The HMPs were computed as 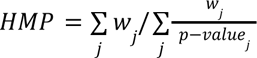, where weights corresponded to the H_0_-rejection rate at a 5% Benjamini-Hochberg (BH) correction cutoff, *w_j_* = *F_j_* (*FDR_BH_* ≤ 0. 05), in order to favor the more likely alternative hypotheses based on the genome-wide data. This grouping of hypothesis tests maintained sensitivity of the least stringent BH-adjusted p-value (FDR min) or improved the sensitivity for mutants with multiple significance tests. HMPs based on equal weights were also computed, but in comparison did improve the power over individual BH-FDR corrections per mutant (see Fig. 1D and S4B and Data File S2, *mutant profiles and parameter groups*).

##### 3.4 Mutant profiling and clustering

Statistically significant mutants, HMP ≤ 0.01, were extracted for clustering. The parameter perturbations were clustered and grouped by computing the log_10_-transformed p-values and performing a hierarchical clustering using Pearson correlation as a similarity metric and ward.D2 clustering. The parameters were split into five groups (Fig. S4D) using the *cutreeDynamic* function from the *dynamicTreeCut* package in R^73^.

To group the mutants into distinct dynamic profiles, the response predictions from the double sigmoidal models for mutants with HMP ≤ 0.01 were subjected to hierarchical clustering using Euclidean distances and ward.D2 (Fig. S4E). For mutants with multiple significant independent experimental measurements the median responses were computed before the analysis. Groups were identified with the *cutreeDynamic* function.

#### 4. Screen quality assessments

##### 4.1 Recall of reference mutants

‘Gold-standard’ test sets for evaluating the sensitivity and accuracy for retrieving true autophagy perturbation phenotypes, were defined through manual inspection of images and literature curation (Table S1)^5,25,26,58,74–81^. This included two true positive test sets, including genes involved in autophagosome formation (‘ATG set’, n=16) and clearance (‘Fusion set’, n=19), respectively. A negative control set was also developed from several independent replicates of two dubious ORFs^82^ that produced WT autophagy responses (n = 23) (Table S1). None of the mutants included in the test sets were part of the training data or used as reference data for the UMAPs.

Precision-recall curves (PRC) and receiver operating characteristic (ROC) curves were computed from the p-value rank orders for the different perturbation parameters, comparing the true positive sets (‘ATG set’, ‘Fusion set’, or combined) with the negative control set (Fig. S5C, E), and area under the curves (AUCs) were computed to score the precision and accuracy of recalling true positives (Fig. S5B,G). Alternatively, negative sets with equal size to the true positive sets were bootstrapped by repeated random sampling with replacement from the GW population (b=1000) for a more statistical assessment of the different perturbation parameters with respect to the average deletion mutant phenotype (Fig. S5A, B and S5D, F, G). For mutants with multiple independent experimental measurements the median p-values or HMP were computed before the analysis. To use ROC curves for a direction-specific separation of the test sets from the GW population average, the p-values were log-transformed and signed by the direction of the perturbation (Fig. S5F, G).

##### 4.2 Gene set enrichment analysis (GSEA)

GSEA was performed using the *gseGO* function of the *clusterProfiler* package (version 4.4.4) and *org.Sc.sgd.db* (version 3.15.0) in R (version 4.2.1)^83^, with 10^5^ permutations, and min and max gene set size set to 5 and 200, respectively. To analyze the directional association of GO-BP (gene ontology - biological processes) terms for the individual parameter perturbations, we used the log-transformed p-values signed by the direction of the perturbation. For mutants with multiple independent experimental measurements the median p-values were computed before the analysis. Subsequently, enriched GO-BP terms with at least one test p-value < 0.005, were analyzed across the multivariate parameter space using PCA (Fig. 2A, B) and hierarchical clustering (Fig. S9A). In both cases, we used a matrix with the signed log-transformed enrichment p-values for the selected GO-BP terms for each parameter (see Data File S3). Hierarchical clustering was performed using Euclidean distances and ward.D2.

To generate an enrichment map (Fig. 2C and S9A), we performed the same GSEA over the −log-transformed HMP values that were zero-centered around −log(0.01). For mutants with multiple independent experimental measurements the median HMP was computed before the analysis. Here, all GO categories BP, MF (molecular function) and CC (cellular compartment) were included. GO terms with p-values < 0.05 and NES > 0 were graphed based on the Jaccard correlation coefficient (JC) for scoring the GO term similarities and a minimum JC > 0.01 for each edge, using the *enrichplot* (version 1.16.2) and *clusterProfiler* packages in R. The Spearman correlation over the profile count matrix for the GO terms was used to quantify the functional co-occurrence and purity of the different profiles in the enrichment map (Fig. 2D).

#### 5. Network analyses

##### 5.1 Spatial analysis of functional enrichment (SAFE)

SAFE was performed on the SGA-PCC network as published by Costanzo et al. 2016 (Fig. 3A)^29^. The SGA-PCC network was projected as in Costanzo et al. using the edge-weighted, spring embedded implementing Kamada-Kawai force-directed algorithm and loaded in Cytoscape (v.3.9.1)^84^. We also prepared a STRING network, based on a combined_score ≥ 990 of the STRING v10 database, and projected using the unweighted spring embedded layout algorithm (Fig. S10A, see Data File S4, *Costanzo and STRING node-edge*). Both networks were annotated with the GO-BP reference data from Costanzo et al. using SAFE^29,34^. The SGA-PCC network displayed a coverage of 84% and an attribute coverage of 56% and the STRING ≥ 990 network displayed a coverage of 99% and an attribute coverage of 66%. The enrichment landscape was built with a Jaccard correlation value of 0.75 and the resulting reference map labels were manually curated according to the functional domains identified. We also used SAFE to annotate the ATG core sub-graph using the ‘ATG set’ including *ATG1* and *ATG8*, and the fusion core sub-graph using the entire tethering complex (including *VTI1* and *YKT6*), the HOPS complex, and *VPS17*, *VPS29*, *VPS5*, *VPS35*, *VAM10*, *VPS1*, *VSP45*, *VPS3*, *VPS36* and *VPS26*. The profiles were enriched using SAFE and the enrichment colors were scaled with the SAFE score ranging from a min=0.35 to a max=0.35 (See Data File S4, *Costanzo and STRING SAFE*)

##### 5.2 Network distance analyses

The shortest network path from any gene to the ATG core machinery (ATG set) was computed by finding the set of genes one degree away from any gene within the ATG set, and iteratively finding the set of genes one degree away from any gene within the previous set, until all possible paths had been found (Fig. 3B). For the shortest causal network path, connector genes within the previous set were required to have a significant phenotype given by HMP < 0.01 or an autophagy perturbation p-value_min_ < 0.01.

##### 5.3 Gene-pair correlations

Correlations in phenotypes between genes were computed using the Pearson correlations of autophagy perturbation dynamics, *PCC*_Δ,{*m*1,*m*2}_ = *cor*(Δ%_*m*1,*t*_, Δ%_*m*2,*t*_), or equivalently using signed −log -transformed p-values over the set of kinetic parameter hypothesis tests performed. The correlations were subsequently grouped based on network interaction or complexome association status of the gene pairs as well as significance status (HMP ≤ 1, 0.01, or 0.001).

For strongly significant gene pairs (HMP ≤ 0.001) the correlations in perturbation dynamics were stratified into intervals for SGA-PCC concordance testing. The three intervals considered positive correlations (PCC_Δ_ > 0.25) representing similar phenotypes, negative correlations (PCC_Δ_ < −0.25) representing opposite phenotypes, and no correlations. Enrichment of SGA-PCCs was computed for each interval given different positive or negative SGA-PCC thresholds (0.1, 0.125, 0.15. 0.175, 0.2). Enrichment was calculated as 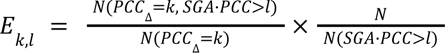, for a PCC_Δ_ interval *k* = (*lower*, *upper*] and a SGA-PCC threshold *l* (here indicated as greater than, but lesser than for − *l*). A Fisher’s exact test was used to compute the p-values for over- or underrepresentation.

##### 5.4 Directional GSEA along SGA-PCC spectra

Directional, positive or negative associations of autophagy perturbation profiles with SGA-PCC spectra for specific core machinery genes were computed using a modified GSEA algorithm (Fig. S11C). A positive or negative enrichment score (ES) was computed as:

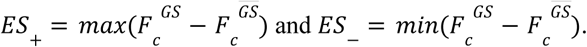

Here the *F* represent the cumulative distribution functions for the ranked SGA-PCC scores (x)

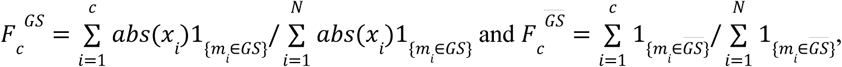

where 1 represent a indicator function for whether a mutant *m* is present in the perturbation profile gene set (*GS*) or among the non-gene set mutants (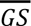). Direction preferential enrichment was computed as Δ*ES* = *ES* + *ES*. For statistical evaluation we used a permutation test where the distribution of *GS* membership along *x* were randomized 500 times to generate respective H0-distributions of *rES*, *rES*, and Δ*rES*, which were used to compute respective p-values, as well as the normalized enrichment scores 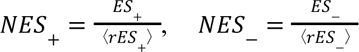, and Δ*NES* = *NES*_+_ − *NES*_-_. NES values were grouped for genes belonging to the ATG core and the fusion core machinery respectively, and profile enrichment per group was tested using a one-sample t-test.

##### 5.5 Module networks

The module networks were generated by identifying complexes through the Complex Portal^35^ or applying the MCODE algorithm^85^ to the STRING network (combined_score ≥ 990)^31^, with multiple significant genes. These complexes were then manually curated for protein-protein interactions using cross-references from SGD, STRING and BioGRID databases^30,31,86^. The networks were embedded in Cytoscape using the prefuse force-directed open layout algorithm and subsequently manually organized with node colors scaling with the signed −log_10_ p-values for the overall perturbation and size of the nodes scaling with their HMP (-log_10_) (see Data File S5).

#### 6. Latent-space and Bayes factors analysis

##### 6.1 Statistical analysis of the UMAP latent space

pUMAP predictions were performed on the latent autophagy feature vectors for all cells. The embedded cell densities were plotted by grouping populations per time points and color coding either for time or predicted autophagy % (*A*%). Moreover, the population means (µ*_U_*_1,*t*_, µ*_U_*_2,*t*_) and standard deviations (σ*_U_*_1,*t*_, σ*_U_*_2,*t*_) were computed for each UMAP coordinate (*x_U_*_1_ and *x_U_*_2_) per time point.

The UMAP embeddings were analyzed for representation of predictive information, population dynamics and noise. To test whether the single-cell UMAP embeddings conserved the information about autophagy from the DNNs, we trained a bag of regression trees to predict the DNN outputs, either represented as the probability of autophagy *P*_*a*_ or its log-odds, 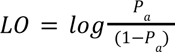. For this we used the *XGBoost* package (version 1.7.5.1) in R to train 50 parallel trees with subsampling = 0.63, depth = 10 and min_child_weight = 3^87^. The regression was performed per plate, where the data was split to 70% training data and 30% test data, and a PCC between the predicted and true values was reported. For comparison, the same procedure was used to test representation of time. We also compared these results with the ability of the UMAPs to separate WT from vam6Δ cells, by using bags of classification trees (with the same hyperparameter settings) subjected to 10-fold training and testing over the entire dataset, and scoring the concordance between the predicted class and the true label using an F1 score. To assess whether the information distinguishing these phenotypes were due to latent information about intermediate autophagic states in the UMAP or simply due to distributional differences based on the overall DNN prediction for the two strains, we stratified the UMAP distributions into percentiles based on the predicted autophagy *P*_1_ or *L0*, and then performed the same training and testing procedure within each interval.

To measure UMAP latent space noise and flux, we first calculated the population bivariate standard deviation, 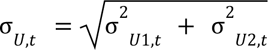, and the average population displacement, 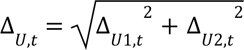, per time point. Here, the population displacement per coordinate, for instance Δ_*U*1,*t*_, is given as Δ_*U*1,*t*_ = µ_*U*1,*t*+1_ − µ_*U*1,*t*_. Δ_*U,t*_ and σ_*U,t*_ were compared with equivalent metrics computed from the unembedded latent feature vectors directly for populations of greater than 100 cells (Fig. S12D). For this comparison outliers for the latent space metrics beyond the range given by the quartiles Q1/Q3 -/+ 1.5 IQR (interquartile range) were filtered out.

As a measure of signal-to-noise we also computed the normalized flux 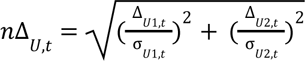 overall population variability and flux was assessed by averaging Δ_*U,t*_, *n*Δ_*U,t*_ and σ_*U,t*_ for each mutant time series per treatment phase. For this comparison only observations with an average of 50 cells were evaluated (see Data File S6).

##### 6.2 Bayes factor computation and assessment

Reference UMAP states of WT, *atg1*Δ, and *vam6*Δ cell populations were collected from 21 independent time series across 7 plates (Mix8, Mix10, and Rec1-5), where the WT cells comprised the dubious ORF negative controls in order for the latent space distributions to match that of unperturbed H_0_-outcomes of the KO collection. Kernel density estimators (KDE) were fitted using the *kde* function of the *ks* (Kernel Smoothing) package (version 1.13.2) in R (version 4.2.1)^88^. The KDEs were computed for the reference states based on their overall distributions independent of treatment phase (*K*_*WT*_, *K*_*atg*1Δ_, *K*_*vam*6Δ_), or their distributions stratified per time-point (*K*_*WT,t*_, *K*_*atg*1Δ,*t*_,*K*_*vam*6Δ,*t*_) in order to perform timewise phenotype comparisons. By using the *dkde* function we computed the probabilities *P*(*D*|*K*_*S*_), which represented the marginal likelihoods for UMAP observations *D* (U1, U2) given a reference state *S*. Thus, ratios of the marginal likelihoods comparing any two given states represented Bayes factors, *BF*(*WT*: *atg*1Δ), *BF*(*WT*: *vam*6Δ) and *BF*(*vam*6Δ: *atg*1Δ) that summarized the evidence favoring a given autophagic reference state in a specific UMAP (Fig. S14A,C). For time-independent phenotype classification of mutants between reference states *A* and *B*, the time-wise KDEs were paired with data for the corresponding time points to compute time-wise likelihoods. The individual KDEs were computed with equal matrix bandwidths generated by taking the medians of the bandwidth selection results using the *Hpi.diag* function on each time point. Subsequently, the log-likelihood ratios were averaged per mutant over all cells and all times points:

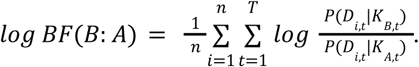

This population average or expected log Bayes factor can be viewed as a time-independent overall phenotype comparison per mutant, where dynamical effects have been ‘integrated’ out with the moving KDEs. This averaging was done for the entire experimental time course (overall), as well as for each treatment phase individually. On the other hand, to quantify the execution ‘activity’ as a timewise transition between temporally fixed reference states, the expected log-likelihood ratio was computed with the time-invariant KDEs over a cell population per time point:

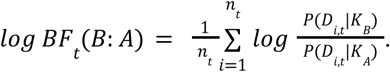

Within this Bayes factor framework, given the execution model {*A* → *B* → *C*} (*ATG*1 → *VAM*6: autophagosome formation; *VAM*6 → *WT*: autophagosome clearance), the overall activity for the transition {*A* → *C*} can be factorized into the intermediate transitions ({*A* → *B*} and {*B* → *C*}) as

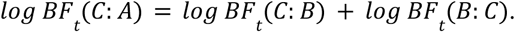

For comparison with the time-independent Bayes factors, and to score the expected activity over a given time frame, these timewise Bayes factors were averaged per mutant for all cells over a given time frame as well. This was done for the entire experimental time course (overall), as well as for each treatment phase individually (see Data File S7). Because these Bayes factors represent comparisons between fixed reference states, they are more sensitive to dynamical variation among the mutants, and hence indicated with the *t*-subscript. All time-averaged Bayes factors were corrected for distributional biases across plates by subtracting the plate-wise median difference from the screen median:

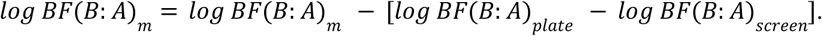

Analysis was restricted to mutants with at least n = 100 cells for time-averaged Bayes factors overall, and n = 50 cells when averaging per treatment phase. For mutants with multiple independent experimental measurements the median Bayes factors were computed. ROC curves and AUCs for analyzing retrieval of test gene sets (‘ATG set’ and ‘Fusion set’) were conducted as described above. Potential overfitting of the density kernels to the reference distributions was assessed by comparing the Bayes factors between *atg1*Δ and the ‘ATG set’, and between *vam6*Δ and the ‘Fusion set’. To compare the performance (ROC analysis) of the Bayes factors directly with the predicted autophagy perturbations (not subjected to sigmoidal curve fitting), Δ%_*m,t*_ = *A*%_*m,t*_ − *A*%_*ctr,t*_ and Δ_*binary*_ %_*m,t*_ = *A*_*binary*_ %_*m,t*_ − *A*_*binary*_ %_*ctr,t*_ were computed per mutant and averaged over the entire time series. Comparison with the fluorescent signal co-occurrence per experimental measurement was done by computing the mean co-occurrence early (0-1 hour) and late (> 7 hours) of the starvation protocol (Fig. S14A). In Fig. S17E, the overlap coefficient was computed by dividing the cell’s average co-occurrence by the average signal intensity of each channel multiplied (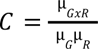), and computing the average across each cell population and time course. Functional analyses of Bayes factors were performed over the KO library only. We removed Bayes factors for experimental measurements where the mean red or green fluorescent signals early (0-1 hour of starvation) were below the 25th percentile - 1.5IQR of the GW screen distribution. For functional analyses we also computed the mean of the Bayes factor from the two neural latent space models (model 22 and model 30). To assess the concordance in phenotype classification between treatment phases, we paired the treatment phase specific Bayes factors and computed the fraction of deletion mutants with with substantial evidence for the same reference state (either log *BF* > 5 or log *BF* < –5) across both phases (Fig. S14C). Analysis of lagged correlations for time-wise Bayes factors (*log BF*_*t*_) were computed using the KO library only.

In Fig. 5C and S14E, where DAmP mutants were included, a harmonization procedure was performed where the Bayes factors were adjusted to the global mean and variance of the two libraries using the time-invariant Bayes factors:

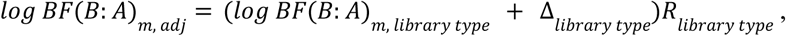

where *m* indicates the mutant, Δ_*type*_ = µ_*logBF*(*B*:*A*), *screen*_ − µ_*logBF*(*B*:*A*), *librarytype*_ is the difference in *screen* average from the *library* specific average (either KO or DAmP), *R*_*type*_ = σ_*logBF*(*B*:*A*), *screen*_ /σ_*logBF*(*B*:*A*), *librarytype*_ is the ratio of standard deviations for the *screen* over the specific *library* (either KO or DAmP).

##### 6.3 Parametric gene-set enrichment analysis

Parametric GO (BP and CC) enrichment analyses of Bayes factors were performed using the *piano* package (version 2.8.0) in R (version 4.2.1)^89^, using the gene sampling permutation test (nPerm = 10^5^) of the mean gene set statistics. The genesets were retrieved and annotated using the *GOdb* package (version 2.9) and org.Sc.sgd.db (version 3.15.0). Only the KO library phenotypes were analyzed, including the median of all *atg1*Δ and *vam6*Δ phenotypes observed. The Bayes factors or their respective treatment phase differentials were standardized over the GW population (to compute log-BF z-scores) prior to the analyses (see Data File S8, *z-scores*).

For analyzing the BF_t_ enrichment, the directional statistics (representing average log BF_t_ z-score per gene set) were visualized for gene sets with a p-value < 0.05 for at least one Bayes factor. For analyzing the BF differential enrichment, the directional statistics were visualized for gene sets with a differential enrichment p-value < 0.05. For the heatmap the gene-sets with a differential enrichment p-value < 0.01 for BF VAM6:ATG1 or BF WT:VAM6 and a minimum absolute statistic of one. Euclidean distance and ward.D2 were used for the clustering.

#### 7. Experimental validation of autophagy predictions

##### 7.1 Yeast and growth condition details

*S. cerevisiae* strains were grown at 30 °C until mid-log phase in standard yeast extract peptone dextrose (YPD) medium or in synthetic medium supplemented with relevant amino acids (SC) before proceeding with genetic manipulation and experimental assays. See *strain*s tab and *purpose* in Data File S1 for strains used in this section.

##### 7.2 Reproducibility library and strains construction for autophagy assays

All strains used in this section were derived from the S288c strain BY4741^90^ using either PCR-targeted gene replacement methods or crossing. Standard cell culture and genetic techniques were employed according to protocols^91,92^. Unless, otherwise noted, all genetic modifications were carried out chromosomally. Deletions and promoter replacements were performed as described^65^.

For creation of the “reproducibility” set, Atg8 was first N-terminal tagged by inserting mNG under control of the *ScTDH3* (GDP) promoter into the 5’ end of *ATG8*. Pep4 was c-terminal tagged with mCherry by inserting the mCherry coding sequence into the 3’ end of *PEP4* using the plasmid pBS35. Finally, all deletion strains and ORF control *YEL074W* were deleted using the *HIS3MX6* selection marker amplified from plasmid pFA6a-his3MX6 (See Data File S1; *strains* tab and *reproducibility* purpose).

For the quantitative Pho8Δ60 assay in Fig. S15A and S21B, the 60 N-terminal amino acid residues of the phosphatase Pho8 were removed by deleting 180 nucleotides from the 5’ end of *PHO8* using the plasmid PYM-N15, rendering expression of the truncated ORF under the control of the *ScTDH3* (GPD) promoter (See Data File S1; *strains* tab and *autophagy assays* purpose)^93,94^.

For the GFP-Atg8 processing assay in Fig. 6G and S15D, yeast cells were transformed with a plasmid expressing GFP-Atg8 under the control of the endogenous ATG8 promoter (pRS416 GFP-ATG8/AUT7; See Data File S1; *strains* tab and *autophagy assays* purpose)^95,96^. All strains were confirmed by PCR and fusion proteins were additionally validated by fluorescent microscopy and Western blot analysis. See Data File S1 for primer sequences.

For the prApe1 processing assay in Fig. 6K, cells were transformed with a CUP promoter-driven Ape1 plasmid (pCK782, a gift from Dr. Claudine Kraft). For Fig. S21, to visualize the overexpression of the Ape1 complex by fluorescent microscopy, cells harboring the pCK782 plasmid were modified to express mNG-Atg8 (see above) as well as Ape1-mCherry This was achieved by integrating an mCherry:hphMX6 PCR fragment, amplified from plasmid pBS35 using the primer pair APE1-S3 and APE1-S2, at the 3’ end of the *APE1* locus (see Data File S1 for strain details, and primer sequences).

##### 7.3 Growth and starvation protocols for screen reproducibility analysis

The screen reproducibility analysis comprised two approaches as described (See section Screen reproducibility analyses). In the first, aimed at replicating starvation observations in the initial screening, strains were grown sequentially to logarithmic phase, following an identical protocol as in the initial screening (See section 1.3-cell growth protocol for high-content screening).

In the second approach, focused on evaluating the uniformity of autophagy phenotypes across various experiments and strain backgrounds, the growth protocol was adjusted to induce autophagy with cells in stationary phase. Cells were grown in SC at 25 °C for three consecutive periods of 48 h each, followed by 8 h at 25 °C in the imaging plates, before being transferred to SD-N to induce nitrogen starvation for 12 h and replenishment for 7 h. Live imaging was performed at 30 °C, capturing images every hour for 19 h.

##### 7.4 Screen reproducibility analyses

###### 7.4.1 Nearest neighbor analysis and repetition set selection

Autophagy perturbation gene sets of different sizes were generated by setting various HMP cutoffs (≤ 0.01, 0.005, 0.001, or 0.0005; see Fig. S6B). Subsequently, we performed a Fisher’s exact test for overrepresentation of autophagy perturbations among the nearest neighbors using different GW yeast network data, including BioGrid PPIs, SGA-PCCs (top 1%), as well as all STRING interactions or only STRING-PPIs with a combined score > 500. These overrepresentation tests resulted in eight p-values, from which a harmonic mean was computed to yield an overall network enrichment significance (network p-value). Potential false positives (FPs) were defined as mutants with significant autophagy perturbations (perturbation FDR_min_ < 0.01; min referring to the lowest value between -N and +N), but non-significant network enrichment (network p-value > 0.1). Potential false negatives (FNs) were defined as mutants with non-significant autophagy perturbations (perturbation FDR_min_ > 0.1), and significant network enrichment (network p-value < 0.01). Potential FNs were also selected based on genes with key autophagy GO-term annotations (see Fig. S6C) and non-significant phenotype (FDR_min_ > 0.1 and HMP > 0.01). Top hits were selected by rank-ordering mutants based on the HMP (See Data File S9 and Data File S10).

###### 7.4.2 Replication of repetition set and outliers

Only mutants from the KO library were used for the validation screen, due to the lack of WT DAmP controls. This included 109 potential FNs, 56 potential FPs, and 75 top hits (See Data File S1, *repetition library* and Data File S10, *repetition responses*). For the validation screen perturbations in predicted autophagy (Δ%_*m*, *t*_) were computed from the mean WT ORFs included in each plate (since plate medians would not be valid WT references due to the enrichment of mutant phenotypes). The replicability was tested by repeating the GW screen protocol up to 8 hours of starvation, and computing the correlation in predicted autophagy dynamics (*PCC*_*A,m*_ = *cor*(*A*%_*m,t,R*1_, *A*%_*m,t,R*2_)) and autophagy perturbation dynamics (*PCC*_Δ,*m*_ = *cor*(Δ%_*m,t,R*1_, Δ%_*m,t,R*2_)) for each mutant between replicates *R*. The dynamics correlations were compared with the mean absolute autophagy perturbations, 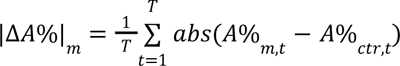. For mutants with multiple independent experimental measurements in the GW screen the median responses were computed before the comparisons.

###### 7.4.3 Reproducibility across experiments, strain background and conditions

To assess the reproducibility of autophagy phenotypes across strains-background, independently generated clones, and growth conditions, 33 mutants covering different cellular functions and with different autophagy dynamic profiles were picked manually. All mutants were strongly significant (HMP ≤ 0.001). Three independently generated new clones were included for each mutant, along with the collection clone, and three new screens were done independently (See Data File S10, *validation responses*; see Fig. S7A-B).

For evaluation of mutant replication error, the mean absolute perturbations were computed per mutant and compared with the mean absolute errors, 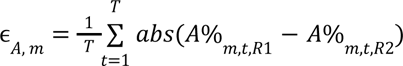, between different types of replicates *R*. These included technical replicates (same mutant clone, different wells within the same plate), experimental replicates (same mutant clone, different plates/experiments), replication of mutant phenotypes generated in a new background strain, and replication between independent clones generated within the same background strain. Correlations in autophagy dynamics (*PCC*_*A*_) and autophagy perturbation dynamics (*PCC*_Δ_) were also computed per mutant across experimental replicates and between collection clones and clones generated in a new background strain. Mutant perturbations in autophagy on the new screen plates were computed with respect to the average from 12 independent WT controls, 6 based on the collection background strain and 6 based on the new background strain.

For assessing screen-level experimental reproducibility of autophagy perturbations and kinetic parameters, Pearson and Spearman correlations (due to large outliers for some parameters) were computed from global pairwise comparisons of the screen metrics across all mutant clones. Comparisons were done between the GW screen and the new screens for autophagy perturbations, Δ%_*t*_, grouped on treatment phase (-N, +N, or both) and strain background (collection clones, new clones, or both), or between the new screens across all clones for the parameters (See Data File S10, *validation statistics*).

##### 7.5 Rapamycin-dependent autophagy response and analysis

###### 7.5.1 Experimental cell growth and conditions

Cells were grown in SC at 25 °C for two consecutive periods of 48 h each, followed by a period of 24 h and a last for 8h at 25 °C in the imaging plates. Cells were then washed twice with sterile MilliQ water and transferred to independent plates containing 200ul of SC+Vehicle, SC+Rapamycin, SD-N+Vehicle and SD-N+Rapamycin. Live imaging was performed at 30 °C, capturing images every hour for 10 h. Rapamycin was dissolved in ethanol and Triton-X in a 9:1 ratio in a concentration of 1 mg/ml. The final concentration of rapamycin used was 440 nM. As a vehicle control, the solvent of rapamycin was added in the same proportions. Each plate included duplicate experimental conditions of every strain (triplicates for the controls), and the complete experiment was repeated three times.

###### 7.5.2 Data Analysis

Autophagy (*A*%_*t*_) and UMAP Bayes factors (*BF*_*t*_) were computed for each well (see Data File S11). Autophagy perturbations (Δ%_*t*_) were computed with respect to the mean of ORF plate controls, and for all metrics the time series averages were computed. Reproducibility between experiments was assessed by computing PCC for each metric.

##### 7.6 Yeast growth conditions for protein and RNA analysis

To correspond to library growth conditions, cells were serially diluted in a fresh medium to maintain them in logarithmic phase before inducing autophagy.

For protein analysis of the GFP cleavage assay, cells were first grown overnight in SC-URA and transferred to grow in three sequential cultures in SC-URA at 30 °C to mid log-phase for 24 hours. Cells were then washed five times with nitrogen-starvation medium (SD-N: 1.9 g/ml yeast nitrogen base without amino acids and ammonium sulfate and 20 g/L glucose) using membrane filtration and transferred to SD-N at an OD_600_ = 0.25, followed by incubation for 4 or 8 h at 30 °C according to the experiment. In the experiments with prolonged starvation (8 h), cells were shifted to SC-URA medium and further incubated for 4 h. Cells were collected at specified time points during nitrogen starvation and replenishment and subjected to protein analysis.

For protein analysis of the pr-Ape1 processing assay, cells were grown in SC-LEU at 30°C to mid-log phase for 24 hours. Cultures were then split into two conditions: one supplemented with 50 µM CuSO₄ to induce Ape1 overexpression for 20 hours, and a control condition without CuSO₄. To induce starvation, cells were washed five times with SD-N using membrane filtration and resuspended in SD-N at an OD₆₀₀ = 0.25. Cells were then incubated at 30 °C for 4 hours in SD-N, and samples were collected at the indicated time points according to the experiment.

For RNA analysis, cells were sequentially grown in SC at 30 °C to mid-log phase, transferred to SD-N, and collected at specified time points during nitrogen starvation and replenishment and subjected to protein analysis. Samples at t=0 were collected from a portion of the culture that was shifted to fresh SC once more and incubated at 30°C for an additional hour.

##### 7.7 GFP-Atg8 and pr-Ape1 processing assay

The GFP-Atg8 and pr-Ape1 processing assays were performed and interpreted as described^64,96,97^. According to each experiment, cells were collected at indicated times and protein extracts were prepared and resolved by SDS-PAGE. Full-length GFP-Atg8 and free GFP were detected by Western blot using an anti-GFP antibody and pr-Ape1 processing using an yeast anti-Ape1 antibody.

##### 7.8 Protein extraction, immunoblotting and antibodies

Cell lysis and immunoblotting were performed as previously described^98^. For protein analysis, 2 x 10^8^ cells were collected by filtration, washed with STOP buffer (150 mM NaCl, 50 mM NaF, 10 mM EDTA, 1 mM sodium azide) and frozen on dry ice before extraction. Total protein extracts were prepared by precipitation with trichloroacetic acid (TCA) as previously described ^99^. Following gel electrophoresis in 4–20% Criterion™ TGX™ Precast Midi (BioRad), proteins were transferred onto a PVDF membrane using a semi-dry blotting system (Trans-Blot® Turbo™ from BioRad). The membrane was blocked in 5% skimmed milk with TBS-0.1% Tween and incubated with primary antibodies in 5% BSA with TBS-0.1% Tween (for antibody against phosphorylated residues) or 5% milk with TBS-0.1% Tween (in all other instances). Western blots were developed using Pierce™ and SuperSignal® Dura ECL detection reagents (Thermo Fisher) and imaged using Chemidoc MP Imaging system (Bio-Rad laboratories). The following primary antibodies were used in this study: rabbit anti-phospho-(Ser/Thr) Akt Substrate Antibody to detect phosphorylated RpS6 (1:1000, Cell Signaling Technology 9611), goat anti-GFP (1:1000, Abcam 5440) to detect GFP-Atg8, rabbit yeast anti-Ape1 (1:10000, a gift from Dr. Claudine Kraft)^54^ and mouse anti-beta Actin (1:1000, Abcam 8224). As secondaries, HRP-conjugated Goat anti rabbit IgG (1: 5000 Sigma), HRP-conjugated Bovine anti goat IgG (1:5000 JIR) and HRP-conjugated Goat anti mouse IgG (1:10000 JIR). The Western blot experiments were performed in replicates as indicated in Data File S12 and representative images were chosen for publishing.

##### 7.9 Quantifications of Western blot with Bio-Rad Lab software

The degradation of GFP-fused Atg8 in the vacuole, the processing of pr-Ape1 in the vacuole, and the phosphorylation of RpS6 in relation to actin as loading control were estimated by calculating the densitometric values of the bands in the raw digital WB images using Image Lab software from Bio-Rad Laboratories 6.0.1. The WB experiments were performed for multiple independent replicates as indicated in Data File S12 (*see GFP quant., Ape1 quant. and Rps6 quant.*). See Data File S12 for all quantifications and numerical transformations.

##### 7.10 Autophagy flux by assessing GFP cleavage and Rps6-p comparison with BFs

WB quantifications were averaged over two exposure times for every experiment before computing the mean and SEM from the independent biological replicates (see Data File S12, *Bayes Factor comparison*). GFP cleavage was computed from the adjusted band volume (intensity) as the percentage of free GFP signal divided by the total GFP signal (free GFP + GFP-Atg8) within each lane using densitometric analysis (Bio-Rad Image Lab Software). The relative RpS6-phosphorylation (RpS6-p) was computed from the absolute band volume by dividing the RpS6-p signal by the β-actin signal and normalizing each experiment to their respective WT T0 value.

The time-averaged BFs, kinetic parameters and average screen metrics were compared with the autophagic flux by computing the mean GFP cleavage from 8 hours of starvation (0, 2, 4, 6 and 8 hours). The similarity in autophagy phenotypes measured across mutants were assessed using the Pearson correlation. For time-series comparison of BF_t_ with GFP cleavage, we matched the time-wise measurements for -N (0, 2, 4, 6 and 8 hours) and +N (0, 1, and 4 hours) for the two time series experiments (see Data Table S12, *GFP quant.* for number of replicates per mutants and conditions). Note that +N for the BF_t_ occurred from 12 hours in -N, while for GFP cleavage +N occurred from 8 hours in -N. Moreover, the 5 min time point in +N for GFP cleavage were used to match with T0 replenishment for BF_t_. The time-wise similarity were then assessed using a mixed model based on BF_t_ WT:ATG1 alone (model 1) or BF_t_ WT:VAM6 and BF_t_ VAM6:ATG1 (model 2), with random intercept and slope for each time point:

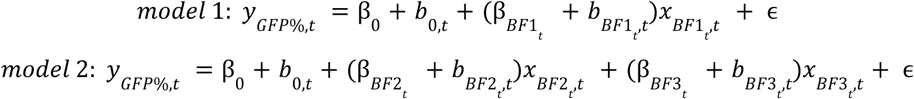

Here, β represent fixed effects, *b* represent random effects per time point *t*, ɛ is the residual error, *x*_*BF*1_*t*__ represent log BF WT:ATG1 values, *x*_*BF*2_*t*__ represent log BF WT:VAM6 values, and *x*_*BF*3_*t*__ represent log BF_t_ VAM6:ATG1 values. Goodness-of-fit was assessed by measuring the Pearson correlations between predicted and observed values across mutants for each time point and across time points for each mutant. The time-wise variable (BF_t_) contribution to autophagic flux point was computed as (β*_BF_t__*+ *b_BF_t_,t_*)σ*_BF_t_,t_*, where σ*_BF_t_,t_* is the standard deviation of the log BF_t_ sample space at time point *t*.

For comparing with the Bayes factors with the RpS6-p status, the time-averaged BF_t_s were correlated with RpS6-p at T0, average RpS6-p in -N (2, 4, 6 and 8 hours), and average RpS6-p in +N (15 and 30 min, 1, 2 and 4 hours) (see Data Table S12, *Rps6 quant.* for number of replicates per mutants and conditions).

Statistical evaluation of genetic interactions effects on GFP cleavage between *rtg1*Δ and *gcn4*Δ or *gln3*Δ, were done using a mixed effects model with random intercept on experiment identity, *rtg1*Δ as baseline and interaction between the mutants and time (see Data File S12, *GFP stats._RTGs*).

##### 7.11 Quantitative Pho8Δ60 assay

Pho8Δ60 assay was performed as previously described^94,100^. For the analysis of alkaline phosphatase (ALP), cells were first grown to mid-log phase (5 x 10^6^ cells/ml) in SC at 30°C and then diluted to 2.5 x 10^6^ cells/ml and shifted to SC for 1h and SD-N for 4h. For sampling, 1 to 1.25 x 10^6^ cells were collected by filtration, washed with 1 ml of PBS pH 7.2 and pellets were frozen on dry ice. For protein extraction, cell pellets were dissolved in 500 µl of lysis buffer at pH 6.8 containing 1 M PIPES, 10% Triton-100X, 1 M KCl, 1M KOAc, 1M MgS0_4_, 10 mM ZnSo4 and 2mM of PMSF. Cell disruption was performed by agitation and homogenization with a cryogenic bead beating grinder (FastPrep®-24 5G). For the phosphatase assay, 0.1-0.5 mg of total protein was used as the phosphatase source, with 125 mM p-nitrophenyl phosphate serving as the substrate in the reaction buffer. The reaction buffer was prepared with 1M Tris-HCl at pH 8.6, 10% Triton-100X, 1M MgSO_4_, and 10 mM ZnSO_4_. The mixture was incubated at 37°C for 5 and 7 min, allowing the phosphatase present in the total protein to dephosphorylate the p-nitrophenyl phosphate substrate. The enzymatic reaction was then stopped by the addition of 1M glycine at pH 11. The ALP activity was calculated as the amount of p-nitrophenol (nmol) produced per minute per milligram of total protein. Both incubation times were measured using independent reactions. The Pho8Δ60 assay experiments were performed for multiple independent replicates as indicated in Data File S12 (*see pho8d60 measures*). Data was presented as mean ± SEM. Significant differences were calculated using a one-way ANOVA with a TukeyHSD post-hoc test. Significance is reported for mutant comparisons with the negative control within each treatment phase (see Data File S12, *pho8d60 stats*).

##### 7.12 RNA extraction and quantitative PCR (qPCR)

RNA extraction and quantitative PCR (qPCR) were performed as previously described^98^. For qPCR analysis, 8 x 10^7^ cells were collected by centrifugation, washed in DEPC-treated water and frozen in dry ice. Total RNA preparation was performed with MasterPure™ Yeast RNA Purification Kit (Epicentre) following the manufacturer instructions. 1 µg of RNA was used for cDNA synthesis using SuperScript® III Reverse Transcriptase (Invitrogen). qPCR was performed with SYBR® Select Master Mix (Applied Biosystems). Primer sequences (fwd and rv) are listed in Data File S1. RNA analysis experiments were performed in triplicate as indicated in Data File S13 (see *input data*).

##### 7.13 qPCR analysis

Three independent experiments were performed, each with two technical replicates for all targets per treatment condition. The mean C_t_ value was calculated for each target per treatment condition and subsequently normalized by subtracting the median value of the actin control to compute a ΔC_t_ value, which was transformed to the signal *S* = 2^-Δ*Ct*^. The complete dataset was batch-corrected with the *ComBat* algorithm from the *sva* package (version 3.44.0) in R^101^, using parametric adjustments with experimental replicate as batch variable and genotype as model variable. *S_adj_* was log transformed before the batch correction, and exponentiated after to generate *S_adj_*. Finally, *S_adj_* was normalized to the mean *S_adj_* of the WT controls at T0 and log -transformed (See Data File S13, *input data*).

Significance for individual time points was assessed by comparing the mutants with the WT using a two-tailed t-test. For statistical evaluation of the overall differential regulation effect by the mutants on every target, a regression was performed per treatment phase (T0 +N, -N, and +N), with WT as baseline and genotype and time point as independent variables. The reported values are the coefficients for the genotype effects (See Data File S13, *statistics*)

##### 7.14 Live-cell imaging of autophagosome dynamics and analysis

###### 7.14.1 Experimental cell growth and imaging

Cells were grown sequentially following the cell growth protocol for high-content screening (See section 1.3). For time-lapse imaging, 4-6 µl of cells were inoculated into 96-well plates (Nunc™ Microwell™, Thermo Fisher) containing 200 µl of SC per well and incubated for 8 hours at 25 °C to ensure cells were in exponential phase. To induce autophagy, 400 µl of cells were inoculated into ibidi µ-Slide 8-well untreated plates (Ibidi®) coated with Con A (0.25 mg/ml) and washed twice with 400 µl of SC medium and shifted to 400 µl of SD-N medium for 2 h. Live imaging was performed using a Nikon ECLIPSE Ti2-E inverted microscope equipped with a temperature control unit, CrestOptics X-Light V3 spinning disk confocal module and Lumencor Celesta multi-line laser (Nikon Crest System). The images were acquired using automated acquisition in wide-field, utilizing the CFI Plan Apo l D 100X Oil NA 1.45 (Oil) coupled to a 16-bit back-illuminated sCMO camera and Z-drive and PFS autofocusing mechanisms. Image acquisition was controlled by the NIS Elements AR software (Nikon Instruments Inc.). Live cell imaging was conducted at 30 °C ± 2 °C. Fluorescence signals were captured using specific filter sets for WF GFP and WF RFP, with exposure times of 30 ms and 100 ms, respectively, using z-sections acquired at 0.2 µm steps over a range of 0.8 µm. For bright field images, an exposure time of 100 ms was used to capture a single focal plane. For quantification and tracking of autophagosomes in WT and TOR signaling mutants (Fig. 5E-G and Fig. S19), cells were imaged every 5 minutes for 2 hours in SD-N, then shifted to SD for an additional 2 hours, capturing one image per well. Experiments were conducted in four replicates using mutants from the yeast mutant libraries harboring the autophagy reporter mNeonGreen-Atg8 and Pep4-mCherry (Open Biosystems). For quantification and tracking of autophagosomes in WT, *rtg1*Δ, *vam3*Δ and DKO cells (Fig. 6H-J and Fig. S21B), cells were imaged every 15 minutes for 2 hours, capturing 3–6 images per well. Experiments were conducted in three replicates, each with two independently generated clones harboring the autophagy reporter mNeonGreen-Atg8 and Pep4-mCherry.

###### 7.14.2 Cell and vesicle segmentation and colocalization analysis

We performed image analysis using a custom FIJI/imageJ pipeline to quantify autophagy dynamics, including the formation of autophagosomes, represented by changes in the intensity and distribution of the green signal, and fusion rates, defined as the colocalization between green and red signals. We categorized the images by position (XY), channel (BF, GFP and RFP), focal plane (Z) and time point (t). We generated maximum intensity z-projections for each position, channel, and time point. We then assembled sequential time points per channel into a time-lapse stack, followed by image processing steps, including applying an unsharp mask filter (radius = 1.0, weight = 0.60) to enhance edges and performing background subtraction (rolling radius = 50 pixels) to reduce noise. For optimal visualization, we applied alignment corrections across time points using SIFT algorithm (rigid, 2048 pixels) for each channel and we merged the GFP and RFP time-series stacks into a single RGB composite image.

For cell segmentation, we applied the ‘Huang Dark’ threshold to convert the non-aligned processed t-stacks into binary masks. We then used the Analyze Particles plugin with a minimum particle size of 20 pixels and a circularity range of 0.1–1.00 to extract measurements, including *Area, Perimeter, Centroid, Mean, Median, Bounding box, Feret’s diameter, Shape, Integrated Intensity, Skewness, Kurtosis, and Stack position*. For autophagosome segmentation, we applied the ‘MaxEntropy’ threshold and used the Analyze Particles plugin with a minimum particle size of 1 pixel and a circularity range of 0.0–1.00, restricting the analysis to autophagosomes within the segmented cell mask. The same measurements used for cell segmentation were applied to vesicle tracking over time. Last, we quantified colocalization by multiplying the pixel values of the GFP and RFP channels using the ‘Multiply create 32-bit’ command.

###### 7.14.2 Data analysis

Cells were filtered for an area between *1400* and *6000 pixels* and *roundness* > *0.7*; vesicles were filtered for an area below *750 pixels*. To track cells, a random forest classifier was used to evaluate the probability of candidate cells within a minimal radius in the following image frame being the same as a cell in the preceding image frame. The classifier was trained to distinguish the absolute distance in different cell features for the closest cell pairs between image frames in the dataset over randomly sampled cell pairs. The features included signal features (*Mean, StdDev, Min, Max, IntDen, Median, Skew, Kurt, RawIntDen*) for each channel and the co-occurrence channel, and shape features (*Area, Perim., Width, Height, Major, Minor, Angle, Circ., Feret, FeretAngle, MinFeret, AR, Round, Solidity*). The cell pair with the highest probability over a *p* > *0.5* threshold was assigned a common trace ID. Equivalently, to track autophagic vesicles within cell traces, a random forest classifier was used to evaluate vesicle pairs between time frames. Here, the differences in vesicle features (signal features and shape features) between following and preceding images were used, along with experimental time and differences in relative positional information (*X, Y, BY, BY, FeretX, FeretY*) of a vesicle within a cell. The classifier was trained to distinguish between the closest vesicle pairs between image frames over randomly sampled vesicles pairs. The vesicle pair with the highest probability over a *p* > *0.5* threshold was assigned a common trace ID.

Autophagosome fusion with the vacuole was quantified using a vesicle-specific overlap coefficient: 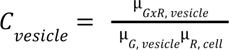, where µ_*GxR*, *vesicle*_ is the average co-occurrence per puncta, normalized by the average puncta intensity for Atg8/green channel, µ_*G*, *vesicle*_, and the average intensity of the Pep4/red channel per cell, µ_*R*, *cell*_. This metric quantifies the Atg8 vesicle-specific enrichment of the vacuolar marker Pep4. To estimate the vesicle fusion rates, we fit the mixed model with random intercept on vesicle ID:

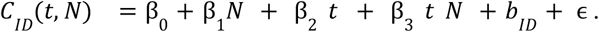

Here, t indicates the time and N indicates the addition of nitrogen. β_0_ is the baseline, β_0_ is the change in baseline after 2 hours of -N and addition of nitrogen, β_2_ is the slope, and β_3_ is the slope under addition of nitrogen. For this analysis, vesicles without a trace between time points or vesicles already present in newly identified cell traces were filtered out.

##### 7.15 Live-cell imaging of overexpressed Ape1 aggregates

Cells expressing mNeonGreen-Atg8 and prApe1-mCherry, along with overexpressing of prApe1 under a copper-inducible promoter (pCK782, see section 7.2) were grown to logarithmic phase. For time-lapse imaging, 1–4 µl of cells were inoculated into 96-well plates (Nunc™ Microwell™, Thermo Fisher) containing 200 µl of SC per well and incubated at 25 °C for 20 hours with either 50 µM CuSO₄ to induce prApe1 overexpression or 0 µM CuSO₄ as a control. To induce autophagy, 400 µl of cells were transferred to ibidi µ-Slide 8-well untreated plates (Ibidi®) coated with Con A (0.25 mg/ml), washed twice with 400 µl of SC medium to remove CuSO₄, and then shifted to SD-N medium for 2 hours. Live imaging was performed using a Nikon ECLIPSE Ti2-E inverted microscope equipped with a temperature control unit and a CSU-W1 spinning disk confocal unit (Nikon SoRa System). Images were acquired using automated wide-field acquisition with the SR HP Apo 100X Oil NA 1.45 objective, coupled to a 12-bit back-illuminated sCMOS camera, with Z-drive and PFS autofocusing mechanisms. Image acquisition was controlled by NIS Elements AR software (Nikon Instruments Inc.). Live-cell imaging was conducted at 30 °C ± 2 °C. Fluorescence signals were captured using specific filter sets for WF GFP and WF RFP, with exposure times of 30 ms and 100 ms, respectively, using z-sections acquired at 0.2 µm steps over a range of 0.8 µm. Bright-field images were captured at a single focal plane with a 100 ms exposure time. To track the ability of cells to capture Ape1 aggregates into autophagosomes, images were acquired every 90 seconds for 900 seconds. Experiments were conducted in two replicates using two independently generated clones, and images were processed and analyzed with FIJI/ImageJ software.

##### 7.16 Electron microscopy

For harvesting, cells were first grown in YPD and transferred to grow in three sequential cultures in SC at 30 °C to mid log-phase for 24 hours. Cells were then washed five times with SD-N using membrane filtration and transferred to SD-N at an OD_600_ = 0.25. Cells were then incubated for 1h, washed with 1 ml of PBS pH 7.2 and resuspended in 30 μl of the remaining buffer. Cells were then immediately high pressure frozen using a Leica HPM 100. Freeze substitution was performed as follows: cryo-vials were filled with freeze substituent (acetone containing 1% (w/v) osmium tetroxide, 0.1 % (w/v) uranyl acetate, 0.25 % glutaraldehyde, 1% water) and placed in a temperature controlling AFS2 (Leica) and cooled down to −90 °C. Sample carriers containing yeast cells were transferred into cold cryo-vials and left at −90 °C for 48 h before the temperature was raised to −45 °C, over a time span of 9 h. The samples were kept at −45 °C for 5 h before being raised stepwise stepwise to −20 °C (within 1 h) and then +4 °C (within 30 min). The samples were removed from the AFS2 once the temperature had reached +4 °C, and samples were washed 3 times in acetone and then infiltrated with increasing concentrations of epon (25 %, 50 %, 75 %, 2 x 100 %, 1 h each) mixed with acetone. Finally, cells were infiltrated a third time with epon (100 %) and left O/N for infiltration at room temperature. Polymerization was conducted at 60 °C for 72 h. Serial sections (50 nm thickness) were cut on an Ultracut UCT ultramicrotome (Leica, Germany) and collected on formvar-coated slot grids and poststained with Reynold’s lead citrate and 4% uranyl acetate solution. Electron micrographs of individual yeast cells were collected with a Thermo ScientificTM TalosTM F200C microscope and vacuolar inclusions that were not continuous with the cytoplasm were annotated manually and measured and analyzed with FIJI/ImageJ software. A one-way ANOVA comparing WT (n= 93) and *rtg1*Δ (n=91) inclusion body areas was concluded as non-significant.

#### 8. Cross-omics inference for autophagy predictions

##### 8.1 Cross-omics prediction benchmarking

Cross-omics modeling and prediction benchmarking performed against the time-averaged BFs and kinetic parameter statistics (signed −log_10_-transformed p-values) for the KO library only. Three omics datasets from a genome-wide deletion library were tested, as well as the SGA-PCC dataset and a STRING protein similarity network (STRING-PCC). The individual datasets were matched for overlapping deletion mutants with autophagy phenotypes as the output and subsequently tested with random forest and 10-fold cross-validation using all features as model inputs.

The proteomics dataset was retrieved from Messner et al. 2023^50^, and we used the complete imputed dataset. Each protein abundance value was normalized by the median value of the WT controls and subsequently log_2_-transformed. The dataset overlapped with 4189 deletion mutants and included 1850 proteins as model features.

The transcriptomic dataset was retrieved from Kemmeren et al. 2014^49^, and represented microarray-based log_2_-fold changes from the WT controls. The dataset overlapped with 1419 deletion mutants and included 6112 transcripts as model features. The expression profiles for each mutant were standardized to have zero mean and unit variance.

The metabolomics dataset was retrieved from Mülleder et al. 2016^51^, and represented concentration changes in deletion mutants for 19 amino acids. The dataset overlapped with 4501 deletion mutants.

The SGA-PCC was retrieved from Costanzo et al. 2016^29^. Because this dataset includes some correlation with multiple deletions for the same gene, we averaged the correlations per ORF over the columns and rows, subsequently. The resulting dataset overlapped with 4565 deletion mutants and included 5707 gene correlations as model features.

The STRING-PCC dataset was retrieved from the STRING database version 10^31^. A binary interactome matrix for combined_sore ≥ 200 was generated and subsequently used to compute pairwise PCC between each column. The resulting correlation matrix overlapped with 4684 deletion mutants and had 6385 gene correlations as model features.

The response variables were standardized before training. Random forest was executed using the regression implementation of the *XGBoost* package (version 1.7.5.1) in R (version 4.2.1)^87^ using squared error as loss and learning_rate = 1. Number of parallel trees were set to 1000, subsample = 0.63 and colsample_bynode = 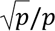, where *p* indicated the number of features and varied per dataset. The number of splits were set to max_depth = 20, and we used a min_child_weight = 3. The r^2^ was computed for each test fold and the median value was reported.

##### 8.2 Random forest mediation analysis

Discovery of transcriptional mediators of regulatory effects of transcription factor (TF) on autophagy responses where executed by matching the time series of the Induction Dynamics gene Expression Atlas (IDEA) by Hackett et al. 2020^52^ with the initial starvation responses dynamics of 186 overlapping TFs, using a 1 hour time shift. The rationale for using the IDEA dataset, is that the gene expression profiles presented in IDEA are directly causally driven by the acute activation of the TFs. Moreover, because of the abundance of autophagy perturbations caused by TF deletions, and the relatively low variation in autophagy at the onset of the starvation protocol, it is likely that a lot of the dynamic variation is related to loss-of-functions of crucial gene expression changes acutely involved in the starvation response. Gene target induction profiles that strongly predict the autophagy perturbation responses generated across multiple TF deletions, where the targets themselves are genes with a measured influence on autophagy, represent probable candidates as targets for transcriptional modulation of autophagy.

In the IDEA datasets we used ‘log2_cleaned_ratio’ variable for differential gene expression time series (*X*), which are normalized to the T0 baseline of each induction protocol. To compensate for varied efficacies on gene expression regulation from induction for different TFs, as well as batch effects, we performed the harmonization procedure *X*_*TF,adj*_ = *X*· _*TF*_(σ_*X*_ /σ_*X_TF_*_), where σ*_X_* is the standard deviation of *X* for the entire screen and σ_*X_TF_*_ is the standard deviation of *X*_*TF*_ for a specific TF. The reported log fold change (log FC) values in the study are based on these adjusted values. Missing values in the time series were imputed using the na_interpolation function with the stine method from the *imputeTS* package (version 3.3) in R^102^, and time series values from 30, 45, 60 and 90 minutes after TF induction were extracted for the remaining analysis. These were matched with 90, 105, 120 and 150 minutes after starvation for *A*%_*t*_ and the *BF*_*t*_ time series, where the intermediate missing values were imputed using the na_interpolation function with the spline method. Thus, the resulting dataset had 744 samples (186 TFs x 4 time points). Before the regression the response variables were centered around the median of each time point, to compute the perturbations, and then standardized. The dataset had 6175 gene targets as features, 5989 after removal of the 186 TFs, and 4508 that were present in the autophagy dataset, where 1354 targets also had a significant autophagy phenotype (HMP ≤ 0.01).

Random forest was executed as described above using the *XGBoost* package (version 1.7.5.1) in R (version 4.2.1), with number of parallel trees set to 5000. For model evaluation 10-fold cross-validation was performed, where each fold was defined by withholding a random sample of individual data points or a random sample of TF time series. The r^2^ was computed for each test fold and the median value was reported. To score the feature importance we used the sumGain, which was computed using the importance function in the *XGBoost* package. For clustering selected TFs with potential mediators, target genes with a sumGain > 2200 against one of the response variables and a HMP < 0.001, were considered (See Data File S14, *importance scores*). The sumGain cutoff was estimated as 2 times the average sumGain of features from control models trained with randomized sample IDs. SHAP (SHapley Additive exPlanations) values were used to assess the predictive effect from gene expression changes on autophagy perturbations. To compute SHAP values we used the SHAPforxgboost package (version 0.1.3)^103^. The individual SHAP values were grouped on whether the corresponding rfvalue (raw feature value; corresponding to the log fold changes in expression, *X*_*TF*_. *adj*) were positive or negative, and then computing the mean per group for each feature (see Data File S14, *SHAP scores*). Thus, this yielded average SHAP values for upregulated and downregulated gene expression, respectively. Differential SHAP values were computed by subtracting the SHAP value for upregulation with the SHAP value for downregulation. To compare the SHAP values for models trained with different number of features they were multiplied with *p*/*p*_*max*_, where *p* indicate the number of features for the specific model and *p*_*max*_ indicate the maximum number of features.

---

**Figure S1.**
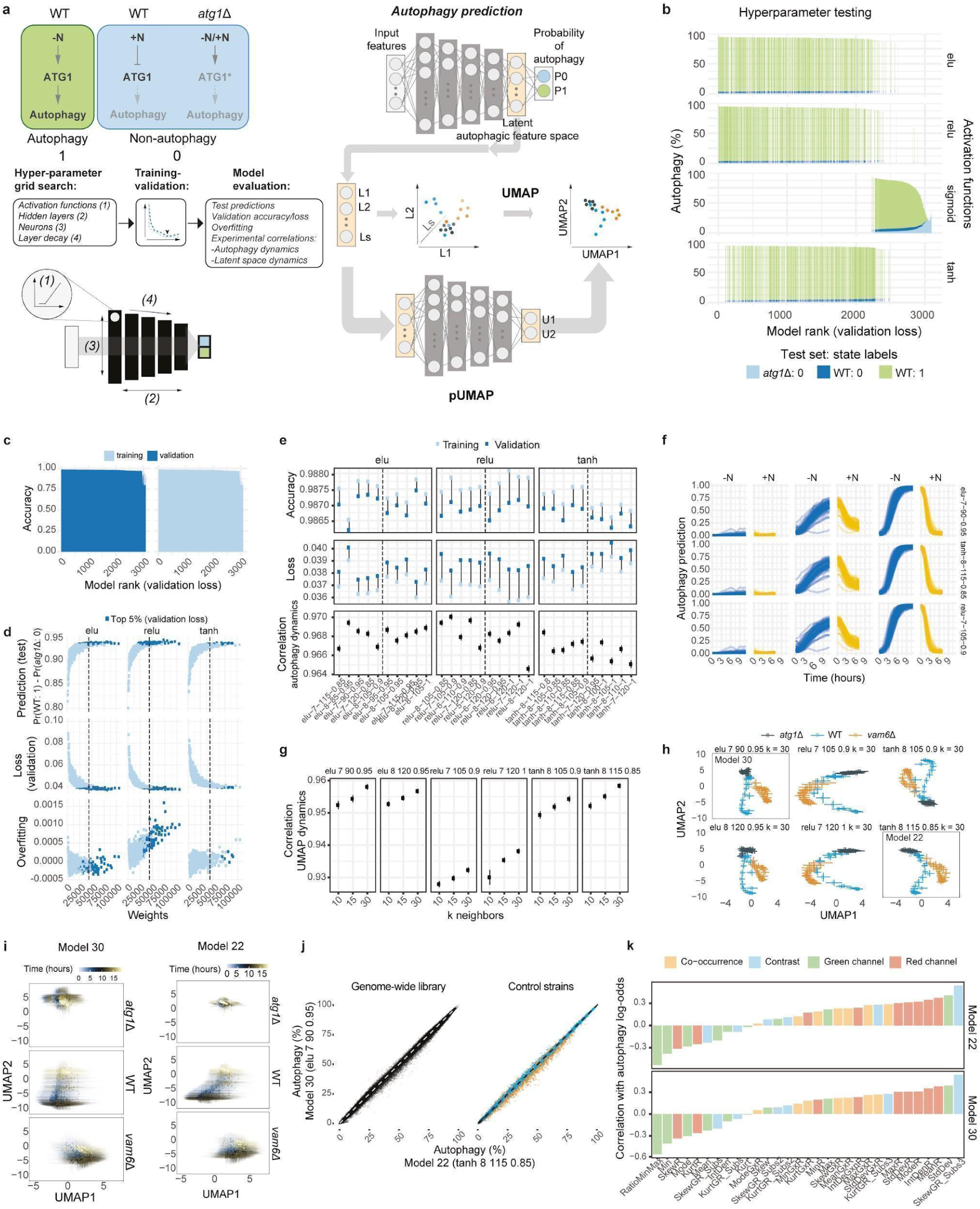
Robust quantification of autophagy using deep learning. **a**, Overview of deep learning approach. *Top left*: Causal control conditions for training data. Non-autophagic states (0) were defined as WT cells at T0of starvation and after 4 hours of replenishment and as *atg1*Δ cells for any time points. Autophagic states (1) were defined as WT cells after 7 hours of starvation until T0 replenishment. *Top right*: Neural nets were trained on extracted single-cell signal distribution features to classify autophagic cells. *Bottom left*: A hyperparameter grid search was initially performed over different neural net models with varying activation functions, hidden layers, number of initial neurons, and decay in the number of neurons per layer. The initial collection of models was trained for 150epochs with early stoppage on validation loss and subsequently evaluated for test predictions, validation accuracy loss, and overfitting. Top models with an *elu*, *relu*, or *tanh* architecture were selected based on the validation loss and retrained for 3000 epochs with early stoppage on validation loss. *Bottom right*: A UMAP embedding of the last latent layer containing predictive autophagic features was used to evaluate the continuity of single-cell distributions between non-autophagic and autophagic states. For fast embedding, we used a parametric UMAP (pUMAP) approach where neural nets were trained to map the high-dimensional latent space features to a two-dimensional UMAP embedding. b, The average autophagy % in test data for neural nets, trained with various hyperparameters controlling the size and shape of the architecture, ranked by validation loss, consistently shows high separation between reference autophagy states for models with *elu*, *relu*, and *tanh* activation functions. c, Similar training and validation accuracies indicate low overfitting across most models. d, Sufficient model complexity is required to predict autophagy confidently. Only *relu-*based models show a positive association between increased model complexity and overfitting. Dashed lines indicate a complexity cutoff based on the model with the lowest complexity in top 1% (validation loss). Top 5 high and low complexity models (based on validation loss) were re-trained and examined further. e, Top model architectures trained for a greater number of epochs show similar performances (accuracy and loss) and robustness (correlation of predicted autophagy dynamics of WT and *vam6*Δ between experiments). Dashed lines indicate separation between low complexity (*left*) and high complexity (*right*) models. For *tanh* and *elu* architectures, low complexity models perform slightly better on average. f, Different model architectures result in very similar predicted autophagy dynamics. g, Top model architectures show high robustness in dynamics of embedded latent autophagic features using UMAP (correlation of population mean for WT and *vam6*Δ between experiments), with higher experimental correlation for models based on *elu* and *tanh*. h, UMAP dynamics of latent autophagic features are similar and consistent with reference phenotypes across different model architectures, with some activation function-dependent distributional differences in mean and variance. Points and bars indicate the mean and standard deviation at given time points across all experiments performed. The two models with the least variation in non-autophagic features are indicated by the boxes. i, Latent space UMAP embedding of control cells (model 30 and model 22). Points and bars indicate the population mean and standard deviation of an experiment at a given time point. j, Independent models yield highly consistent autophagy predictions across the genome-wide yeast library *(left panel)*, with a similar correlation to the reference controls color-coded as in *h (right panel)*. k, Association between input features and predicted autophagy log-odds.

**Figure S2.**
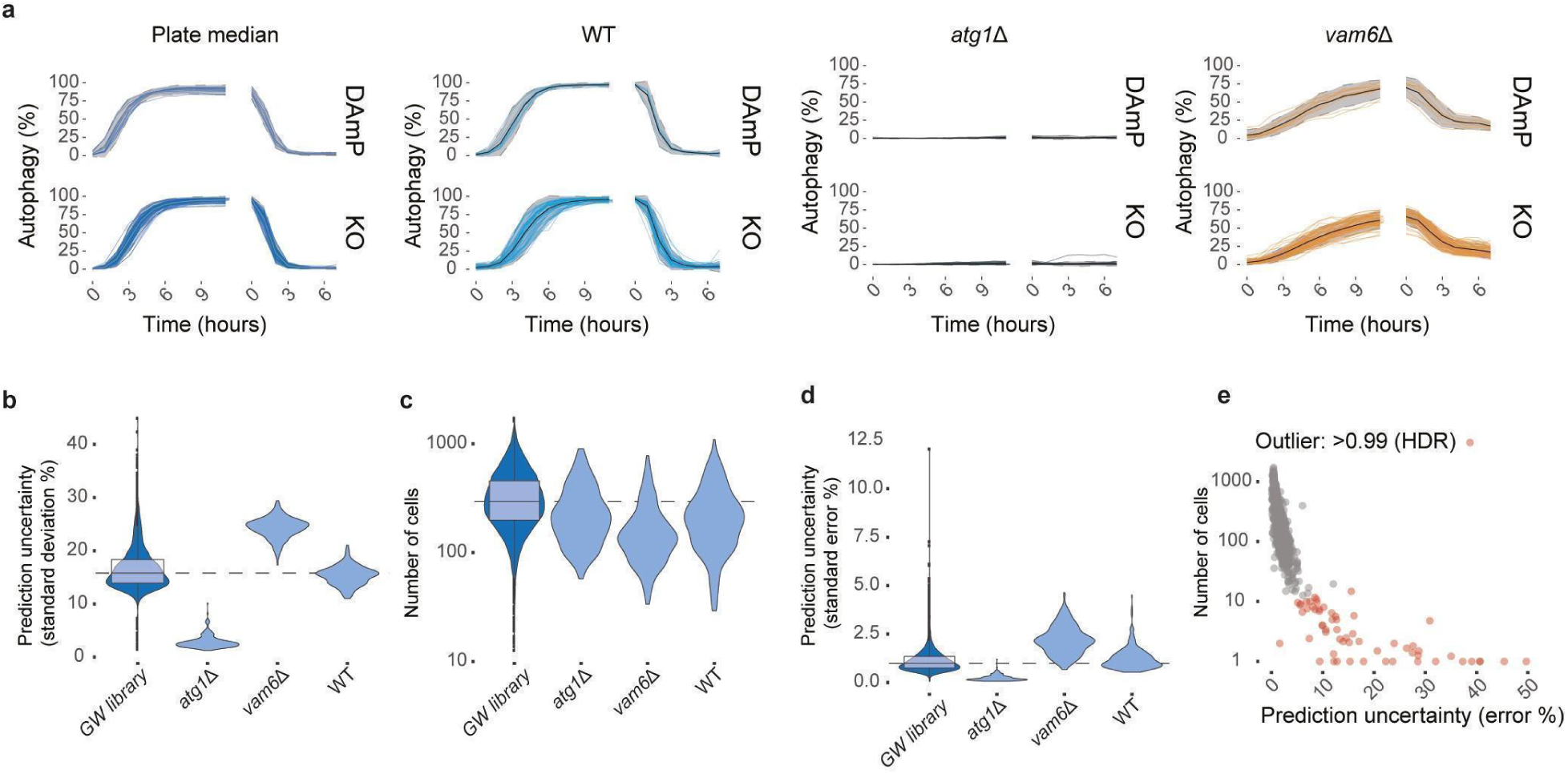
Quality control of genome-wide data. **a**, Predicted autophagy response dynamics for plate median and controls (model 30). The ribbons indicate the upper and lower boundary for plate and control filtering given by the 1.5xIQR +/- the 25%/75% quartiles, respectively. Plate medians with an average deviation from the screen median greater than 2xIQR were repeated. **b**, Prediction uncertainty, given by the average standard deviation per well for the genome-wide library and reference controls. **c**, Average number of analyzed cells per well for the genome-wide library and reference controls. **d**, Prediction uncertainty given by the average standard error per well for the genome-wide library and reference controls. **e**, Filtering of high uncertainty predictions due to low cell count, indicated by red dots. These mutants were repeated with higher seeding density.

**Figure S3.**
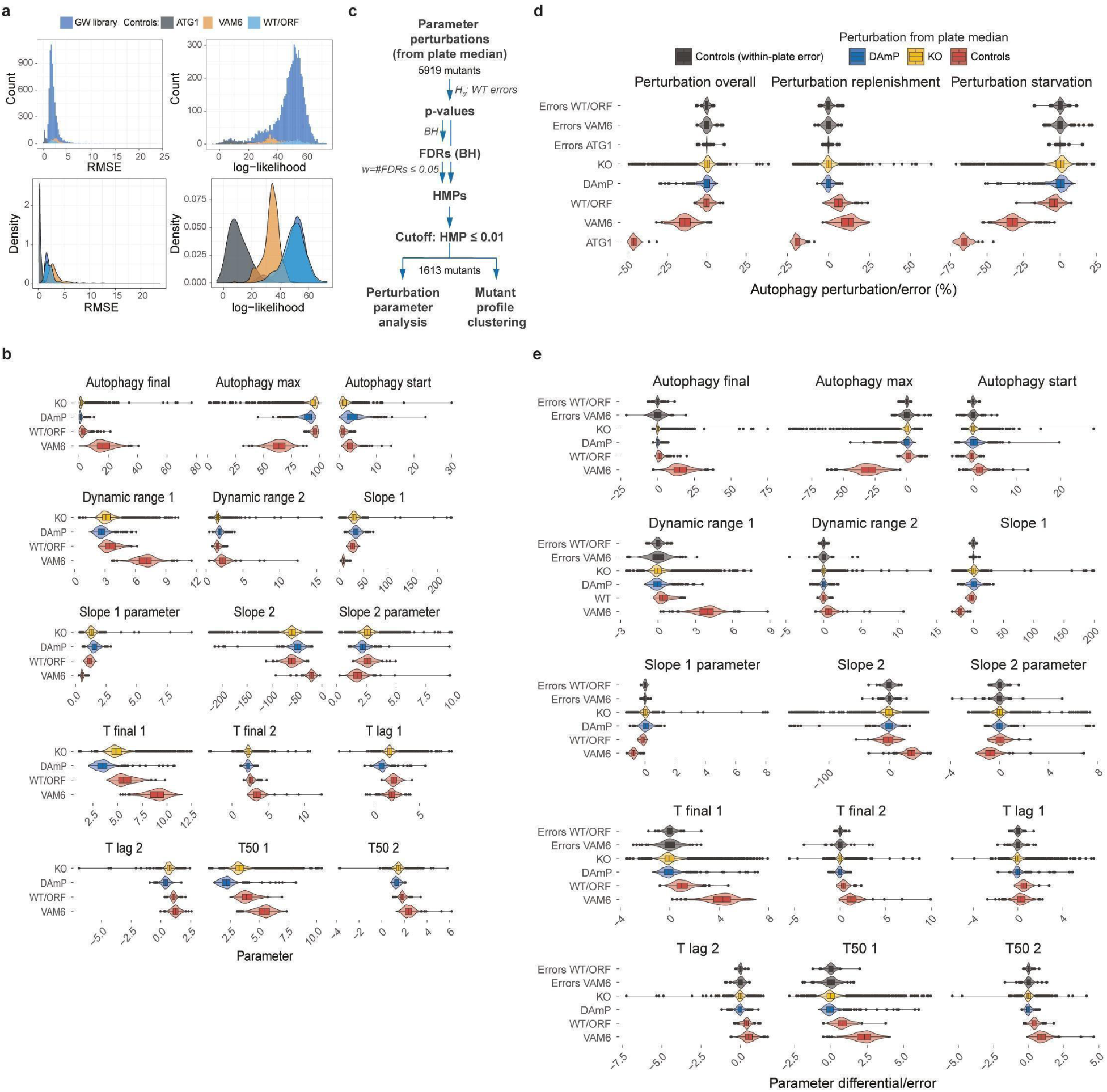
Double sigmoidal curve-fit results. **a**, Goodness-of-fit distributions for the genome-wide library and the reference controls. The RMSE distributions *(left panels)* show that all mutants have similar and low absolute curve-fit errors, while the log-likelihood score on scaled responses *(right panels)* indicates that *atg1Δ* and *vam6Δ* have lower compatibility with a sigmoidal model, consistent with the observed response curves. **b**, Sigmoidal model parameter distributions highlight differences between the KO collection strains, DAmP collection strains, and the WT control strains. **c**, Workflow for statistical analysis of autophagy perturbations. **d**, Distribution of autophagy perturbations from the plate medians for the KO collection strains, DAmP collection strains, and the control strains compared with the error distributions for the control strains. **e**, Distributions of sigmoidal model parameter differentials with the plate median for the KO collection strains, DAmP collection strains, and the control strains compared with the error distributions for the control strains.

**Figure S4.**
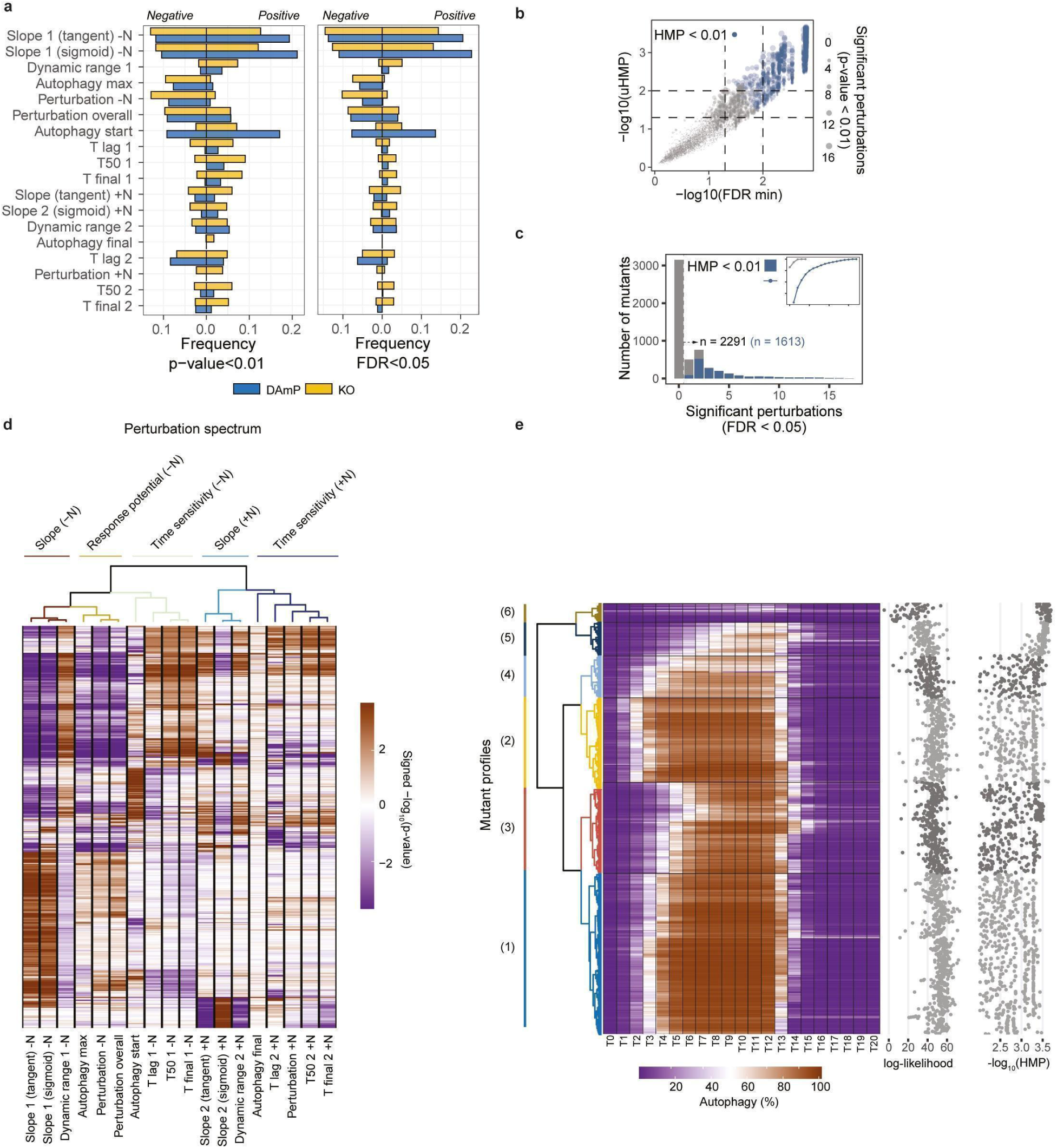
Statistics of parameter perturbations, and autophagy profile clustering. **a**, Frequency of mutants from the KO and DAmP collections having perturbations given a 1% p-value cutoff (*left*) or 5% FDR cutoff (*right*). **b**, Correlation between unweighted HMP (uHMP) and the minimum FDR over the set of hypothesis tests performed. **c**, Distribution of mutants with significant parameter perturbations (FDR ≤ 0.05), with inset indicating the cumulative probability. **d**, Hierarchical clustering of perturbation parameters using Pearson’s correlations of signed significance scores for the significant mutants (HMP ≤ 0.01), with ward.D2 clustering and dynamic tree cutting. **e**, Hierarchical clustering of autophagy response dynamics for the significant mutants (HMP ≤ 0.01), with Euclidean distances, ward.D2 clustering and dynamic tree cutting.

**Figure. S5.**
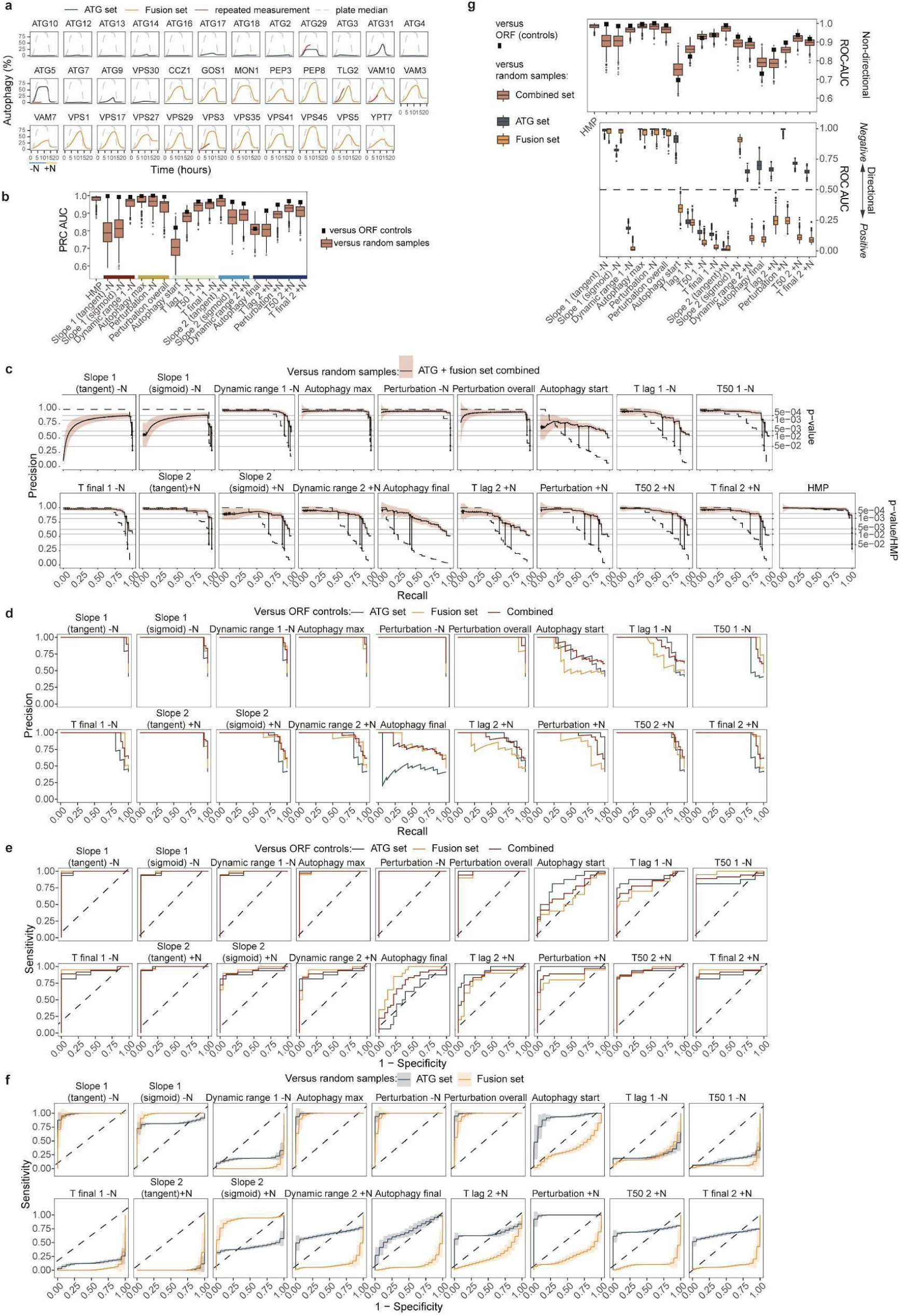
Tests for detecting autophagy reference genes. **a**, Model-fitted autophagy response curves for reference mutants. Repeated starvation measurements from a repetition screen are indicated in red for some genes. **b**, Area under the curve for PRC curves over ranked p-values for kinetic perturbation parameters comparing autophagy reference test sets versus equisized randomly sampled negative sets or ORF negative controls (squares). **c**, PRC curves for perturbation parameters and HMP ranked by p-values comparing a combined autophagy reference test set (n=35) versus equisized randomly sampled negative sets. The dashed lines represent the p-values and HMP for the recalled genes. **d**, PRC curves for perturbation parameters ranked by p-values comparing a combined autophagy reference test set (ATG: n=16; Fusion: n=19) versus an ORF negative control set (n=23). **e**, ROC curves for perturbation parameters ranked by p-values comparing a combined autophagy reference test set (ATG: n=16; Fusion: n=19) versus an ORF negative control set (n=23). **f**, ROC curve for perturbation parameters ranked by their signed p-values comparing ATG genes reference test set (n=16) and fusion genes reference test set (n=19) versus equisized randomly sampled negative sets. **g**, Area under the curve for ROC curves over ranked p-values (*top panel*) and over ranked signed p-values (*bottom panel*) for kinetic perturbation parameters comparing autophagy reference test sets versus equisized randomly sampled negative sets or ORF negative controls (squares).

**Figure S6.**
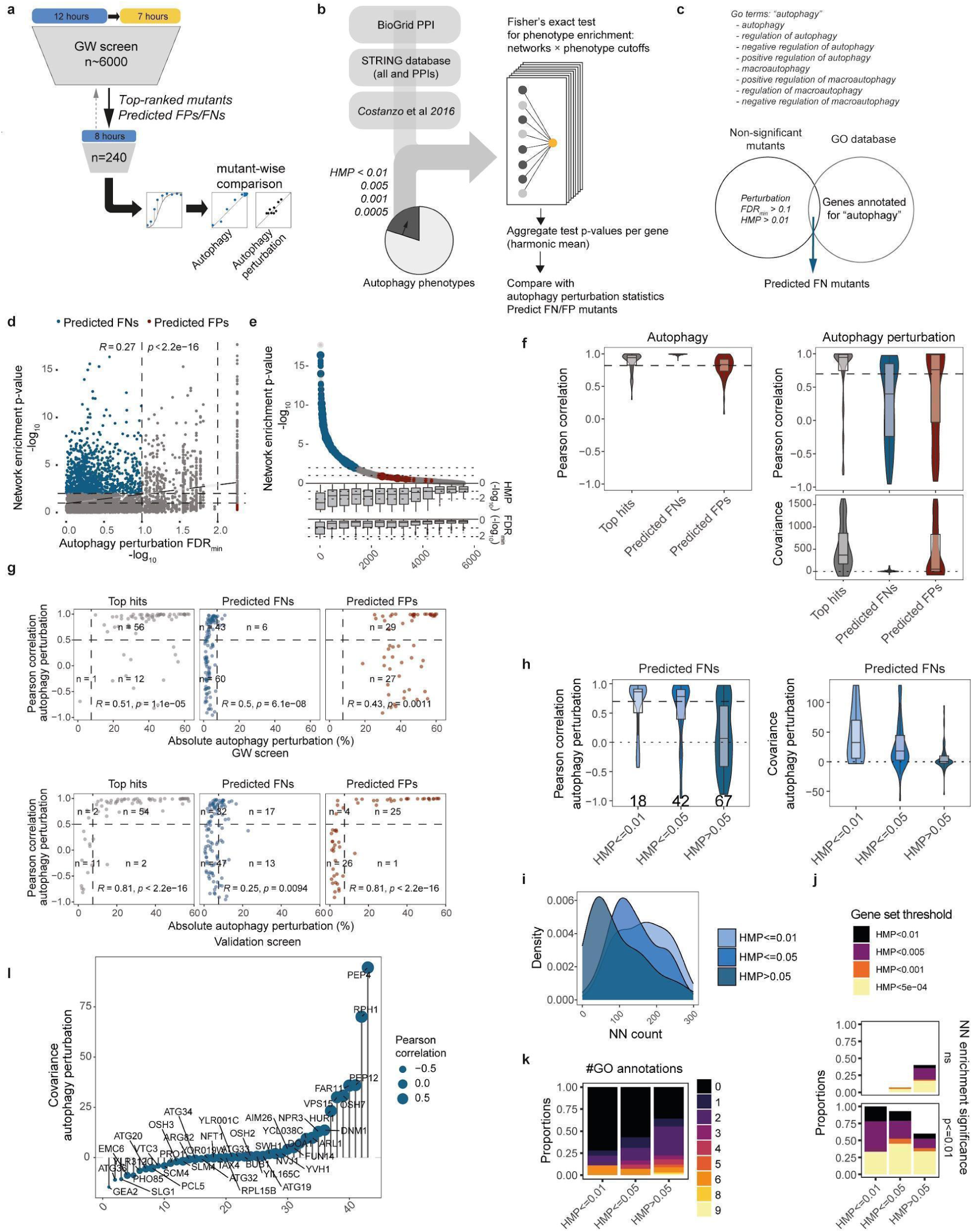
Reproducibility of top hits, and potential errors. **a**, The first stage of reproducibility testing was done by selecting the top-ranked mutants based on the HMP along with predicted false positives and negatives and repeating the starvation protocol (see methods 1.3). Subsequently, the mutant-wise correlation in autophagy dynamics and perturbation dynamics was compared between the two screens. **b**, Prediction of false positives and negatives based on enrichment tests of significant phenotypes (generated by different HMP cutoffs) in the set of first-degree neighbors for different network sources. The enrichment test p-values were aggregated by taking their harmonic mean to an overall network enrichment p-value and compared with the autophagy perturbation statistics of the respective genes. **c**, Non-significant genes annotated with the indicated autophagy GO terms were predicted as false negatives. **d**, Network enrichment p-value correlates with autophagy perturbation FDRmin (minimum perturbation BH-FDR value). Potential false positives or negatives were predicted in the off-diagonal areas as indicated. **e**, Network enrichment p-value is concordant with both autophagy perturbation FDRmin and HMP. **f**, Distribution of mutant replicate correlation of autophagy (top left) and autophagy perturbation (top right) for top hits and predicted false positives and negatives. Bottom right panel shows the replicate perturbation covariance. **g**, Mutant replicate correlation of autophagy perturbations for top hits and predicted false positives and negatives versus average absolute autophagy perturbation for GW data *(top panel)* and validation data *(bottom panel)*. The vertical lines represent the 2x median perturbation of the screen. *R* indicates the Spearman correlations, and *p* indicates their respective p-values. **h**, Stratifying the predicted false negatives based on less stringent HMP cutoffs, shows that weakly significant screen mutants display weak but reproducible responses, in contrast to non-significant mutants. *i-j*, Weakly significant and reproducible have higher counts of nearest neighbours with autophagy phenotypes (*i*) that are more likely to be significantly enriched (adjusted p-value ≤0.01) within their most enriched network (*j*). **k**, Predicted false negatives that are non-significant under the HMP were disproportionately predicted based on GO annotation. l, Perturbation covariance of non-significant mutants with autophagy related GO annotation.

**Figure S7.**
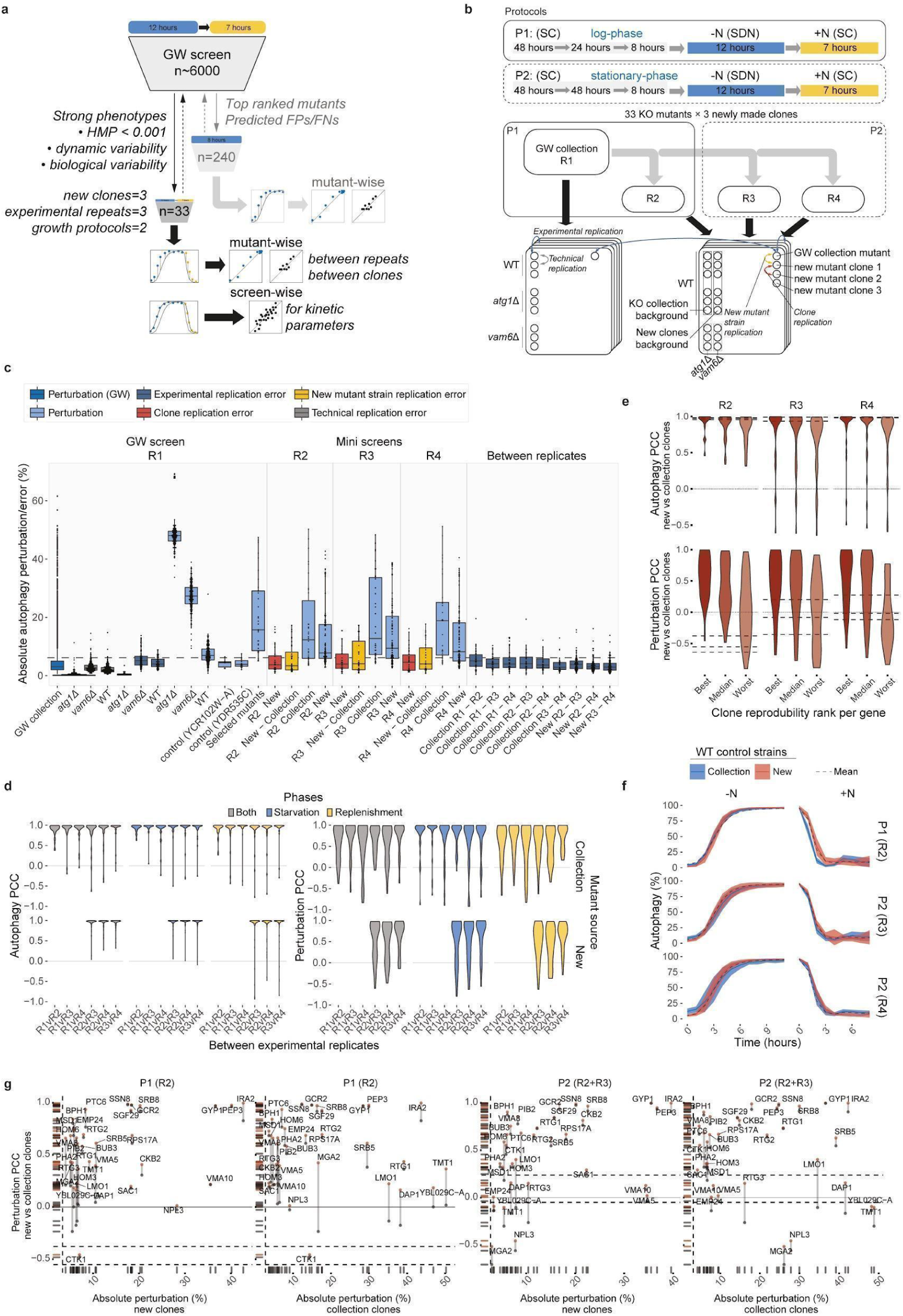
Reproducibility across experimental replicates, strain background, and experimental protocols. **a**, The second stage of reproducibility testing was performed by selecting a sub-library of strongly significant (HMP ≤ 0.001) verified gene deletions with diverse perturbation phenotypes (dynamic variability and covering multiple gene classes for biological variability). For each mutant three new clones were made in a new background strain, and the entire experimental protocol was repeated three times under two distinct growth preparation protocols (see methods 7.1 and 7.3). Reproducibility was assessed in a mutant-wise manner by comparing autophagy dynamics and perturbation dynamics, and in a screen-wise manner by comparing computed kinetic parameters. **b**, Workflow of experimental organization for comparing reproducibility. P1 refers to a growth preparation protocol where the cells were grown to log phase before being transferred to fresh media 8 hours before the experimental procedure. P2 refers to a growth preparation protocol where the cells were grown to stationary phase before being transferred to fresh media 8 hours before the experimental procedure. The genome-wide screen R1 and replicate screen R2 were subjected to P1 and replicate screens R3 and R4 were subjected to P2. Colored arrows indicate the different levels of replication tested. **c**, Comparison of autophagy perturbation and mutant-wise replication errors with the same color-coding as in *b*. The horizontal dashed line represents the 2x median perturbation of the screen. **d**, Mutant-wise replicate correlation of autophagy (*left panels*) and autophagy perturbation (*right panels*) for collection clones and new clones across biological replicates. **e**, Distribution of autophagy (*top panels*) and autophagy perturbation (*bottom panels*) correlation between collection clones and new clones for each experimental replicate. The distributions are stratified based on the ranked clone correlation from best to worst for each gene. The dashed horizontal lines represent the best, median, and worst correlation between WT control strains of the collection and new mutants. The median of all WT replicates was used as a plate control for computing autophagy perturbations, and a negative correlation for R2 indicates a distinct WT strain effect. **f**, Autophagy response dynamics for collection (Y7092) and new (BY4741) WT control strains. Dashed lines indicate the common median across all strains and experimental replicates for reference. **g**, Autophagy perturbation correlation between collection clones and new clones versus average absolute autophagy perturbation for new clones and collection clones across protocols. The red and black dots represent the best and median clone correlation for each gene, respectively. The dashed horizontal lines represent the best and median correlation between WT control strains of the collection and new mutants.

**Figure S8.**
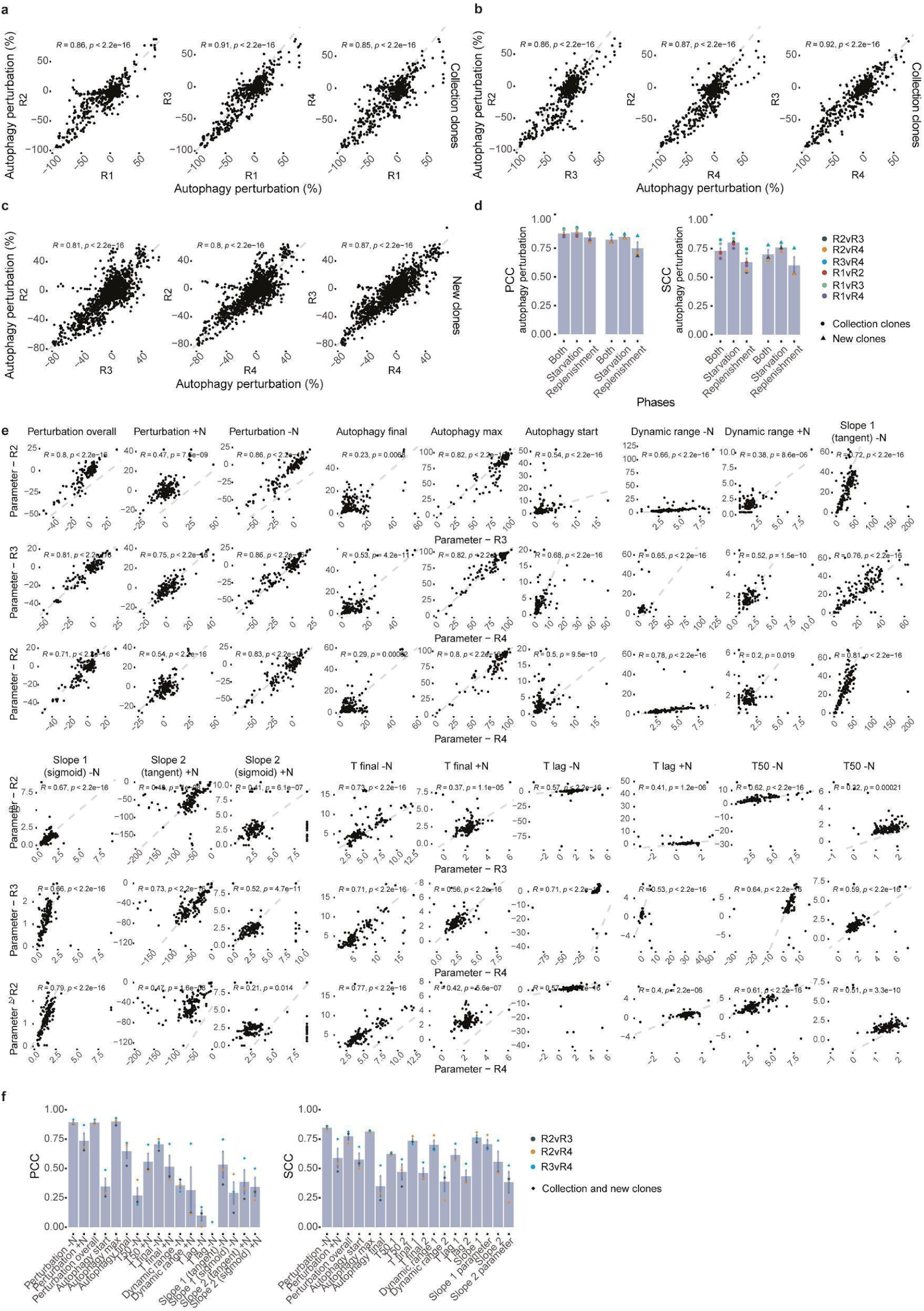
Between-screen reproducibility. **a-b**, Correlation scatter plots of point-wise autophagy perturbations for collection clones between experimental replicates. **c**, Correlation scatter plots of point-wise autophagy perturbations for new clones between experimental replicates. In *a-c, R* indicates the Pearson correlations, and *p* indicates their respective p-values. **d**, Pearson correlations (*left*) and Spearman correlations (*right*) summarizing the between-screen similarities in point-wise autophagy perturbations depending on phase or mutant source. **e**, Correlation scatter plots of response kinetic parameters from both collection clones and new clones across the experimental replicates. *R* indicates the Spearman correlations, and *p* indicates their respective p-values. **f**, Pearson correlations (*left two panels*) and Spearman correlations (*right two panels*) summarizing the between-screen similarities in response kinetic parameters.

**Figure S9.**
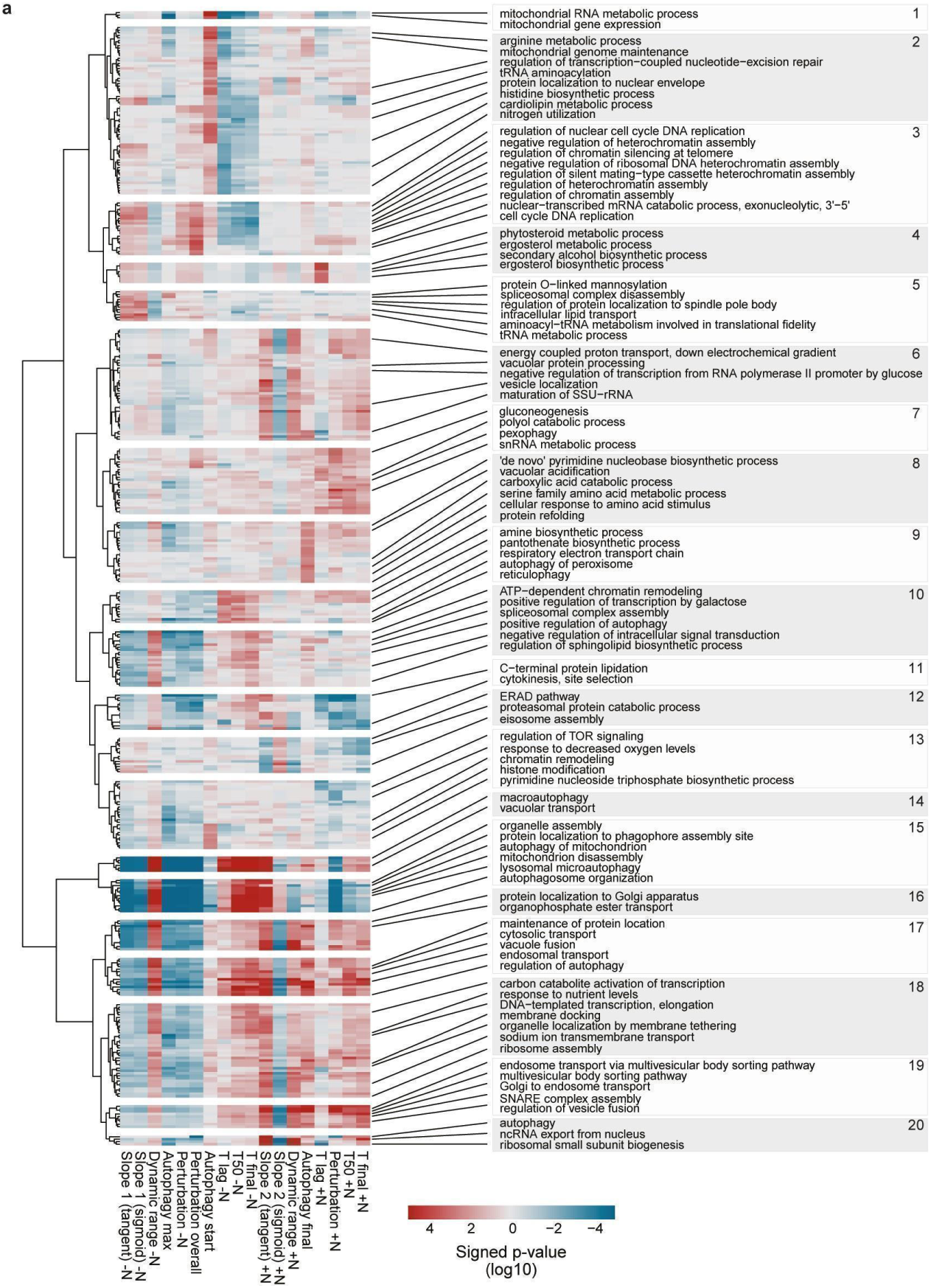
GSEA of kinetic perturbations. **a**, Hierarchical clustering of signed GSEA p-values for GO-BP terms and different kinetic perturbation parameters.

**Figure S10.**
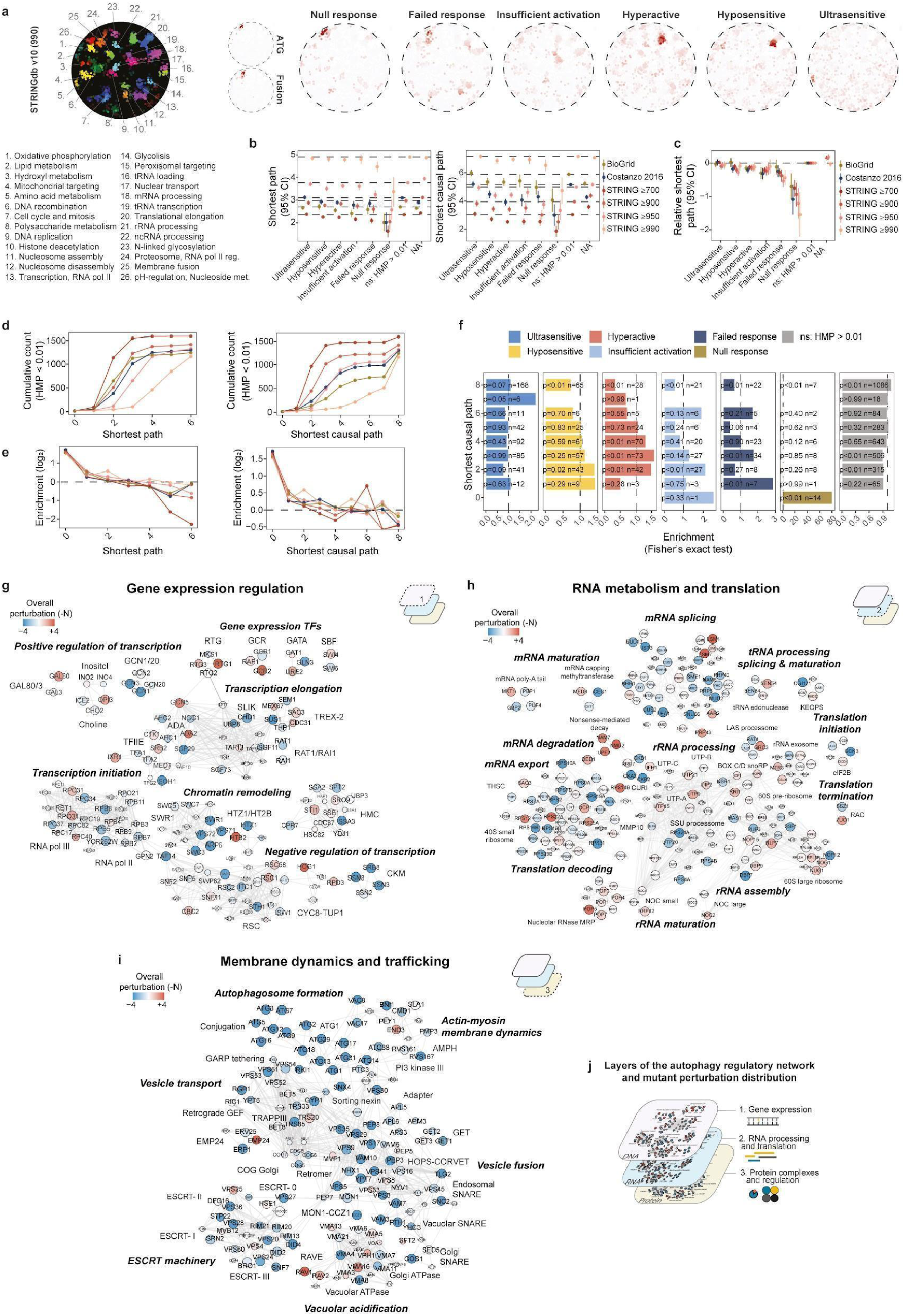
Distribution of autophagy regulating genes in genome-wide networks. **a**, SAFE analysis of the genome-wide STRING network (version 10), with a combined score cutoff of 990 for edges. Reference map of bioprocesses (left) and of ATG and fusion set (middle) for interpreting the spatial enrichment of autophagy perturbation profiles (right). **b**, Shortest pathlength and shortest causal pathlength per autophagy perturbation profile. Causal paths were determined by requiring HMP ≤ 0.01 for all intermediate nodes. **c**, Shortest path length per autophagy perturbation profiles relative to the non-significant genes (ns). **d**, Cumulative count of significant genes (HMP ≤ 0.01) per shortest path or shortest causal path distances. **e**, Cumulative enrichment (log2) of significant genes (HMP ≤ 0.01) per shortest path or shortest causal path distances. f, Enrichment of perturbation profiles per shortest causal path distances in the SGA-PCC network (Costanzo et al. 2016). **g-i**, Distribution of autophagy perturbations across gene modules spanning three regulatory network hierarchies. Node colors indicate the overall perturbation statistics (signed −log10 p-values) and the size of the nodes are proportional to their HMP (-log10). j, Hierarchy of autophagy regulatory network layers presented in *g-i*.

**Figure S11.**
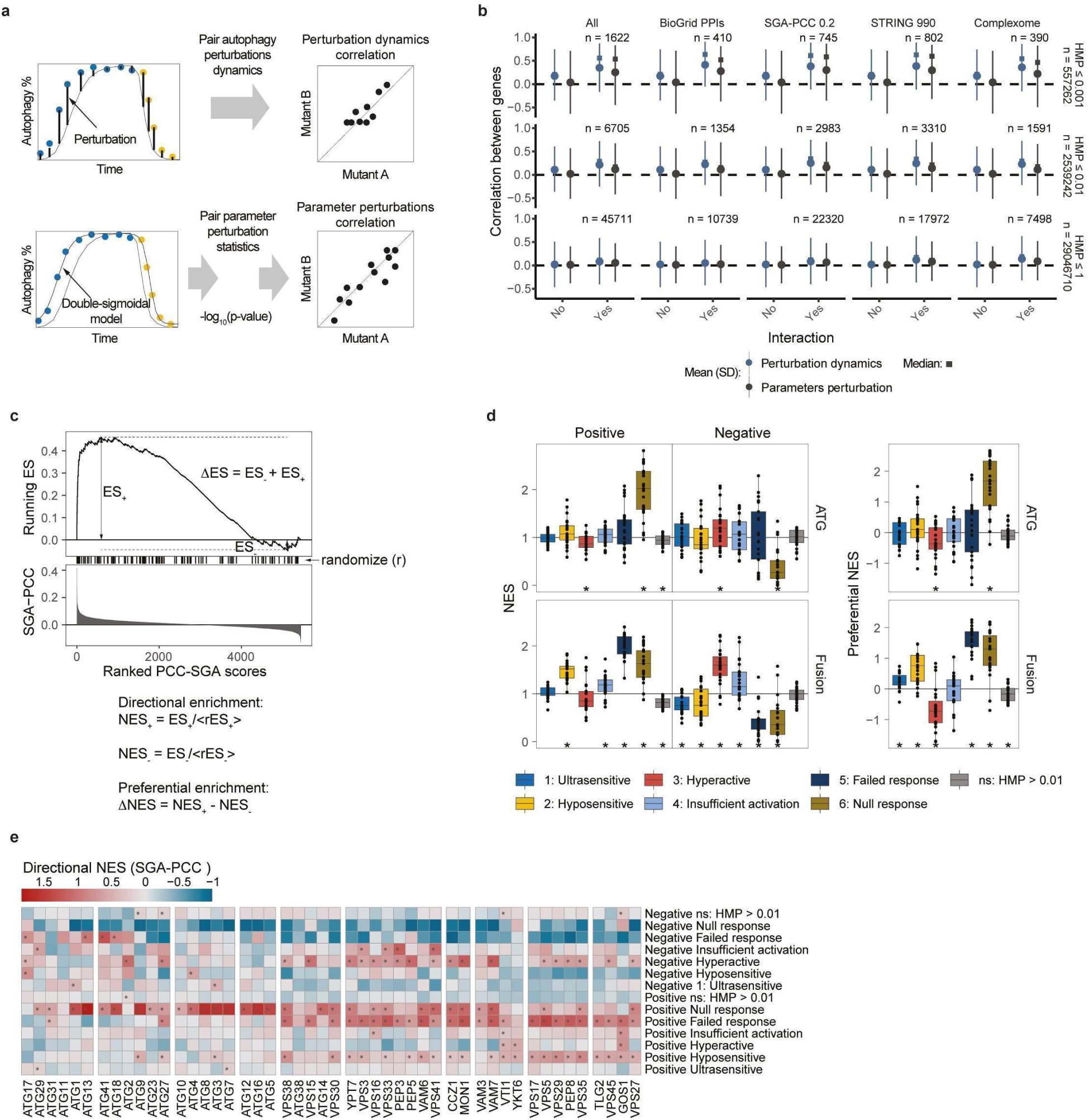
Network associations of autophagy regulatory genes. **a**, Overview of the two strategies for gene pair comparisons in *b*. **b**, Correlations in temporal autophagy (dynamics) perturbations or sets of parameter perturbations (signed −log10 p-values) between genes that share network interactions or complexome associations. Correlations were considered for all possible gene pairs (HMP ≤ 1) or only between subsets of genes with different HMP cutoffs (HMP ≤ 0.01 or 0.001). **c**, Modified gene set enrichment analysis (GSEA) to account for directional enrichment along SGA-PCC scores. **d**, Directional normalized enrichment score (*left panels*) or direction preferential normalized enrichment score (*right panels*) per perturbation profile for core ATG genes (*upper panels*) and fusion genes (*lower panels*). *p-value ≤ 0.05 (one-sample t-test). **e**, Directional enrichment of perturbation profiles along the SGA-PCC spectra for individual genes of the core autophagy machinery. *p-value ≤ 0.05 (permutation test).

**Figure S12.**
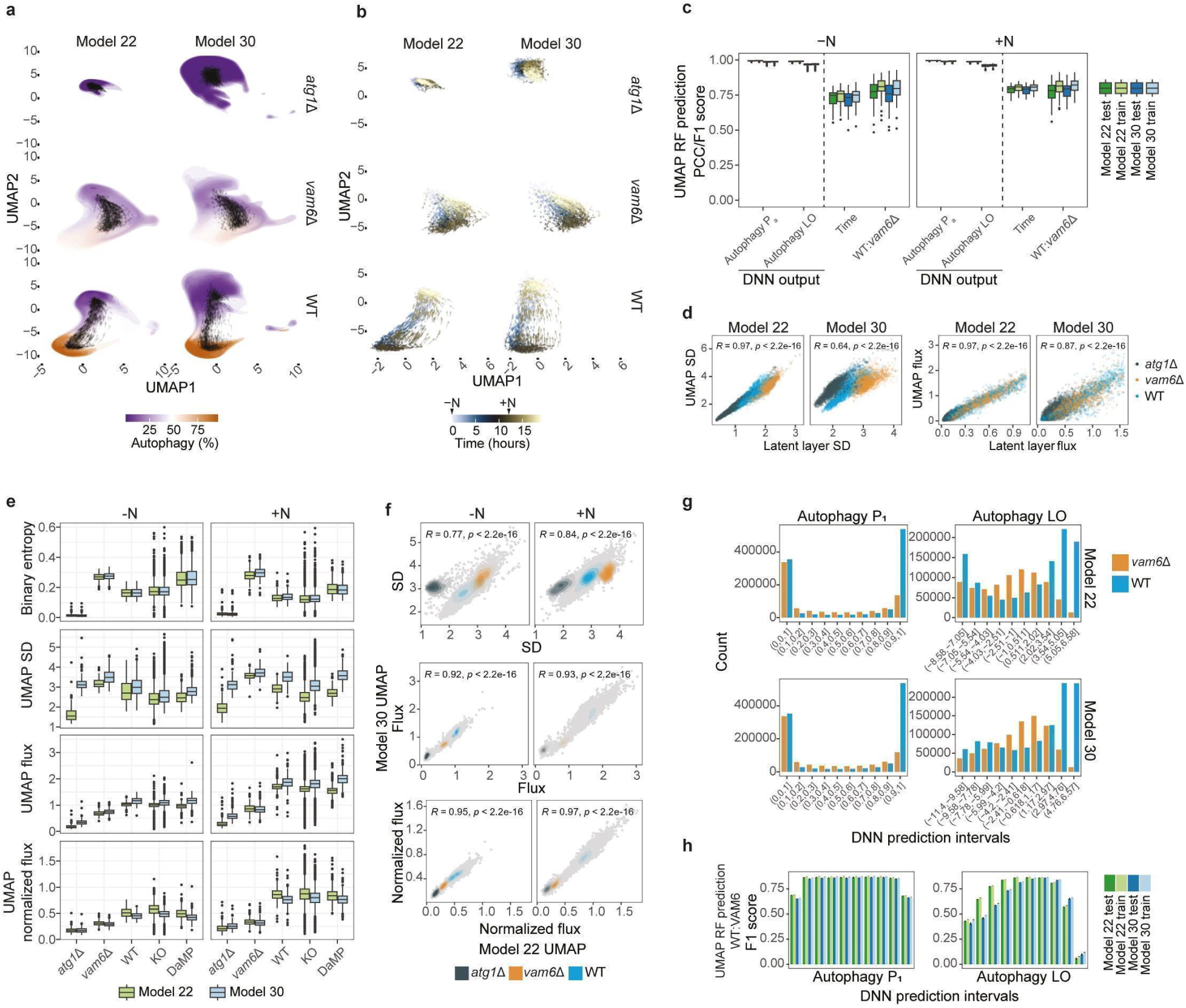
Autophagy dynamics and noise in the latent-space. **a**, UMAP-embedded latent-space distributions for control cells from model 22 and model 30. The densities represent distributions from 6 independent plates, grouped per time point and colored with the average autophagy prediction. Arrows indicate the position and direction of the average latent-space flux per plate. Left panel is presented with representative micrographs in Figure *4b*. **b**, Latent space UMAP flux for control cells from model 22 and model 30. Arrows indicate the position and direction of the average latent-space flux per plate, and color indicates the time for each population average. Note the slight greater variability of non-autophagic features (*atg1*Δ) in model 30. **c**, Training and test scores for UMAP predictions, showing that predictive information from the DNN models is fully conserved. The DNN outputs, autophagy probability (Pa) or log-odds (LO), or the sample time per cell were predicted from bagged regression trees and scored using the Pearson correlation coefficient (PCC). Classification of WT versus *vam6*Δ cells were performed with bagged classification trees and scored using an F1 score. **d**, Comparing multivariate properties of the UMAP embeddings and latent feature vectors from the DNN models of the control cells. Each point represents standard deviation (SD) and flux per cell population for a given experiment and time point. **e**, Comparing phase-averaged UMAP deviation and flux, as well as prediction uncertainty (binary entropy) across models and datasets. **f**, Comparing UMAP variance and flux between the two DNN models. Note that UMAP flux is more reproducible between the two models. *R* indicates the Pearson correlations, and *p* indicates their respective p-values. **g**, Distribution of WT and *vam6*Δ for different DNN prediction intervals. **h**, Training and test F1 scores for UMAP classification of WT versus *vam6*Δ cells per DNN prediction intervals. The classifications were performed with bagged classification trees.

**Figure S13.**
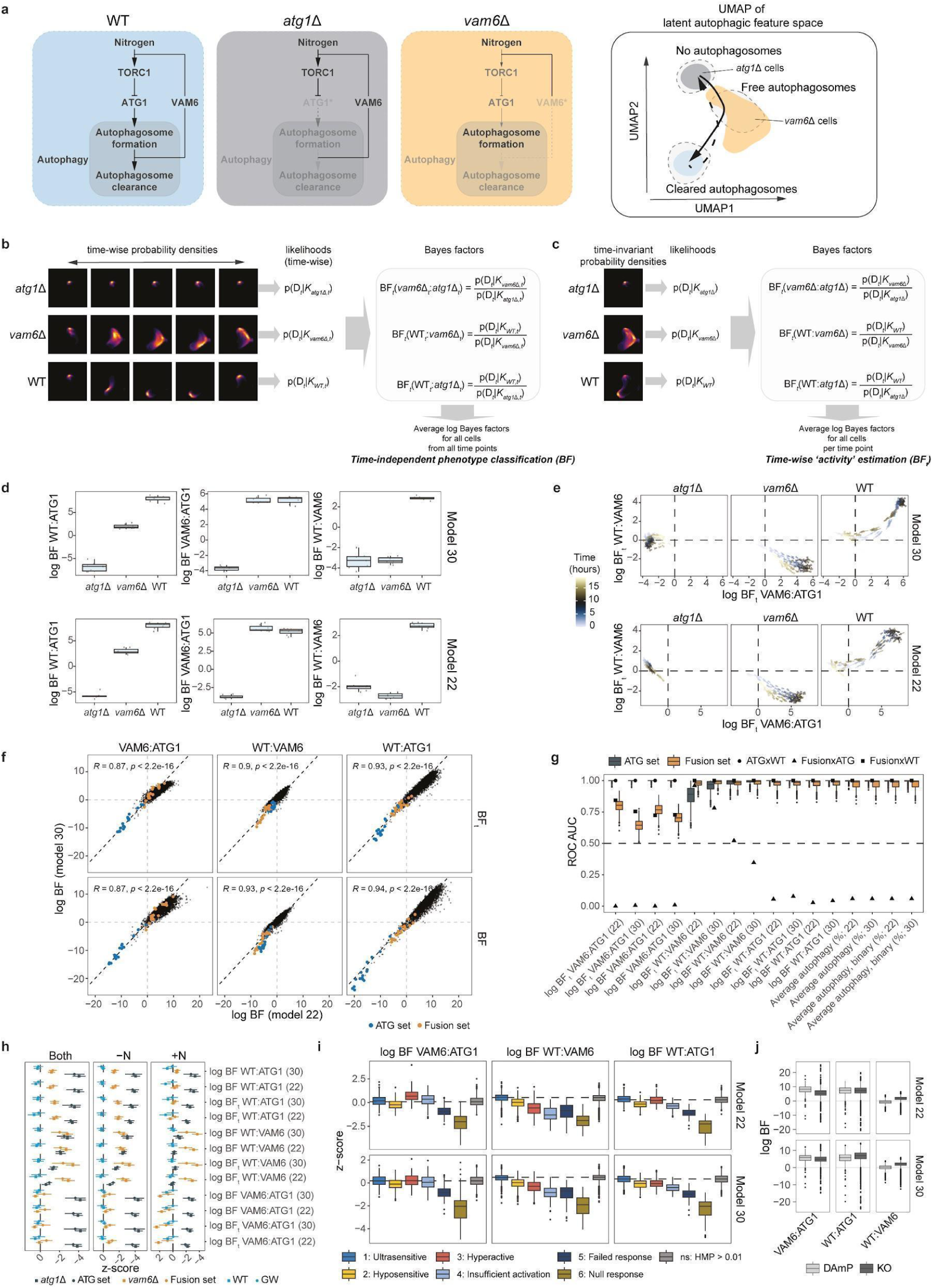
Bayesian modeling of autophagy execution from “out-of-distribution” latent-space features. **a**, Causal control conditions for learning intermediate states of autophagy execution. WT cells can execute autophagy based on nitrogen status. *atg1*Δ cells cannot produce autophagosomes. *vam6*Δ cannot clear autophagosomes and has de-regulated inhibition of autophagosome formation due to diminished TORC1 activation. The consequence is empirical latent-space distributions of *vam6*Δ cells that are populating an area between the non-autophagic and maturely autophagic feature space of *atg1*Δ and WT cells. The latent-space distributions for the three mutants can be used as reference states in a Bayesian modeling framework using Bayes factors (BFs). **b**, BF scores for classifying mutant phenotypes where time is a marginalized variable by using timewise reference probability densities. **c**, BFt’s for quantifying latent-space movements between fixed reference states by using time-invariant probability densities. **d**, Time-independent log BFs from *b* for the reference mutants from 6 independent experiments. **e**, Time-wise log BFt’s from *c* for the reference mutants from 6 independent experiments, quantifying the dynamics for autophagy execution. **f**, Comparison of log BF and BFt between model 22 and model 30. *R* indicates the Pearson correlations, and *p* indicates their respective p-values. **g**, ROC-AUC scoring the separation of ATG and fusion set mutants versus equisized randomly sampled negative sets, versus ORF negative controls, or the test sets versus each other. The BF and BFt scores are compared with the predicted average autophagy perturbation and average binarized autophagy perturbation from the same neural classifiers. **h**, log BF and BFt z-scores for control cells and ATG and fusion set cells. Points and bars indicate the means and standard deviations from collection of mutants. **i**, log BF z-scores per autophagy perturbation profiles. **j**, Comparison of log BF for KO and DAmP collection.

**Figure S14.**
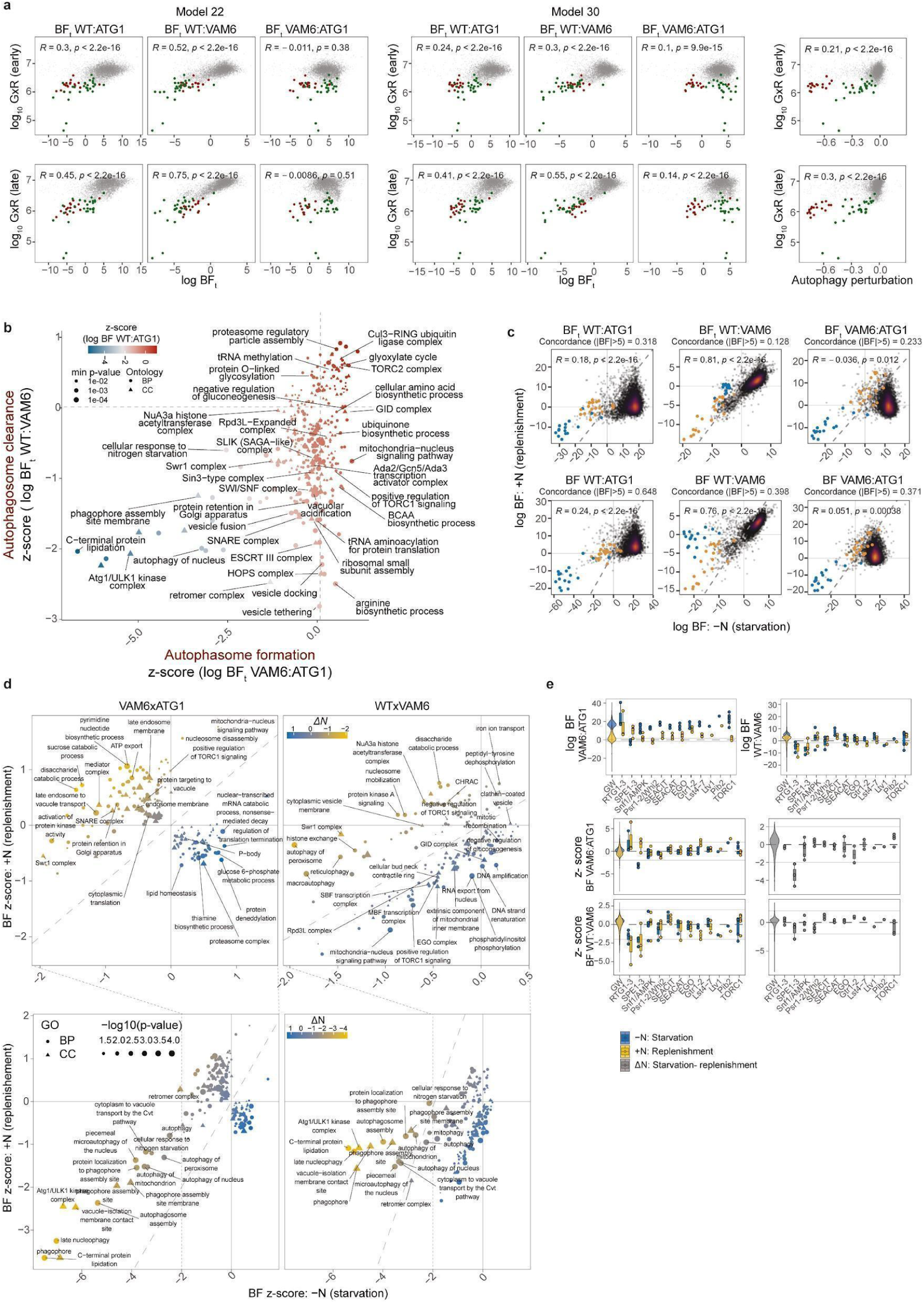
Genome-wide Bayes factor relations and GO-enrichment. **a** Genome-wide associations between BFt or average autophagy perturbation and average signal co-occurrence for (GxR) early (first two time points; *upper panels*) or (GxR) late (time points after 7 hours in -N; *lower panels*). *R* indicates the Pearson correlations, and *p* indicates their respective p-values. Green and red indicate outliers with low mNeonGreen-Atg8 and Pep4-mCherry for early time points, where red outliers were filtered out from the genome-wide analyses of BFs. **b**, Average BFt z-scores for significantly enriched GO terms. **c**, Genome-wide associations between treatment phase averages of BF and BFt for the KO collection. Blue and orange points indicate the ATG and fusion mutants, respectively. *R* indicates the Pearson correlations, and *p* indicates their respective p-values. BF classification concordances between the treatment phases (-N: starvation and +N: replenishment) are reported above each plot. Note that time-independent BFs have higher concordances compared to the phase-averaged time-wise BFt values. **d**, Comparing the average BF z-scores for GO terms per treatment phase, starvation and replenishment, that had a significant differential enrichment between the phases (p-value ≤ 0.05). **e**, BFs (*upper panels*) or BF z-scores (middle and *lower panels*) for different components of the TORC1 signaling pathway, the RTG pathway, and the spermidine synthesis pathway. The BFs have been computed per phase (-N and +N; *upper* and *middle panels*) and as a differential between phases (starvation-replenishment) (*lower panels*).

**Figure S15.**
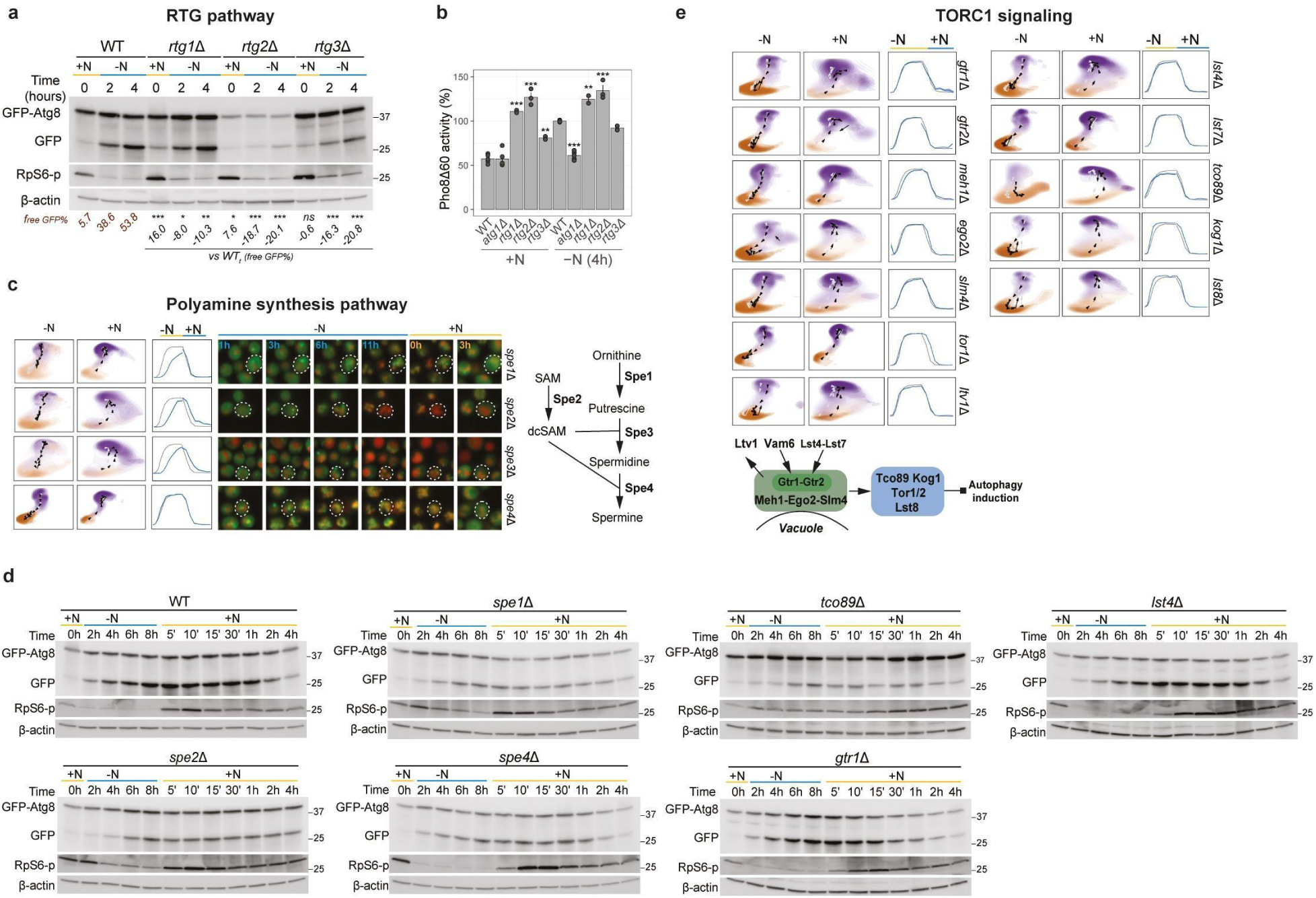
Experimental validation of autophagic flux perturbations. **a**, Immunoblot analysis of GFP-Atg8 cleavage and RpS6 phosphorylation for *rtg1*Δ, *rtg2*Δ, *rtg3*Δ, and WT cells in +N and -N for the indicated time points. +N t=0h refers to cells before shifting to -N. β-actin is used as a loading control. *p ≤0.05**p ≤0.01;***p ≤0.001; mixed model regression (n=4). **b**, Pho8Δ60 activity (%) as a measure of autophagic flux in WT (n=5), *atg1*Δ (n=5), *rtg1*Δ (n=2), *rtg2*Δ (n=3), and *rtg3*Δ (n=2) cells in +N (before starvation) and followed by -N for 4 hours. *p ≤0.05**p ≤0.01***p ≤0.001; ANOVA with TukeyHSD. **c**, UMAP-embedded latent-space distributions and autophagy response dynamics for gene deletions of the spermidine synthesis pathway with representative micrographs from indicated time points, **d**, Representative images of immunoblot analysis of GFP-Atg8 cleavage and Rps6 phosphorylation for WT (n=7), *spe1*Δ (n=3), *spe2*Δ (n=3), *spe4*Δ (n=3), *tco89*Δ (n=2), *gtr1*Δ (n=2), and *lst4*Δ (n=2) in -N followed by +N for the indicated time points. 1-actin is used as a loading control. See methods for replicas and quantifications. **e**, UMAP-embedded latent-space distributions and autophagy response dynamics for gene deletions of selected gene deletions related to TORC1 activation.

**Figure S16.**
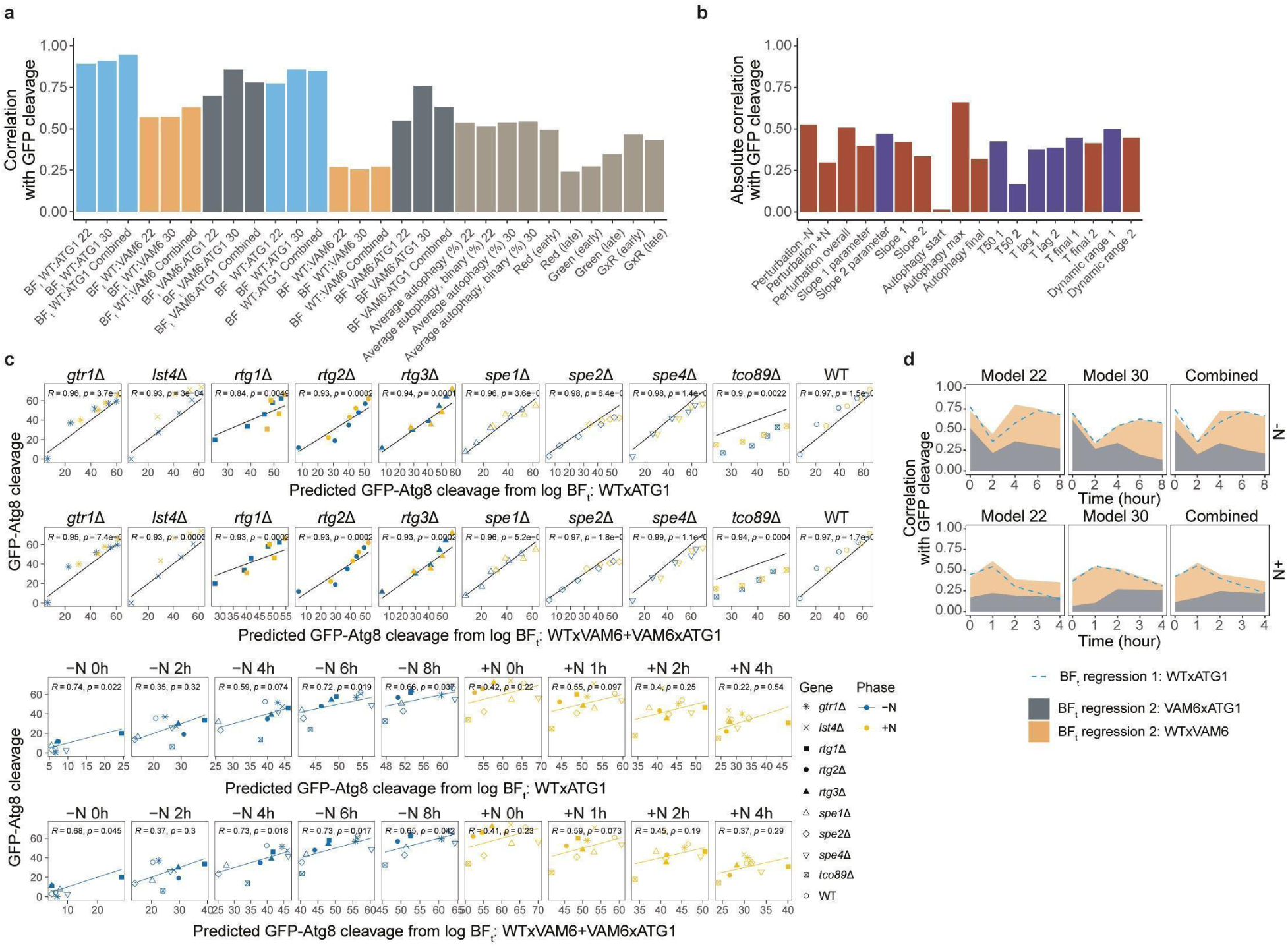
Comparing Bayes factors with autophagic flux. **a**, GFP-Atg8 cleavage (%) correlations with average Bayes factors, autophagy predictions, and fluorescence signals. **b**, GFP-Atg8 cleavage (%) correlations with autophagy perturbation parameters. Red and blue indicate positive and negative correlations, respectively. **c**, Time-wise GFP-Atg8 cleavage (%) correlations with BFt-based regression predictions, using either a univariate model (WT:ATG1) or a bivariate model (VAM6.ATG1 + WT:VAM6). *R* indicates the Pearson correlations, and *p* indicates their respective p-values. **d**, Time-wise correlations of GFP-Atg8 cleavage % with regression model predictions based on BFt from DNN model 22, model 30 or the average of the models combined. The colored area for regression model 2 represents the relative contribution of the two model covariates.

**Figure S17.**
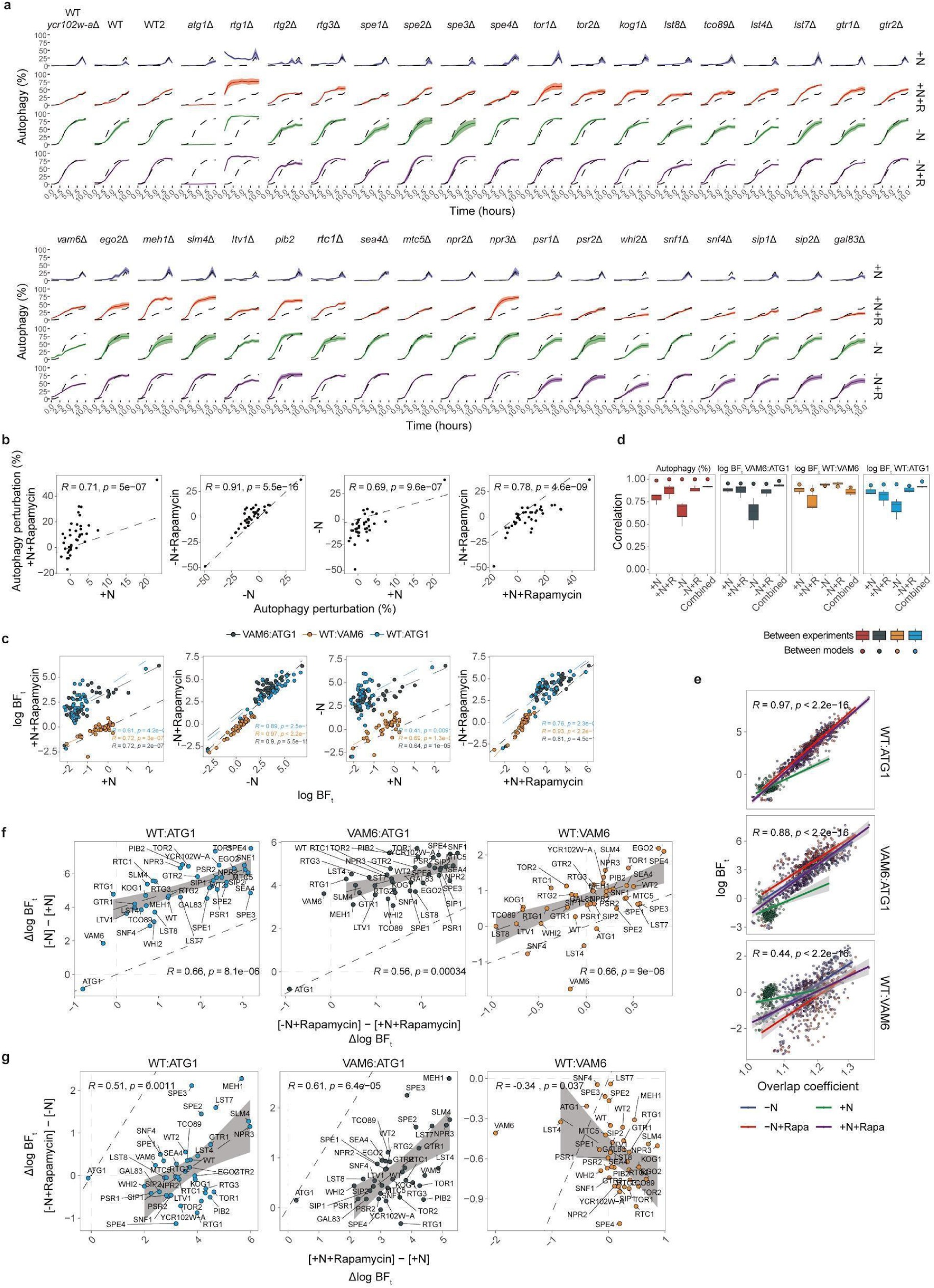
Rapamycin response screen. **a**, Autophagy (%) response dynamic in response to starvation, rapamycin or both for WT, ORF control and selected components of the TORC1 signaling pathway, the RTG pathway, and the spermidine synthesis pathway. **b**, Similarity in autophagy perturbations (%) between different treatment conditions. **c**, Similarity in BF responses between different treatment conditions. **d**, Reproducibility in autophagy % and BFs between models and between experimental conditions. **e**, Correlation between log BFs and overlap coefficient showing strong correspondence with overall autophagy (BF WT:ATG1) across all conditions. **f**, Comparison of the BF response to starvation dependent on the rapamycin treatment status. **g**, Comparison of the BF response to rapamycin treatment dependent on the nitrogen status.

**Figure S18.**
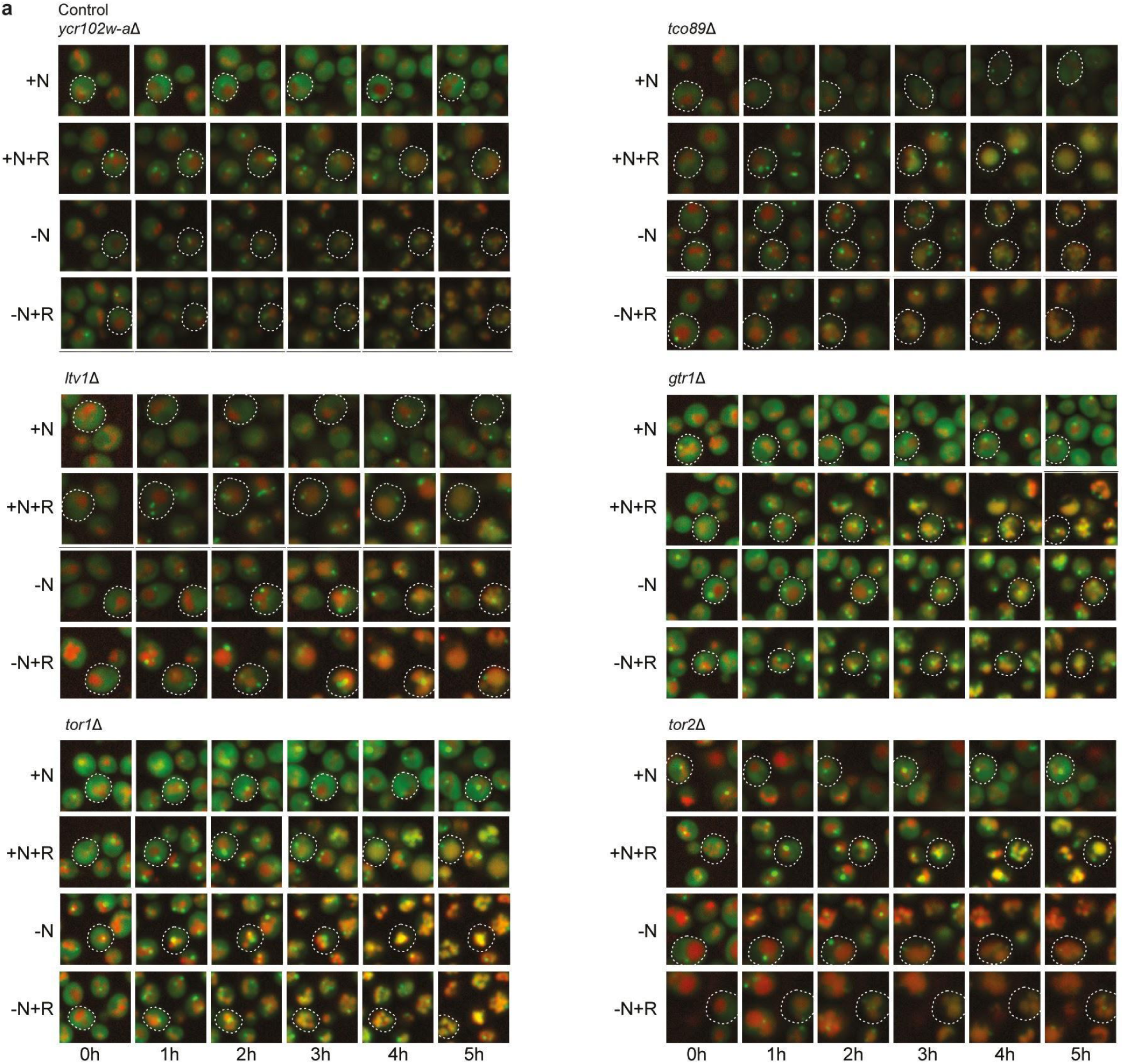
Effect of rapamycin on autophagy in TORC1 mutants. **a**, Representative micrographs depicting control and TORC1 signaling pathway mutants in response to nitrogen presence (+N) and starvation (-N), both with and without rapamycin (R), at indicated time points.

**Figure 19.**
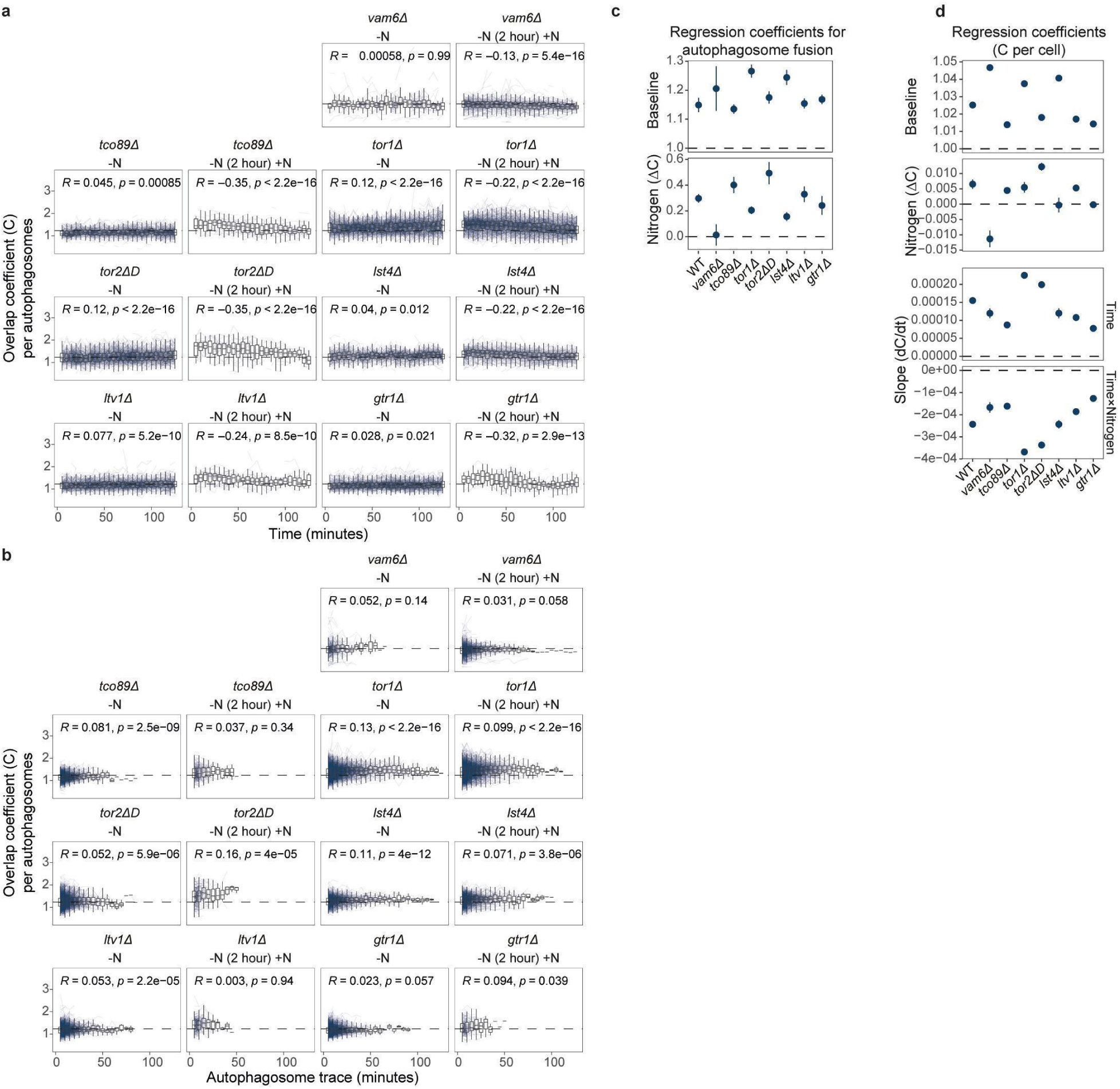
Autophagic vesicle tracking and fusion quantification. **a-b**, Mutant overlap coefficients per autophagosome over the treatment time, -N or +N (after 2 hours in -N) (*a*) or life span of vesicle traces (*b*). Blue lines indicate individual vesicle traces. Dashed horizontal lines indicate the average of all measurements. *R* indicates the Pearson correlations, and *p* indicates their respective p-values. **c**, Regression parameters for the autophagosome overlap coefficients per mutant (supplementing Figure *5g*), with the baselines (*top panel*) and changes in baselines after starving 2 hours and adding of nitrogen (*bottom panel*). **d**, Regression parameters for the cell overlap coefficients per mutant, with the baselines (*top panel*), changes in baselines after starving 2 hours and adding of nitrogen (*first middle panel*), slopes for changes over time (*second middle panel*), and changes in slopes under addition of nitrogen (*bottom panel*). In *c* and *d* points and bars indicate mean and 95% confidence interval for the regression estimates.

**Figure S20.**
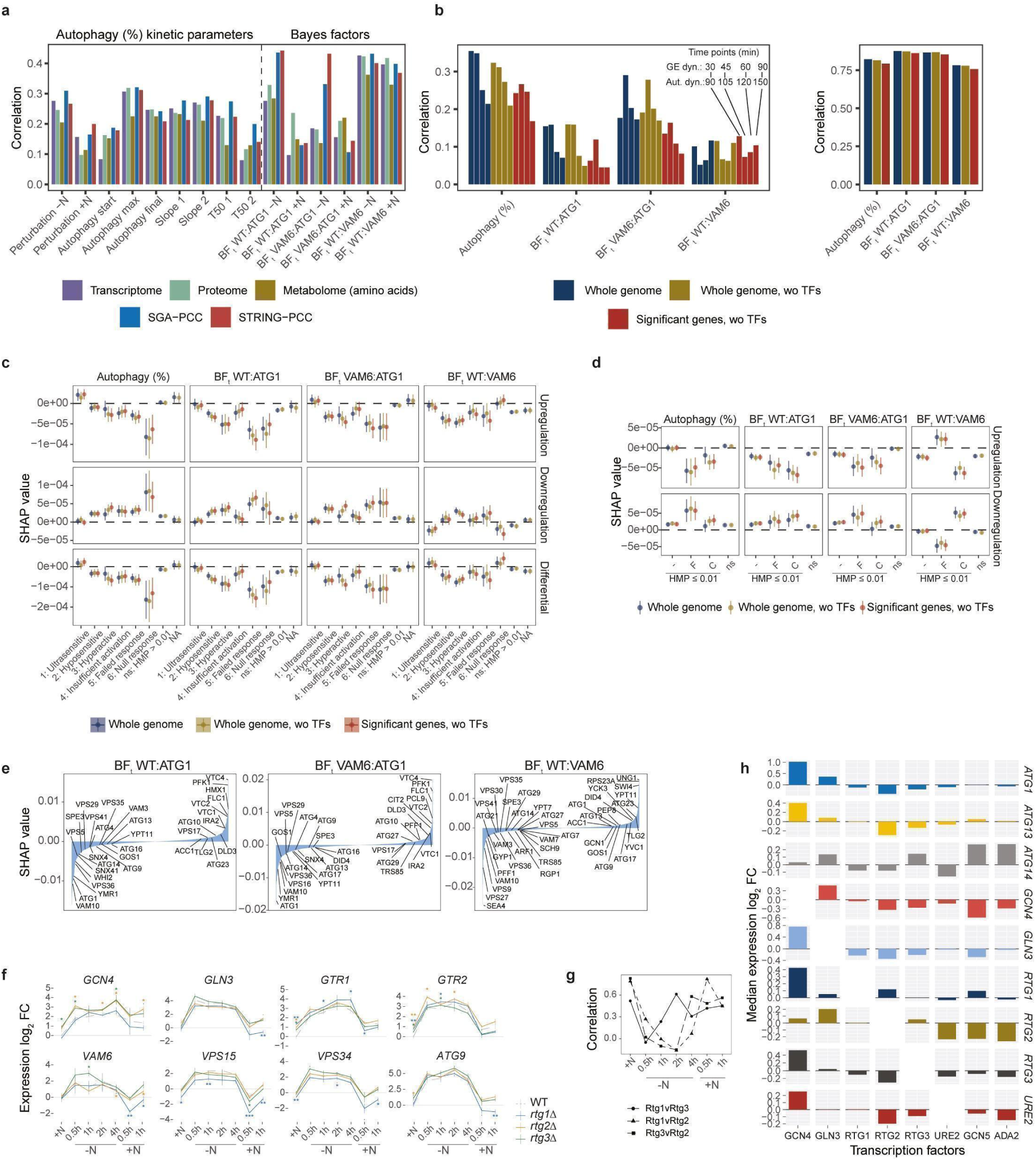
Cross-omics prediction of autophagy dynamics. **a**, Benchmarking prediction of different autophagy metrics using 10-fold training and testing. Model performance as median test correlation between measured and predicted values. Transcriptome: Kemmeren et al. 2014 ^49^; Proteome: Messner et al. 2023 ^50^; Metabolome: Mülleder et al. 2017 ^51^; SGA-PCC: Costanzo et al. 2016 ^29^. **b**, Benchmarking prediction of autophagy perturbation dynamics from TF expression induction profiles ^52^ using 10-fold training and testing, with random sampling of the entire TF time-series (*left panel*) or random sampling of individual measurements (*right panel*). Model performance as median test correlation between measured and predicted values. Color coding indicates different feature pre-selection criteria. **c**, Adjusted SHAP values for regulation of gene expression across models and autophagy metrics comparing feature classes based on autophagy perturbation profiles. Points and bars indicate mean and standard error. **d**, Adjusted SHAP values for regulation of gene expression comparing feature classes across models and autophagy metrics. F: formation genes; C: clearance genes with negative BFt z-scores beyond two standard deviations from the mean. Points and bars indicate mean and standard error. **e**, Gene-wise SHAP values for differential regulation of gene expression predicting BF responses. **f**, qRT-PCR for expression of *GCN4*, *ATG13*, *GTR1*, *GTR2*, *VAM6*, *VPS15*, *VPS34*, and *ATG9* in *rtg1*Δ, *rtg2*Δ, *rtg3*Δ, and WT cells subjected to 4 hours of -N followed by 1 hour of +N. Graphs show mean and standard error (n=3). *p ≤ 0.05;**P ≤ 0.01;***P ≤ 0.001; t-test. **g**, Correlations in differential expression between per time-point for the panel of genes targets analyzed in Figure *6e* and Suppl. Figure *S20f*. **h**, Median gene expression change in response to TF induction in Hackett et al. 2020 ^52^.

**Figure S21.**
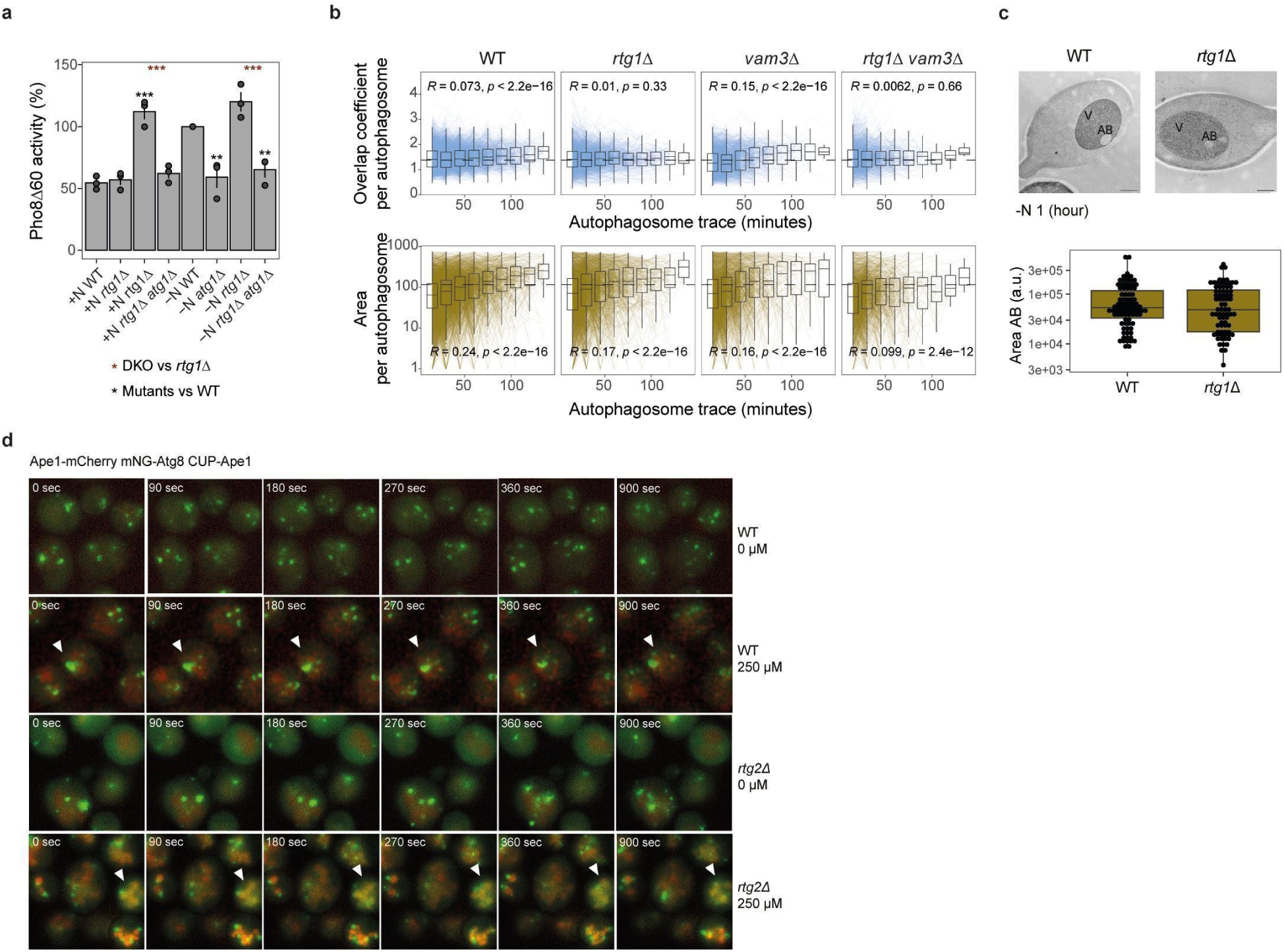
Influence RTG pathway on autophagosome formation and fusion. **a**, Pho8Δ60 activity (%) as a measure of autophagic flux in WT, *atg1*Δ, *rtg1*Δ, and *rtg1*Δ *atg1*Δ cells in +N (before starvation) and followed by -N for 4 hours. **p ≤0.01***p ≤0.001; ANOVA with TukeyHSD (n=3). **b**, Overlap coefficients (*top panels*) and area (a.u.; *bottom panels*) per autophagosome over the life span of vesicle traces, measured at 15-minute intervals, for WT, *vam3*Δ, *rtg1*Δ, and *rtg1*Δ *vam3*Δ cells after 2 hours of nitrogen starvation-induced autophagy. Blue or brown lines indicate individual vesicle traces. Dashed horizontal lines indicate the average of all measurements; cell traces were collected from n=6 independent replicates. *R* indicates the Pearson correlations, and *p* indicates their respective p-values. **c**, Representative electron micrographs showing autophagic bodies (AB) after 1 hour of nitrogen starvation in WT and *rtg1*Δ cells (*top panel*), along with quantifications of AB area (a.u.; *bottom panel*). WT: n= 93 and *rtg1*Δ: n=91. Scale bar: 500 nm; n.s., one-way ANOVA. **d**, Representative micrographs (n=4) showing Ape1 overexpression under a copper-inducible promoter, along with the Ape1-mCherry sequestration assay, with or without aggregate induction by Ape1 overexpression through CuSO4 pre-treatment for 20 hours. Forms of Ape1 encapsulation by Atg8 are indicated by arrows.

**Table S1.**
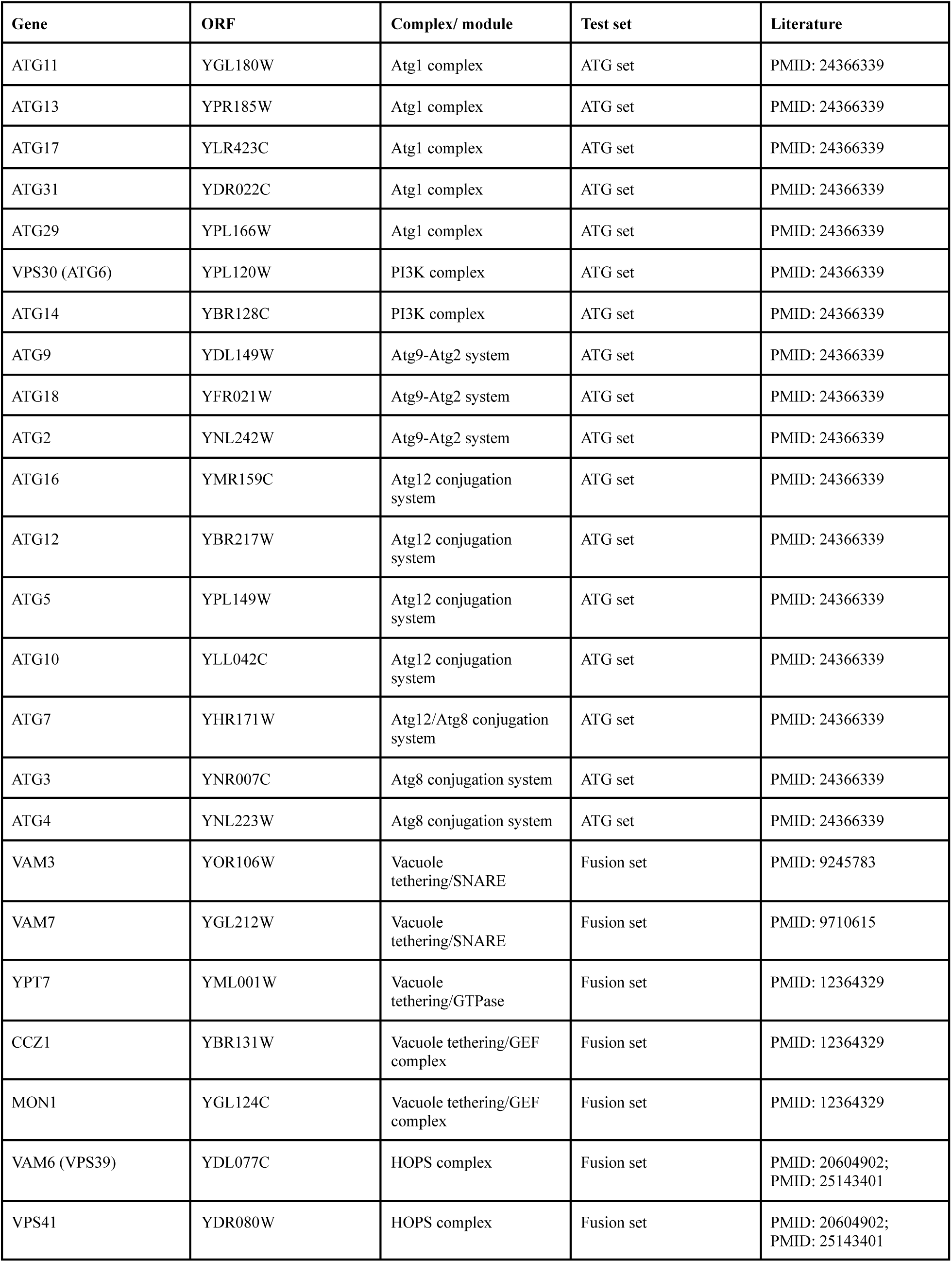
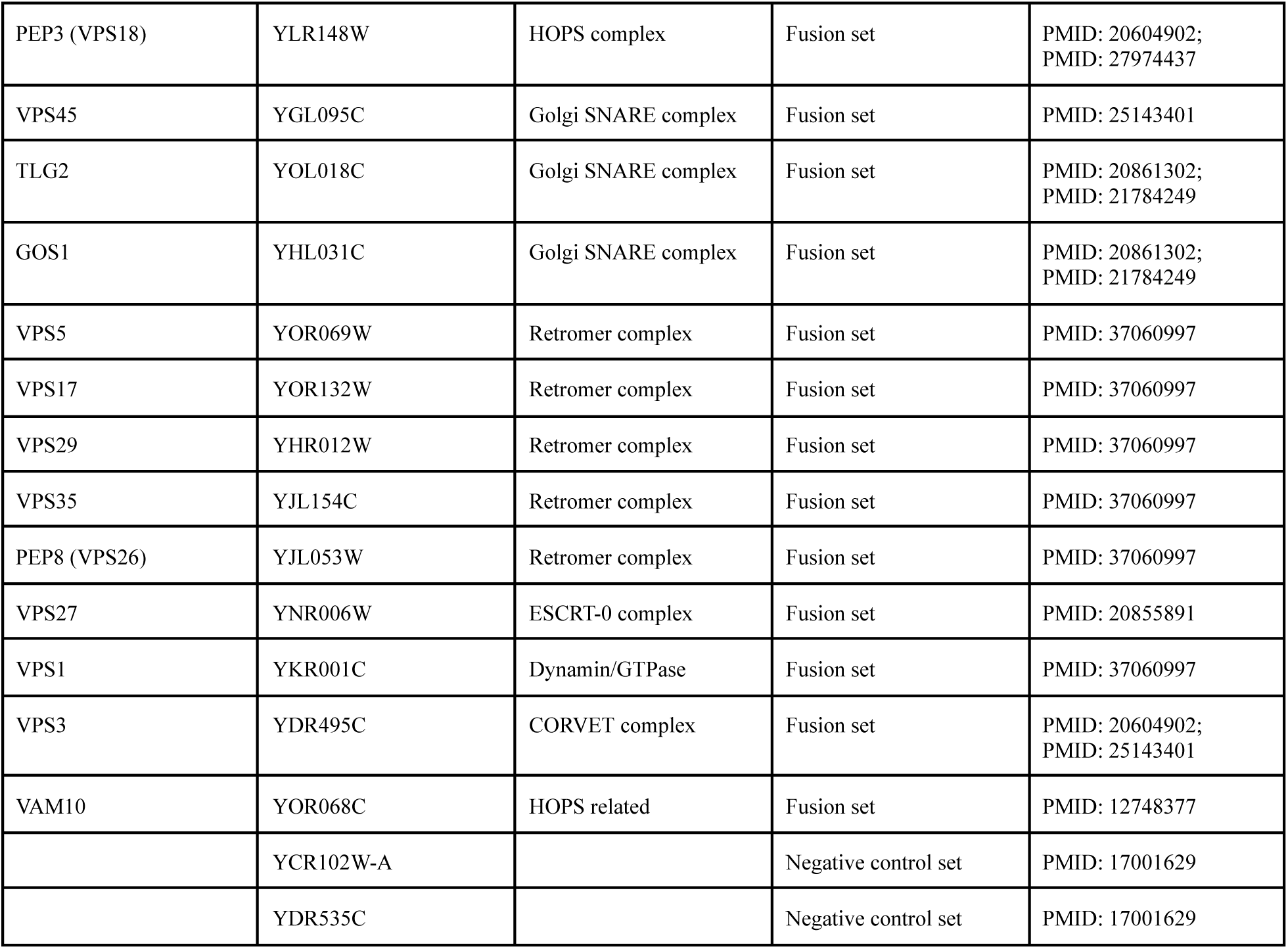
List of reference mutants for test sets.

| Gene | ORF | Complex/ module | Test set | Literature |
| --- | --- | --- | --- | --- |
| ATG11 | YGL180W | Atg1 complex | ATG set | PMID: 24366339 |
| ATG13 | YPR185W | Atg1 complex | ATG set | PMID: 24366339 |
| ATG17 | YLR423C | Atg1 complex | ATG set | PMID: 24366339 |
| ATG31 | YDR022C | Atg1 complex | ATG set | PMID: 24366339 |
| ATG29 | YPL166W | Atg1 complex | ATG set | PMID: 24366339 |
| VPS30 (ATG6) | YPL120W | PI3K complex | ATG set | PMID: 24366339 |
| ATG14 | YBR128C | PI3K complex | ATG set | PMID: 24366339 |
| ATG9 | YDL149W | Atg9-Atg2 system | ATG set | PMID: 24366339 |
| ATG18 | YFR021W | Atg9-Atg2 system | ATG set | PMID: 24366339 |
| ATG2 | YNL242W | Atg9-Atg2 system | ATG set | PMID: 24366339 |
| ATG16 | YMR159C | Atg12 conjugation system | ATG set | PMID: 24366339 |
| ATG12 | YBR217W | Atg12 conjugation system | ATG set | PMID: 24366339 |
| ATG5 | YPL149W | Atg12 conjugation system | ATG set | PMID: 24366339 |
| ATG10 | YLL042C | Atg12 conjugation system | ATG set | PMID: 24366339 |
| ATG7 | YHR171W | Atg12/Atg8 conjugation system | ATG set | PMID: 24366339 |
| ATG3 | YNR007C | Atg8 conjugation system | ATG set | PMID: 24366339 |
| ATG4 | YNL223W | Atg8 conjugation system | ATG set | PMID: 24366339 |
| VAM3 | YOR106W | Vacuole tethering/SNARE | Fusion set | PMID: 9245783 |
| VAM7 | YGL212W | Vacuole tethering/SNARE | Fusion set | PMID: 9710615 |
| YPT7 | YML001W | Vacuole tethering/GTPase | Fusion set | PMID: 12364329 |
| CCZ1 | YBR131W | Vacuole tethering/GEF complex | Fusion set | PMID: 12364329 |
| MON1 | YGL124C | Vacuole tethering/GEF complex | Fusion set | PMID: 12364329 |
| VAM6 (VPS39) | YDL077C | HOPS complex | Fusion set | PMID: 20604902;<br>PMID: 25143401 |
| VPS41 | YDR080W | HOPS complex | Fusion set | PMID: 20604902;<br>PMID: 25143401 |
| PEP3 (VPS18) | YLR148W | HOPS complex | Fusion set | PMID: 20604902;<br>PMID: 27974437 |
| VPS45 | YGL095C | Golgi SNARE complex | Fusion set | PMID: 25143401 |
| TLG2 | YOL018C | Golgi SNARE complex | Fusion set | PMID: 20861302;<br>PMID: 21784249 |
| GOS1 | YHL031C | Golgi SNARE complex | Fusion set | PMID: 20861302;<br>PMID: 21784249 |
| VPS5 | YOR069W | Retromer complex | Fusion set | PMID: 37060997 |
| VPS17 | YOR132W | Retromer complex | Fusion set | PMID: 37060997 |
| VPS29 | YHR012W | Retromer complex | Fusion set | PMID: 37060997 |
| VPS35 | YJL154C | Retromer complex | Fusion set | PMID: 37060997 |
| PEP8 (VPS26) | YJL053W | Retromer complex | Fusion set | PMID: 37060997 |
| VPS27 | YNR006W | ESCRT-0 complex | Fusion set | PMID: 20855891 |
| VPS1 | YKR001C | Dynamin/GTPase | Fusion set | PMID: 37060997 |
| VPS3 | YDR495C | CORVET complex | Fusion set | PMID: 20604902;<br>PMID: 25143401 |
| VAM10 | YOR068C | HOPS related | Fusion set | PMID: 12748377 |
|  | YCR102W-A |  | Negative control set | PMID: 17001629 |
|  | YDR535C |  | Negative control set | PMID: 17001629 |

## References

1. He, C. Balancing nutrient and energy demand and supply via autophagy. Curr. Biol. 32, R684–R696 (2022).

2. Hansen, M., Rubinsztein, D. C. & Walker, D. W. Autophagy as a promoter of longevity: insights from model organisms. Nat. Rev. Mol. Cell Biol. 19, 579–593 (2018).

3. Galluzzi, L., Pietrocola, F., Levine, B. & Kroemer, G. Metabolic control of autophagy. Cell 159, 1263–1276 (2014).

4. Aman, Y. et al. Autophagy in healthy aging and disease. Nat Aging 1, 634–650 (2021).

5. Feng, Y., He, D., Yao, Z. & Klionsky, D. J. The machinery of macroautophagy. Cell Res. 24, 24–41 (2014).

6. González, A. & Hall, M. N. Nutrient sensing and TOR signaling in yeast and mammals. EMBO J. 36, 397–408 (2017).

7. Tsukada, M. & Ohsumi, Y. Isolation and characterization of autophagy-defective mutants of Saccharomyces cerevisiae. FEBS Lett. 333, 169–174 (1993).

8. Mizushima, N. et al. A protein conjugation system essential for autophagy. Nature 395, 395–398 (1998).

9. Kirisako, T. et al. Formation process of autophagosome is traced with Apg8/Aut7p in yeast. J. Cell Biol. 147, 435–446 (1999).

10. Galluzzi, L., Bravo-San Pedro, J. M., Levine, B., Green, D. R. & Kroemer, G. Pharmacological modulation of autophagy: therapeutic potential and persisting obstacles. Nat. Rev. Drug Discov. 16, 487–511 (2017).

11. Behrends, C., Sowa, M. E., Gygi, S. P. & Harper, J. W. Network organization of the human autophagy system. Nature 466, 68–76 (2010).

12. Kramer, M. H. et al. Active Interaction Mapping Reveals the Hierarchical Organization of Autophagy. Mol. Cell 65, 761–774.e5 (2017).

13. Hu, Z. et al. Multilayered Control of Protein Turnover by TORC1 and Atg1. Cell Rep. 28, 3486–3496.e6 (2019).

14. Diehl, V. et al. Minimized combinatorial CRISPR screens identify genetic interactions in autophagy. Nucleic Acids Res. 49, 5684–5704 (2021).

15. Türei, D. et al. Autophagy Regulatory Network - a systems-level bioinformatics resource for studying the mechanism and regulation of autophagy. Autophagy 11, 155–165 (2015).

16. Mizushima, N. & Murphy, L. O. Autophagy Assays for Biological Discovery and Therapeutic Development. Trends Biochem. Sci. 45, 1080–1093 (2020).

17. Loos, B., du Toit, A. & Hofmeyr, J.-H. S. Defining and measuring autophagosome flux—concept and reality. Autophagy 10, 2087–2096 (2014).

18. Barth, H. & Thumm, M. A genomic screen identifies AUT8 as a novel gene essential for autophagy in the yeast Saccharomyces cerevisiae. Gene 274, 151–156 (2001).

19. Kira, S. et al. Reciprocal conversion of Gtr1 and Gtr2 nucleotide-binding states by Npr2-Npr3 inactivates TORC1 and induces autophagy. Autophagy 10, 1565–1578 (2014).

20. Beesabathuni, N. S., Park, S. & Shah, P. S. Quantitative and temporal measurement of dynamic autophagy rates. Autophagy 19, 1164–1183 (2023).

21. Martin, K. R. et al. Computational model for autophagic vesicle dynamics in single cells. Autophagy 9, 74–92 (2013).

22. Kholodenko, B. N. Cell-signalling dynamics in time and space. Nat. Rev. Mol. Cell Biol. 7, 165–176 (2006).

23. Di-Bella, J. P., Colman-Lerner, A. & Ventura, A. C. Properties of cell signaling pathways and gene expression systems operating far from steady-state. Sci. Rep. 8, 17035 (2018).

24. Binda, M. et al. The Vam6 GEF controls TORC1 by activating the EGO complex. Mol. Cell 35, 563–573 (2009).

25. Ostrowicz, C. W. et al. Defined subunit arrangement and rab interactions are required for functionality of the HOPS tethering complex. Traffic 11, 1334–1346 (2010).

26. Chen, Y. et al. A Vps21 endocytic module regulates autophagy. Mol. Biol. Cell 25, 3166–3177 (2014).

27. Caglar, M. U., Teufel, A. I. & Wilke, C. O. Sicegar: R package for sigmoidal and double-sigmoidal curve fitting. PeerJ 6, e4251 (2018).

28. Wilson, D. J. The harmonic mean -value for combining dependent tests. Proc. Natl. Acad. Sci. U. S. A. 116, 1195–1200 (2019).

29. Costanzo, M. et al. A global genetic interaction network maps a wiring diagram of cellular function. Science 353, (2016).

30. Oughtred, R., et al. BioGRID: A Resource for Studying Biological Interactions in Yeast. Cold Spring Harb. Protoc. 2016, db.top080754 (2016).

31. Szklarczyk, D. et al. STRING v10: protein-protein interaction networks, integrated over the tree of life. Nucleic Acids Res. 43, D447–52 (2015).

32. Hao, B. & Kovács, I. A. A positive statistical benchmark to assess network agreement. Nat. Commun. 14, 2988 (2023).

33. Huang, J. K. et al. Systematic Evaluation of Molecular Networks for Discovery of Disease Genes. Cell Syst 6, 484–495.e5 (2018).

34. Baryshnikova, A. Systematic Functional Annotation and Visualization of Biological Networks. Cell Syst 2, 412–421 (2016).

35. Meldal, B. H. M. et al. Analysing the yeast complexome-the Complex Portal rising to the challenge. Nucleic Acids Res. 49, 3156–3167 (2021).

36. Collins, S. R. et al. Functional dissection of protein complexes involved in yeast chromosome biology using a genetic interaction map. Nature 446, 806–810 (2007).

37. Ulitsky, I., Shlomi, T., Kupiec, M. & Shamir, R. From E-MAPs to module maps: dissecting quantitative genetic interactions using physical interactions. Mol. Syst. Biol. 4, 209 (2008).

38. Schuldiner, M. et al. Exploration of the function and organization of the yeast early secretory pathway through an epistatic miniarray profile. Cell 123, 507–519 (2005).

39. Yu, L. et al. Termination of autophagy and reformation of lysosomes regulated by mTOR. Nature 465, 942–946 (2010).

40. Takeda, E. et al. Vacuole-mediated selective regulation of TORC1-Sch9 signaling following oxidative stress. Mol. Biol. Cell 29, 510–522 (2018).

41. Chan, T. F., Bertram, P. G., Ai, W. & Zheng, X. F. Regulation of APG14 expression by the GATA-type transcription factor Gln3p. J. Biol. Chem. 276, 6463–6467 (2001).

42. Pilauri, V., Bewley, M., Diep, C. & Hopper, J. Gal80 dimerization and the yeast GAL gene switch. Genetics 169, 1903–1914 (2005).

43. Albert, B. et al. A Molecular Titration System Coordinates Ribosomal Protein Gene Transcription with Ribosomal RNA Synthesis. Mol. Cell 64, 720–733 (2016).

44. Butow, R. A. & Avadhani, N. G. Mitochondrial signaling: the retrograde response. Mol. Cell 14, 1–15 (2004).

45. MacDonald, C. & Piper, R. C. Genetic dissection of early endosomal recycling highlights a TORC1-independent role for Rag GTPases. J. Cell Biol. 216, 3275–3290 (2017).

46. Hatakeyama, R. et al. Spatially Distinct Pools of TORC1 Balance Protein Homeostasis. Mol. Cell 73, 325–338.e8 (2019).

47. Ma, J. et al. Using deep learning to model the hierarchical structure and function of a cell. Nat. Methods 15, 290–298 (2018).

48. Culley, C., Vijayakumar, S., Zampieri, G. & Angione, C. A mechanism-aware and multiomic machine-learning pipeline characterizes yeast cell growth. Proc. Natl. Acad. Sci. U. S. A. 117, 18869–18879 (2020).

49. Kemmeren, P. et al. Large-scale genetic perturbations reveal regulatory networks and an abundance of gene-specific repressors. Cell 157, 740–752 (2014).

50. Messner, C. B. et al. The proteomic landscape of genome-wide genetic perturbations. Cell 186, 2018–2034.e21 (2023).

51. Mülleder, M. et al. Functional Metabolomics Describes the Yeast Biosynthetic Regulome. Cell 167, 553–565.e12 (2016).

52. Hackett, S. R. et al. Learning causal networks using inducible transcription factors and transcriptome-wide time series. Mol. Syst. Biol. 16, e9174 (2020).

53. Natarajan, K. et al. Transcriptional profiling shows that Gcn4p is a master regulator of gene expression during amino acid starvation in yeast. Mol. Cell. Biol. 21, 4347–4368 (2001).

54. Papinski, D. et al. Early steps in autophagy depend on direct phosphorylation of Atg9 by the Atg1 kinase. Mol Cell 53, 471–483 (2014).

55. Schreiber, A. et al. Multilayered regulation of autophagy by the Atg1 kinase orchestrates spatial and temporal control of autophagosome formation. Mol. Cell 81, 5066–5081.e10 (2021).

56. Jazwinski, S. M. & Kriete, A. The Yeast Retrograde Response as a Model of Intracellular Signaling of Mitochondrial Dysfunction. Front. Physiol. 3, 26575 (2012).

57. Mattiazzi Usaj, M., et al. Systematic genetics and single-cell imaging reveal widespread morphological pleiotropy and cell-to-cell variability. Mol. Syst. Biol. 16, e9243 (2020).

58. Nair, U. et al. SNARE proteins are required for macroautophagy. Cell 146, 290–302 (2011).

59. van der Vaart, A., Griffith, J. & Reggiori, F. Exit from the Golgi is required for the expansion of the autophagosomal phagophore in yeast Saccharomyces cerevisiae. Mol Biol Cell 21, 2270–2284 (2010).

60. Bruch, A., Laguna, T., Butter, F., Schaffrath, R. & Klassen, R. Misactivation of multiple starvation responses in yeast by loss of tRNA modifications. Nucleic Acids Res 48, 7307–7320 (2020).

61. Buchan, J. R., Kolaitis, R.-M., Taylor, J. P. & Parker, R. Eukaryotic stress granules are cleared by autophagy and Cdc48/VCP function. Cell 153, 1461–1474 (2013).

62. Huang, H. et al. Bulk RNA degradation by nitrogen starvation-induced autophagy in yeast. EMBO J 34, 154–168 (2015).

## References

63. Baryshnikova, A. et al. Quantitative analysis of fitness and genetic interactions in yeast on a genome scale. Nat. Methods 7, 1017–1024 (2010).

64. Klionsky, D. J. et al. Guidelines for the use and interpretation of assays for monitoring autophagy (4th edition). Autophagy 17, 1–382 (2021).

65. Janke, C. et al. A versatile toolbox for PCR-based tagging of yeast genes: new fluorescent proteins, more markers and promoter substitution cassettes. Yeast 21, 947–962 (2004).

66. Shaner, N. C. et al. A bright monomeric green fluorescent protein derived from Branchiostoma lanceolatum. Nat. Methods 10, 407–409 (2013).

67. Kuzmin, E. et al. Systematic analysis of complex genetic interactions. Science 360, (2018).

68. Tong, A. H. et al. Systematic genetic analysis with ordered arrays of yeast deletion mutants. Science 294, 2364–2368 (2001).

69. Chollet, F. & j. Allaire, J. Deep Learning with R. (Pearson Professional, 2018).

70. McInnes, L., Healy, J. & Melville, J. UMAP: Uniform Manifold Approximation and Projection for Dimension Reduction. arXiv [stat.ML] (2018).

71. Sainburg, T., McInnes, L. & Gentner, T. Q. Parametric UMAP: learning embeddings with deep neural networks for representation and semi-supervised learning. arXiv. Preprint] arXiv2009.

72. Loader, C. & Liaw, M. A. Package ‘locfit’. https://mirror.linux.duke.edu/cran/web/packages/locfit/locfit.pdf.

73. Langfelder, P., Zhang, B. & Horvath, S. dynamicTreeCut: Methods for detection of clusters in hierarchical clustering dendrograms. R package version.

74. Darsow, T., Rieder, S. E. & Emr, S. D. A multispecificity syntaxin homologue, Vam3p, essential for autophagic and biosynthetic protein transport to the vacuole. J. Cell Biol. 138, 517–529 (1997).

75. Sato, T. K., Darsow, T. & Emr, S. D. Vam7p, a SNAP-25-like molecule, and Vam3p, a syntaxin homolog, function together in yeast vacuolar protein trafficking. Mol. Cell. Biol. 18, 5308–5319 (1998).

76. Wang, C.-W., Stromhaug, P. E., Shima, J. & Klionsky, D. J. The Ccz1-Mon1 protein complex is required for the late step of multiple vacuole delivery pathways. J. Biol. Chem. 277, 47917–47927 (2002).

77. Yang, S. & Rosenwald, A. A High Copy Suppressor Screen for Autophagy Defects in Δ and Δ Strains. G3 7, 333–341 (2017).

78. Ohashi, Y. & Munro, S. Membrane delivery to the yeast autophagosome from the Golgi-endosomal system. Mol. Biol. Cell 21, 3998–4008 (2010).

79. Arlt, H. et al. The dynamin Vps1 mediates Atg9 transport to the sites of autophagosome formation. J. Biol. Chem. 299, 104712 (2023).

80. Shimobayashi, M., Takematsu, H., Eiho, K., Yamane, Y. & Kozutsumi, Y. Identification of Ypk1 as a novel selective substrate for nitrogen starvation-triggered proteolysis requiring autophagy system and endosomal sorting complex required for transport (ESCRT) machinery components. J. Biol. Chem. 285, 36984–36994 (2010).

81. Kato, M. & Wickner, W. Vam10p defines a Sec18p-independent step of priming that allows yeast vacuole tethering. Proc. Natl. Acad. Sci. U. S. A. 100, 6398–6403 (2003).

82. Fisk, D. G. et al. Saccharomyces cerevisiae S288C genome annotation: a working hypothesis. Yeast 23, 857–865 (2006).

83. Yu, G., Wang, L.-G., Han, Y. & He, Q.-Y. clusterProfiler: an R package for comparing biological themes among gene clusters. OMICS 16, 284–287 (2012).

84. Shannon, P. et al. Cytoscape: a software environment for integrated models of biomolecular interaction networks. Genome Res. 13, 2498–2504 (2003).

85. Bader, G. D. & Hogue, C. W. V. An automated method for finding molecular complexes in large protein interaction networks. BMC Bioinformatics 4, 2 (2003).

86. Cherry, J. M. et al. Saccharomyces Genome Database: the genomics resource of budding yeast. Nucleic Acids Res. 40, D700–5 (2012).

87. Chen, T. et al. Xgboost: extreme gradient boosting. R package version 0. 4-2 1, 1–4 (2015).

88. Duong, T. ks: Kernel Density Estimation and Kernel Discriminant Analysis for Multivariate Data in R. J. Stat. Softw. 21, 1–16 (2007).

89. Väremo, L., Nielsen, J. & Nookaew, I. Enriching the gene set analysis of genome-wide data by incorporating directionality of gene expression and combining statistical hypotheses and methods. Nucleic Acids Res. 41, 4378–4391 (2013).

90. Brachmann, C. B. et al. Designer deletion strains derived from Saccharomyces cerevisiae S288C: a useful set of strains and plasmids for PCR-mediated gene disruption and other applications. Yeast 14, 115–132 (1998).

91. Guthrie, C. & Fink, G. R. Guide to Yeast Genetics and Molecular and Cell Biology: Guide to Yeast Genetics and Molecular Biology. (Gulf Professional Publishing, 2004).

92. Gietz, R. D. & Woods, R. A. Transformation of yeast by lithium acetate/single-stranded carrier DNA/polyethylene glycol method. Methods Enzymol. 350, 87–96 (2002).

93. Araki, Y., Kira, S. & Noda, T. Quantitative Assay of Macroautophagy Using Pho8△60 Assay and GFP-Cleavage Assay in Yeast. Methods Enzymol. 588, 307–321 (2017).

94. Noda, T. & Klionsky, D. J. The quantitative Pho8Delta60 assay of nonspecific autophagy. Methods Enzymol. 451, 33–42 (2008).

95. Suzuki, K. et al. The pre-autophagosomal structure organized by concerted functions of APG genes is essential for autophagosome formation. EMBO J. 20, 5971–5981 (2001).

96. Cheong, H. & Klionsky, D. J. Biochemical methods to monitor autophagy-related processes in yeast. Methods Enzymol. 451, 1–26 (2008).

97. Torggler, R., Papinski, D. & Kraft, C. Assays to Monitor Autophagy in Saccharomyces cerevisiae. Cells 6, (2017).

98. Chica, N., Portantier, M., Nyquist-Andersen, M., Espada-Burriel, S. & Lopez-Aviles, S. Uncoupling of Mitosis and Cytokinesis Upon a Prolonged Arrest in Metaphase Is Influenced by Protein Phosphatases and Mitotic Transcription in Fission Yeast. Front Cell Dev Biol 10, 876810 (2022).

99. Foiani, M., Marini, F., Gamba, D., Lucchini, G. & Plevani, P. The B subunit of the DNA polymerase alpha-primase complex in Saccharomyces cerevisiae executes an essential function at the initial stage of DNA replication. Mol. Cell. Biol. 14, 923–933 (1994).

100. Mitchell, J. K., Fonzi, W. A., Wilkerson, J. & Opheim, D. J. A particulate form of alkaline phosphatase in the yeast, Saccharomyces cerevisiae. Biochim. Biophys. Acta 657, 482–494 (1981).

101. Zhang, Y., Jenkins, D. F., Manimaran, S. & Johnson, W. E. Alternative empirical Bayes models for adjusting for batch effects in genomic studies. BMC Bioinformatics 19, 262 (2018).

102. Moritz, S. & Bartz-Beielstein, T. ImputeTS: Time series missing value imputation in R. R J. 9, 207 (2017).

103. Liu, Y., Just, A. & Mayer, M. SHAPforxgboost: SHAP plots for ‘XGBoost’. R package version 0.1. 0.

